# A CAR-T Cell-Based Strategy for Eliminating Pathogenic Microglia in ALS

**DOI:** 10.64898/2025.12.29.696307

**Authors:** Maria Elena Cicardi, Daniela Tejada-Martinez, Shashirekha S. Markandaiah, Karthik Krishnamurthy, Chiara Scopa, Brij Oza, Ivette Martorell Serra, Andrea Feci, Edoardo Manca, Adam E. Snook, Aaron R. Haeusler, Piera Pasinelli, Davide Trotti

## Abstract

Neurodegenerative diseases are defined by the propagation of neuroinflammation, driven in part by disease-associated microglia (DAM) that amplify inflammatory signaling and hasten neurodegeneration. Strategies to selectively eliminate DAM to attenuate disease progression remain elusive. Using existing datasets in combination with multiplexed immunofluorescence analysis of post-mortem ALS tissues, we identified the urokinase-type plasminogen activator receptor (uPAR) as a novel surface marker of DAM. uPAR protein is markedly elevated in IBA1⁺/CD68⁺ microglia within ALS-affected regions of both sporadic and familial cases, with negligible expression in unaffected areas or control tissues. These uPAR-high microglia drive neurite retraction in iPSC-derived neurons. To target these cells, we engineered 3^rd^-generation CAR-T cells expressing an anti-uPAR single-chain variable fragment, enabling specific recognition and elimination of uPAR-expressing microglia. Target specificity was assessed in human microglia C20 cells driven into a pathogenic state by poly(IC) or IFNγ stimulation, which resulted in robust surface uPAR expression alongside phagocytic (CD68⁺) and antigen-presenting (CD80⁺) markers. uPAR-CAR-T cells induced antigen-dependent cytolysis of uPAR-high microglia, reducing their viability by over 80% while sparing resting microglia and neurons in a mixed culture system. These results position uPAR-directed CAR-T cells as a viable immunotherapeutic approach to selectively disrupt disease-amplifying microglial subsets and modify the trajectories of neuroinflammatory diseases.

**One Sentence Summary:** CAR-T cells targeting uPAR selectively ablate pathogenic microglia while sparing neurons, enabling precision immunotherapy for ALS.

## INTRODUCTION

Therapeutic success in the cancer field has been driven, in part, by innovative approaches to eliminate cancer cells and prevent or cure metastatic disease. More recently, the specificity of cancer cell elimination has been enhanced by the development of chimeric antigen receptor (CAR)-T cell therapy, which harnesses the intrinsic cytotoxicity of T cells genetically reprogrammed to recognize specific surface antigens in a major histocompatibility complex-independent manner^1^. These products were initially developed for refractory B-cell malignancies, and several CAR-T agents are now approved for hematological cancers^1^. Given the accumulating evidence of their clinical success and the fact that T cells can traverse the blood-brain barrier even in homeostatic conditions, CAR-T platforms have also been proposed for diseases of the CNS^2^. Early-phase trials in glioblastoma and progressive multiple sclerosis demonstrate that engineered T cells can infiltrate the CNS and remain pharmacodynamically active^3^. Yet, durable efficacy against solid tumors or diffuse neuroinflammation remains unknown. Nevertheless, these findings suggest that cell-based immunotherapy approaches being developed to target cancerous cells in the CNS could be more broadly applied to eliminate other deleterious cell types in the CNS.

ALS and its clinical-pathological overlap with frontotemporal dementia (FTD) exemplify neurodegenerative conditions in which immunotherapeutic approaches for selective elimination of disease-spreading cells could be transformative^4^. Post-mortem analyses of spinal cord and frontal cortex tissue consistently reveal intense neuroinflammation, marked by gliosis^5^. Classically defined reactive astrocytes and microglia upregulate complement components (C1q, C3)^6,7^, intermediate filaments (GFAP)^8^, phagolysosomal proteins (CD68)^9^, and secrete pro-inflammatory cytokines such as IL-1α/β, IL-6, and TNF-α, all of which correlate with rapid clinical decline^10,11^. Observations such as these, as well as recent single-cell transcriptomic studies, support a model in which non-cell-autonomous toxicity from glia accelerates neuronal loss^12^. Among glial populations, microglia occupy a unique immunological niche. They continuously survey the parenchyma, sculpt synapses, and orchestrate leukocyte recruitment under injury conditions^13,14^. The diverse functions of microglia are reflected in the vast heterogeneity at the multi-omics level^15^, which all depend on the context^16^. In neurodegenerative diseases, including ALS, microglia transition into disease-associated states marked by excessive synapse stripping, heightened MHC-II antigen presentation, and the release of pro-inflammatory cytokines and reactive oxygen species^17–20^. Selectively eliminating or reprogramming these disease-associated pathogenic microglia, rather than broadly suppressing neuroinflammation, may disrupt self-perpetuating injury while preserving the protective functions of homeostatic microglia.

Here, we provide a foundational evaluation of CAR-T cells as a tractable immunotherapeutic strategy for eliminating disease-associated microglia (DAM) shared across ALS etiologies. By integrating insights from preclinical studies examining tractable surface antigens in preclinical cancer models^21^ with emerging transcriptomic profiles in ALS patients and models^22,23^, we converged on a rationale candidate target: the urokinase-type plasminogen activator receptor uPAR/CD87^24^. Using multiplexed immunofluorescence imaging, we first demonstrate that the uPAR protein has limited expression levels in the CNS but is elevated in post-mortem tissues from both sporadic and genetic ALS cases, where uPAR^+^ cells frequently colocalize with CD68/IBA1-positive DAM. Second, we then modeled inflammatory up-regulation of uPAR in human C20 microglia, defined its co-expression with phagocytic (CD68) and antigen-presentation (CD80) markers, and demonstrated their consequent neurotoxicity in iPSC-derived neuron-glia co-cultures. Finally, we engineered a third-generation human uPAR-CAR construct and demonstrated antigen-dependent cytolysis of uPAR^+^ microglia while sparing surface uPAR-negative neurons. Collectively, these results identify uPAR as a viable surface marker of disease-amplifying microglia, providing a therapeutic framework in which CAR-T cells targeting these cell populations could recalibrate the neuroinflammatory environment and slow disease progression in ALS, with potential applicability to other microglia-driven neurodegenerative disorders.

## RESULTS

### Region-Specific uPAR Up-Regulation Identifies a Therapeutically Actionable Microglial Subset in ALS

We postulated that, rather than broadly suppressing neuroinflammation, a more precise strategy would be to eliminate only the disease-associated microglial subset that drives neurodegeneration. To this end, we sought a cell-surface immunotherapy target with clear translational potential, guided by three criteria: (1) it must be a plasma-membrane antigen amenable to CAR-T cell recognition; (2) it should exhibit minimal surface expression on microglia under non-disease conditions; and (3) it must be robustly induced in disease.

By integrating emerging CAR-T cell targets validated in preclinical cancer models^24,25^ with transcriptomic signatures of ALS disease-associated tissues^26–31^, a single convergent antigen was rationally selected: the urokinase plasminogen activator receptor (uPAR; encoded by *PLAUR*). uPAR is a glycosyl-phosphatidylinositol-anchored receptor for the zymogen uPA that regulates extracellular matrix remodeling and cell migration. Outside of wound healing and embryogenesis, uPAR expression is negligible in healthy adult tissues but is strongly induced in cancer, fibrotic organs, and senescent cell populations^32^. In the CNS, uPAR is virtually undetectable under homeostatic conditions, yet it becomes conditionally expressed on myeloid-lineage cells in states of chronic tissue stress, neuroinflammation, and senescence^33,34^. Importantly, murine studies have demonstrated that uPAR-targeted CAR-T cells can ablate senescent hepatic stellate cells and ameliorate fibrosis without on-target/off-tumor toxicity^25,35,36^, further supporting their translational tractability.

Our analysis of publicly available transcriptomic datasets (GSE137810; GSE219281) revealed that *PLAUR* transcripts are enriched in cervical spinal cord samples in bulk ALS tissue and that single-nucleus UMAP projections show elevated *PLAUR* signals localized to a significant fraction of microglia in both frontal and motor cortices (**Supplementary Fig. 1A,B**). Consistent with these human datasets, two ALS genetic mouse models (SOD1^G93A^ and TDP^ΔNLS^) exhibited minimal uPAR expression before disease onset, followed by a rapid and robust induction in disease-affected areas over time (**Supplementary Fig. 2C, D**). This disease-restricted transcriptomic expression profile, combined with preclinical evidence demonstrating the safety and efficacy of uPAR-targeted CAR-T cells, positions uPAR as an attractive antigen for cell-directed immunotherapies designed to eliminate pathogenic microglia while sparing bystander cells.

Based on these transcriptomic findings, we next asked whether the increase in uPAR transcript extends to the protein level. Consistent with the transcriptomic results, western blot analysis of mid-motor cortex lysates from seven C9-ALS/FTD cases, five sporadic ALS cases, and eight controls revealed significantly elevated uPAR expression in ALS (**Figure 1A, B**). The occipital cortex, generally regarded as relatively spared in ALS/FTD compared with the motor cortex, displayed uPAR protein levels comparable to those observed in the motor cortex. Western blotting was employed to quantify uPAR protein levels; however, spatial distribution remains unresolved. This finding suggests that regional sparing may not extend to uPAR expression, although the precise anatomical extent remains indeterminate (**Supplementary Figure 2A, B**).

**Fig. 1.**
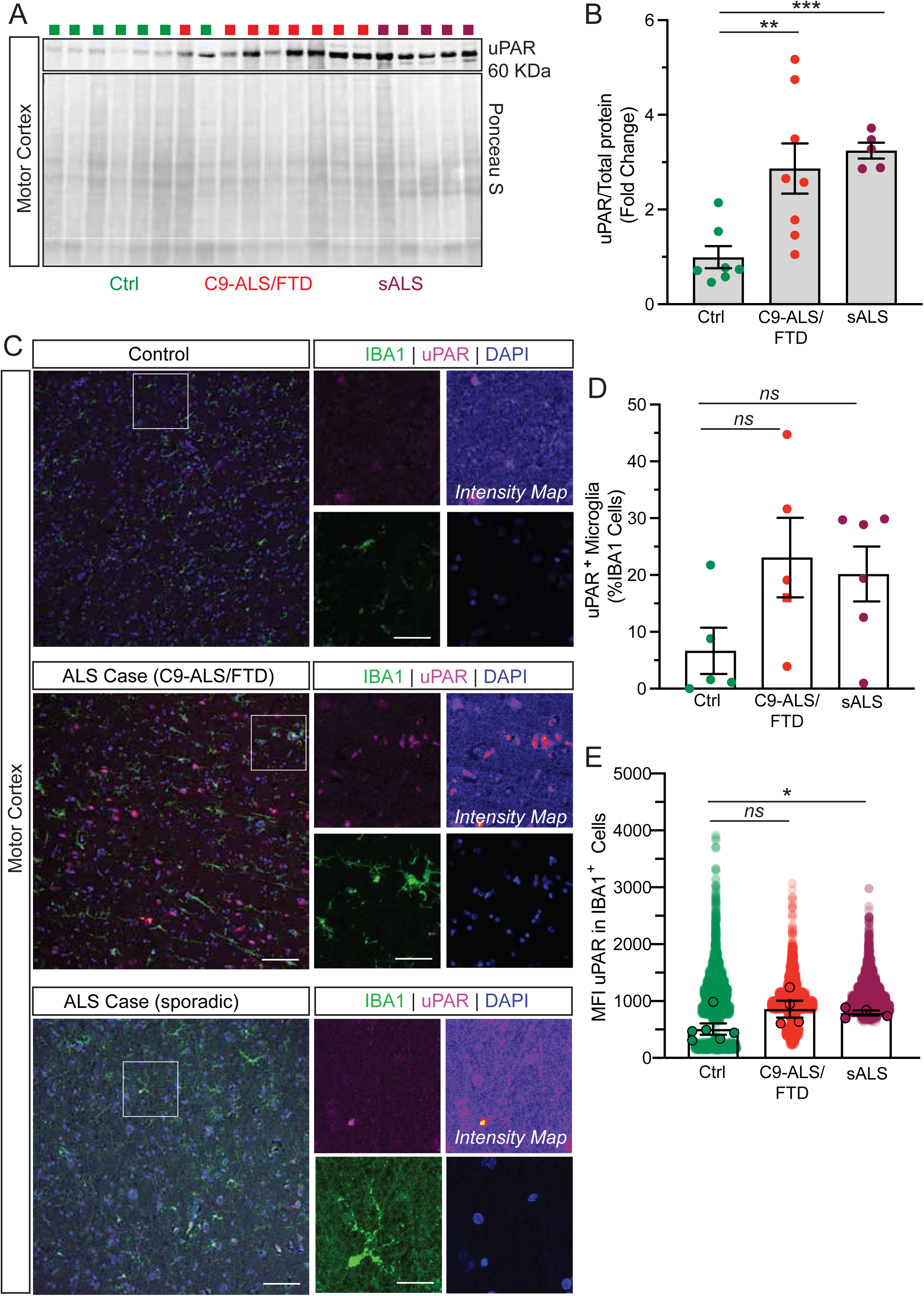
uPAR is selectively up-regulated in ALS motor cortex. **(A)** Western blot of total protein lysates prepared from the mid motor cortex of neurologically normal controls (Ctrl, green), C9orf72-linked ALS/FTD patients (C9-ALS, red), and sporadic ALS patients (sALS, purple). Ponceau-S staining of the same membrane is shown as an internal loading/transfer control. Each lane corresponds to an individual donor (see Table 2 for demographics and post-mortem interval). **(B)** Quantification of uPAR signal normalized to total Ponceau-S, and expressed as fold-change over the control group mean. Bars show mean ± S.E.M.; points represent individual brains (Ctrl n = 8, C9-ALS/FTD n = 7, sALS n = 6). One-way ANOVA with Tukey post-hoc: (*p < 0.05, **p < 0.01). **(C)** Representative sections of mid motor cortex were stained for microglia (Iba1, Alexa 488, green), uPAR (CD87, Alexa 647, magenta) and nuclei (DAPI, blue). Left panels: low-magnification fields (495 µm × 495 µm, 20× objective). Right panels: 63× optical zooms (digital zoom = 2) of the boxed regions together with a rainbow-coded “Intensity Map” highlighting relative uPAR signal. Scale bars: overview = 100 µm; insets = 20 µm. **(D)** Percentage of IBA1⁺ cells that were also uPAR⁺ was calculated from three non-overlapping regions of interest per donor (total cells analyzed: Ctrl 3,042; C9-ALS 3433; sALS 3,161). Bars depict mean ± S.E.M.; symbols are biological replicates (Ctrl n = 8, C9-ALS n = 7, sALS n = 6). One-way ANOVA with Dunnett post-hoc versus control: p = 0.06 (Ctrl vs C9-ALS/FTD), p = 0.08 (Ctrl vs sALS). **(E)** Plot shows the distribution of uPAR mean fluorescence intensity (MFI) within individual IBA1⁺ microglia. One-way ANOVA with Tukey post-hoc: (*p < 0.05, **p < 0.01).

We also detected increased uPAR expression in spinal cord tissue from mutant SOD1 and sporadic ALS patients. In contrast, spinal cords from C9-ALS/FTD cases did not exhibit elevated uPAR levels compared to controls (**Supplementary Figure 2C, D, Table 1, 2**).

Given prior reports of microglia-specific upregulation of uPAR^33,37^, we then asked whether microglia display elevated uPAR expression in ALS. We therefore performed immunofluorescence (IF) staining on mid-motor cortex sections from C9-ALS/FTD and sporadic ALS cases using IBA1 and GFAP to mark microglia and astrocytes, respectively. IF analysis revealed a trend to elevated uPAR expression in microglia in C9-ALS/FTD cases (**Figure 1C, D**), while uPAR^+^ astrocytes remain essentially unchanged (**Supplementary Figure 2E, F**). Further, quantitative IF analysis revealed a trend toward increased uPAR^+^ microglia in both C9 and sporadic ALS cases, with approximately 25% of IBA^+^ microglia expressing uPAR compared to fewer than 10% in control samples (**Figure 1C, D**). Although this difference did not reach statistical significance, the mean fluorescence intensity (MFI) of uPAR per IBA^+^ microglial cell was significantly elevated in ALS tissues relative to controls (**Figure 1C, E; Supplementary Figure 2E, G**). In contrast, astrocytic uPAR expression remained constant across all groups. These findings suggest that ALS-associated pathological mechanisms lead to a selective increase in uPAR expression in a subset of microglia, offering a potentially tractable opportunity for targeted immunotherapeutic interventions.

### Identification of a uPAR-high (uPAR^hi^) Microglial Subpopulation in ALS Cases via Multiplexed Immunofluorescence

To further identify the specific cell types and subtypes associated with uPAR expression in ALS, we performed CODEX-based multiplex imaging microscopy using a curated 23-antibody multiplex panel (**Table 3**) to assemble a comprehensive single-cell spatial proteomic map of the human CNS. This dataset profiles four distinct tissue regions - motor cortex (MC), spinal cord (SC), frontal cortex (FC), and occipital cortex (OC) from a cohort including a non-pathological control, a stroke control, and different ALS genotypes (sALS, C9_ALS, and SOD1_ALS) (**Supplementary Figure 3 A-D**). Unsupervised clustering of single-cell protein expression data identified eight major cell lineages: neurons, oligodendrocytes, endothelial cells, vascular cells, ependymal cells, astrocytes, M2 macrophages, and microglia. The UMAP projections show the clear segregation of these lineages, and cell identity was validated by the distinct and canonical expression patterns of cell-specific markers (**Figure 2A-D**). Importantly, cell-type proportions align with those reported in spatial transcriptomic datasets from diseased and non-diseased tissues that appreciate white- and gray-matter contributions (**Table 4**).

**Fig. 2.**
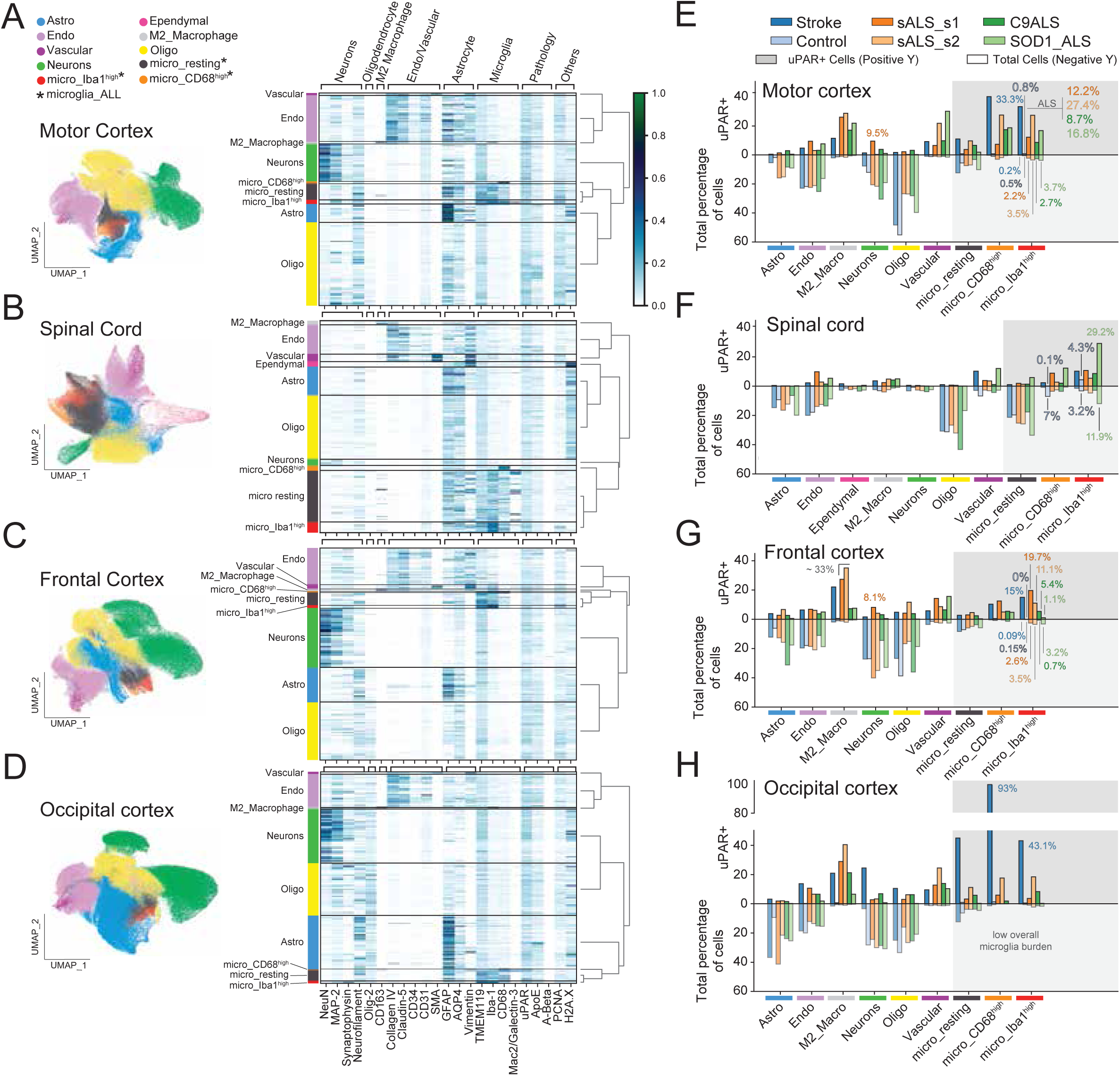
Single-cell profiling and uPAR^+^ distribution across four CNS regions. For each region: **(A)** Motor Cortex, **(B)** Spinal Cord, **(C)** Frontal Cortex, **(D)** Occipital Cortex, the left panel displays a UMAP projection of all single cells, colored by annotated cell type. The right panel shows a single-cell heatmap of the scaled expression for the final 23-plex antibody panel across the cell types. **(E-H)** Divergent bar plots for each corresponding region quantify cellular composition and uPAR+ prevalence. The negative y-axis shows the proportion of each cell type across samples. The positive y-axis shows the proportion of uPAR+ cells within each of those cell types.

In this multiplex spatial imaging analysis, the MC and SC emerged as hallmarks of uPAR^+^ microglial activation in ALS. Analysis of the MC revealed uPAR^+^ prevalence in IBA1^+^ microglia across ALS genotypes, ranging from 8.7% in C9_ALS to 27.4% in sALS, in contrast to the control (0.8%) (**Figure 2E**). Similar patterns of uPAR positivity were observed in the microglia CD68^+^ cell population across ALS genotypes and the stroke case. Likewise, M2 macrophages consistently showed a high proportion of uPAR^+^ cells across ALS and stroke cases, with the highest levels observed in sALS in the FC (∼ 33.0%) and MC (∼27%) (**Figure 2E,G**). In contrast, the SOD1_ALS case was characterized by a strong microgliosis with uPAR^+^ signature in the spinal cord (**Figure 2F**). The total microglial burden (comprising micro_resting, micro_CD68, and micro_IBA1 populations; see Materials and Methods for inclusion criteria) was highest in the SOD1_ALS case (46.1%), representing a marked enrichment compared to the control (29.9%). This increase was primarily driven by a substantial expansion of IBA1^+^ microglia, which accounted for 11.9% of all cells. Additionally, an inflammatory response explicitly characterized by uPAR^+^ activated microglial states was shared across ALS genotypes. The uPAR^+^ signature was more pronounced in the CD68^+^ microglial state, where SOD1_ALS exhibited a 12.4% uPAR^+^ proportion compared to only 0.1% in the control, a 124-fold increase. A robust uPAR^+^ signature was also present in the IBA1^+^ microglial state, with SOD1_ALS showing a 29.2% uPAR^+^ proportion, compared to 4.3% in the control. Furthermore, IBA1^+^ microglia also showed a high proportion of uPAR^+^ cells in the sALS (∼10.7%), C9_ALS (8.6%), and stroke (10.2%) cases. Interestingly, the lowest level of uPAR^+^ cells was found in the stroke case in the SC, which was not impacted by the stroke. This observation indicates that uPAR may serve as a standard marker of pathogenic microglial activation, independent of ALS genotype. Its robust induction not only in ALS tissues but also in the stroke sample, an acute condition marked by intense neuroinflammation, suggests that uPAR expression marks a broader microglial injury-response program. This positions uPAR as a potential hallmark of disease-associated and disease-amplifying microglia across both chronic neurodegenerative and acute inflammatory CNS insults. The OC exhibited the lowest overall microglial burden in ALS cases (ranging from 3.8% to 4.8%). However, in the stroke case, the OC displayed a unique and intense focal activation, where nearly all (93%) CD68^+^ microglia were uPAR^+^, and a significant 43.1% of IBA1^+^ microglia were also uPAR^+^. This indicates a robust inflammatory upregulation specific to the stroke pathology in this region (**Figure 2H**). On the contrary, the OC was among the least affected areas of uPAR^+^ engagement regions in ALS.

In other cell types, a notable uPAR^+^ signature was also observed in vascular and endothelial cells across ALS genotypes, consistent across tissues. While neurons exhibited very low uPAR^+^ proportions across most tissues and genotypes (typically <0.5%), a specific sALS case (sALS_s1) showed an increase, with uPAR^+^ proportions reaching 9.5% in the MC and 8.1% in the FC (**Figure 2E-G**). This contrasted with control samples, where the neuronal uPAR^+^ signature was minimal (<0.02%).

Analysis of expression profiles of microglial markers revealed that ALS and stroke cases were characterized by higher mean expression of IBA1 and uPAR compared to controls across tissues (**Figure 3A-D**). Spatial analysis confirmed this microglial specificity, demonstrating that uPAR^+^ cells primarily co-localized with microglial cells across tissues (**Figure 3E-G**, **left panels**), with negligible signal observed in neurons, astrocytes, or endothelial cells (**Figure 3E-G**, **right panels**). The highest expression of uPAR in microglial cells was localized to the spinal cord in SOD1_ALS (**Figure 3B**), where these uPAR^+^ microglia form dense aggregates and are distributed across both white matter and towards the ventral horn in the grey matter (**Figure 3F**, **Supplementary Figure 4**). Additionally, a marked specific overexpression of Mac2/Galectin-3 was observed in ALS cases (**Figure 3B**), with the strongest signal in the spinal cord of an SOD1 ALS patient (**Supplementary Figure 5**). The consistent low expression of Mac2/Galectin-3 in control and stroke cases within this region suggests its potential upregulation as a specific feature of ALS pathology (**Figure 3B, Supplementary Figure 5**). The OC displayed the lowest levels of microglial uPAR expression across all conditions (**Figure 3D**), consistent with its overall lower microglial burden.

**Fig. 3.**
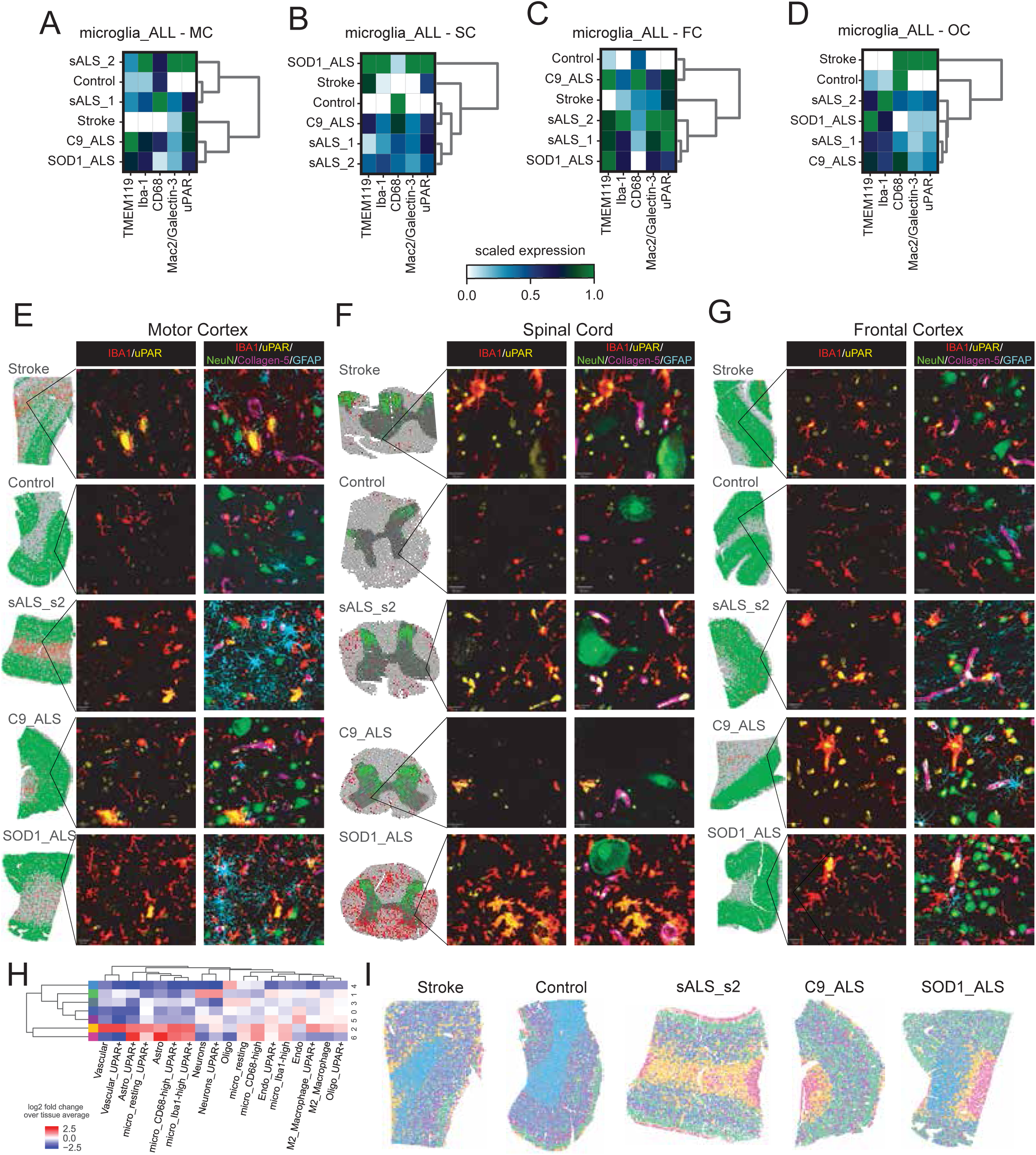
Spatial and disease-specific microglial uPAR expression. **(A-D)** Scaled expression of microglial markers (TMEM119, Iba-1, CD68, Mac2/Galectin-3) and uPAR across disease conditions for the **(A)** MC, **(B)** SC, **(C)** FC, and **(D)** OC. **(E-G)** Spatial distribution of uPAR+ microglia_ALL. For each region, the left panel is a spatial scatter plot showing cell coordinates (Red: uPAR+ microglia; Green: neurons; Grey: other cells) to differentiate grey and white matter. The right panels show representative multiplexed images from QuPath (v0.6.0) and canonical markers for major cell types: the left image shows merged Iba-1 (green) and uPAR (red) channels; the right Iba-1, uPAR, NeuN (neurons), Collagen IV (endothelial cells), and GFAP (astrocytes). **(H)** Motor Cortex Cellular Neighborhood (CN) heatmap with the log2 fold-change enrichment of specific cell types within each CN. The spatial map illustrates the distribution of the identified neighborhoods in space.

To better understand the relationship between uPAR^+^ cells and the surrounding tissue microenvironment, we performed a cellular neighborhood (CN) analysis. These analyses revealed distinct spatial niches for uPAR^+^ cells, which varied by anatomical region (**Supplementary Figure 6-9**). In the MC, uPAR^+^ cells, predominantly uPAR^+^ microglia including CD68^+^ and IBA1^+^ states, and vascular cells, were enriched within a specific CN (yellow, **Figure 3H-I**) localized to the white matter and spatially associated with an astrocyte-enriched CN (pink, **Figure 3H, I, Supplementary Figure 6A-C**). This uPAR^+^ CN had a generally low frequency of spatial proximity (<3%, **Supplementary Figure 6D-I**) to the neuron-enriched CN (green, **Figure 3H,I**). A similar pattern of uPAR^+^ cell localization in white matter was observed in the SC and OC (**Supplementary Figure 7, 9 A-C**). Notably, in the SC, the uPAR^+^ microglia-enriched CN was distributed across both the posterior white matter and the grey matter of the ventral horn (**Supplementary Figure 7C**), proximate to the anatomical location of motor neurons. This pattern was most prominent in the SOD1_ALS case, which exhibited a high aggregation of uPAR^+^ microglia (**Supplementary Figure 4**). Furthermore, in the FC, the CN with the highest log_2_ fold-change enrichment was defined by uPAR^+^ vascular cells and uPAR^+^ M2 macrophages, identifying a primary non-microglial niche for uPAR expression (pink cluster, **Supplementary Figure 8B-C**).

Collectively, our spatial proteomic imaging analysis indicates that the uPAR^+^ cellular phenotype is not uniformly distributed across the ALS CNS, but potentially follows a distinct anatomical, genotypic, and/or clinical pattern. We observed a pronounced spatial gradient, with the lowest uPAR^+^ signal in the OC, approaching the near-zero baseline of controls, and a substantially increased signal in the MC and SC. This pattern was critically defined by the ALS genotypes, with the spinal cord of the SOD1_ALS case showing an increase of microgliosis and a high component of uPAR^+^ across all microglial stages (microglia_ALL). Beyond the microglia_ALL, the uPAR^+^ phenotype was also present in vascular cells but was essentially negligible in neurons. As previously reported by others^38^, Mac2/Galectin-3 was identified as a potential ALS-specific marker, showing marked overexpression in the SC of ALS cases, compared to controls (including stroke). Spatially, uPAR^+^ microglia_ALL localized primarily to white matter in cortical regions but also to the grey matter ventral horn in the SC. Our results from the single-cell proteomic imaging analysis establish the uPAR^+^ cellular landscape and specifically identify microglia as a key cellular target.

### uPAR is Upregulated in Disease-Associated Microglia and Defines Functionally Distinct States

Based on the spatial proteomic imaging results, we next asked whether these phenotypes could be recapitulated *in vitro* and whether the pathophysiological properties of uPAR⁺ microglia could be further evaluated. To accomplish this, we utilized C20 microglial cells derived from a primary human microglia cell line. These cells preserve key microglial markers and exhibit a robust spectrum of responses to pro-inflammatory stimuli, such as IFNγ^37,39^. This heterogeneity makes C20 cells a valuable system for dissecting the molecular and functional features of the disease-associated microglia spectrum and uPAR regulation under defined conditions, as well as a tractable model for testing target engagement and specificity of uPAR-CAR-T cells. To simulate disease-relevant microglial states *in vitro*, we exposed C20 cells to a panel of toll-like receptor (TLR) agonists and pro-inflammatory cytokines known to stimulate innate immune challenges that may be encountered during neurodegeneration. Specifically, cells were treated with Pam3CSK4 (a TLR2/1 agonist), LPS (TLR4), R848 (TLR7/8), and polyIC (TLR3), as well as the cytokines interferon-gamma (IFNγ) and tumor necrosis factor-alpha (TNFα). These stimuli were selected to engage distinct TLR pathways and inflammatory cascades implicated in ALS^40^, thereby enabling a systematic evaluation of their ability to upregulate uPAR and trigger disease-associated phenotypes. Flow cytometry analysis revealed that Pam3CSK4, polyIC, and IFNγ significantly increased both the percentage of uPAR^+^ cells and the surface expression levels of uPAR, as measured by MFI (**Supplementary Figure 10A-C**). The most robust increases were observed with polyIC and IFNγ, which approximately doubled the proportion of uPAR^+^ cells and resulted in a ∼4-fold increase in uPAR surface expression. To evaluate downstream TLR pathway activation, we measured the phosphorylation of NF-κB p65 (also known as ReIA) at serine 536 (pNF-κB S536), a readout that decreases upon canonical NF-κB activation. Except for TNFα, all stimuli significantly reduced pNF-κB S536 levels, suggesting pathway engagement (**Supplementary Figure 10D, E**). This activation was accompanied by visible morphological changes in the C20 cells, including alterations in actin cytoskeleton organization (**Supplementary Figure 10D)**.

Using the two most robust inducers of uPAR expression identified in our stimulus panel, polyIC and IFNγ, we next asked whether classical markers of activated microglia preferentially associate with elevated uPAR levels in vitro (**Figure 4A,B**). To capture stimulus-induced phenotypic heterogeneity in an unbiased manner, we performed t-distributed stochastic neighbor embedding (t-SNE) on flow cytometry data using a multidimensional parameter set that included forward scatter height (FSC-H), side scatter height (SSC-H), uPAR (CD87-PE) intensity, and viability (DRAQ7), thereby integrating cell size, granularity, activation-related surface expression, and cell health into a single low-dimensional representation. Importantly, uPAR was included as one dimension within this feature space and subsequently used as an overlay to visualize its distribution across emergent phenotypic clusters, rather than serving as the sole driver of dimensionality reduction. In this analysis, polyIC- and IFNγ-treated microglia formed discrete clusters that segregated from PBS-treated controls, indicating that these stimuli induce broad, multidimensional phenotypic remodeling beyond changes in uPAR expression alone (**Figure 4C**). Notably, regions of the t-SNE map corresponding to stimulated clusters were enriched for cells with higher uPAR signal, suggesting that uPAR upregulation tracks with specific activation-associated cellular states rather than occurring uniformly across the population. To further contextualize these phenotypic shifts, we examined the expression of two well-characterized markers of microglial activation, CD68 and CD80^41^, in conjunction with uPAR. CD68, a lysosomal glycoprotein associated with phagocytic activity, showed minimal baseline surface expression, with only ∼1.2% of unstimulated C20 cells scoring positive (**Figure 4D,E**). Similarly, CD80, a co-stimulatory molecule involved in antigen presentation and immune activation, was detected in ∼0.7% of cells under basal conditions (**Figure 4F,G**). Following polyIC or IFNγ stimulation, however, the mean fluorescence intensity of CD68, CD80, and uPAR increased significantly, indicating coordinated upregulation of activation-associated markers at the single-cell level (**Figure 4H**). Subsequent co-expression analysis by conventional flow cytometry (**Supplementary Figure 11A**, gating strategy) demonstrated a marked increase in the proportion of CD68⁺/uPAR⁺ and CD80⁺/uPAR⁺ double-positive cells following stimulation (**Figure 4G,L**). These findings indicate that uPAR upregulation preferentially occurs within microglial subsets exhibiting phagocytic and immune-activation signatures, rather than representing a generalized activation marker.

**Fig. 4.**
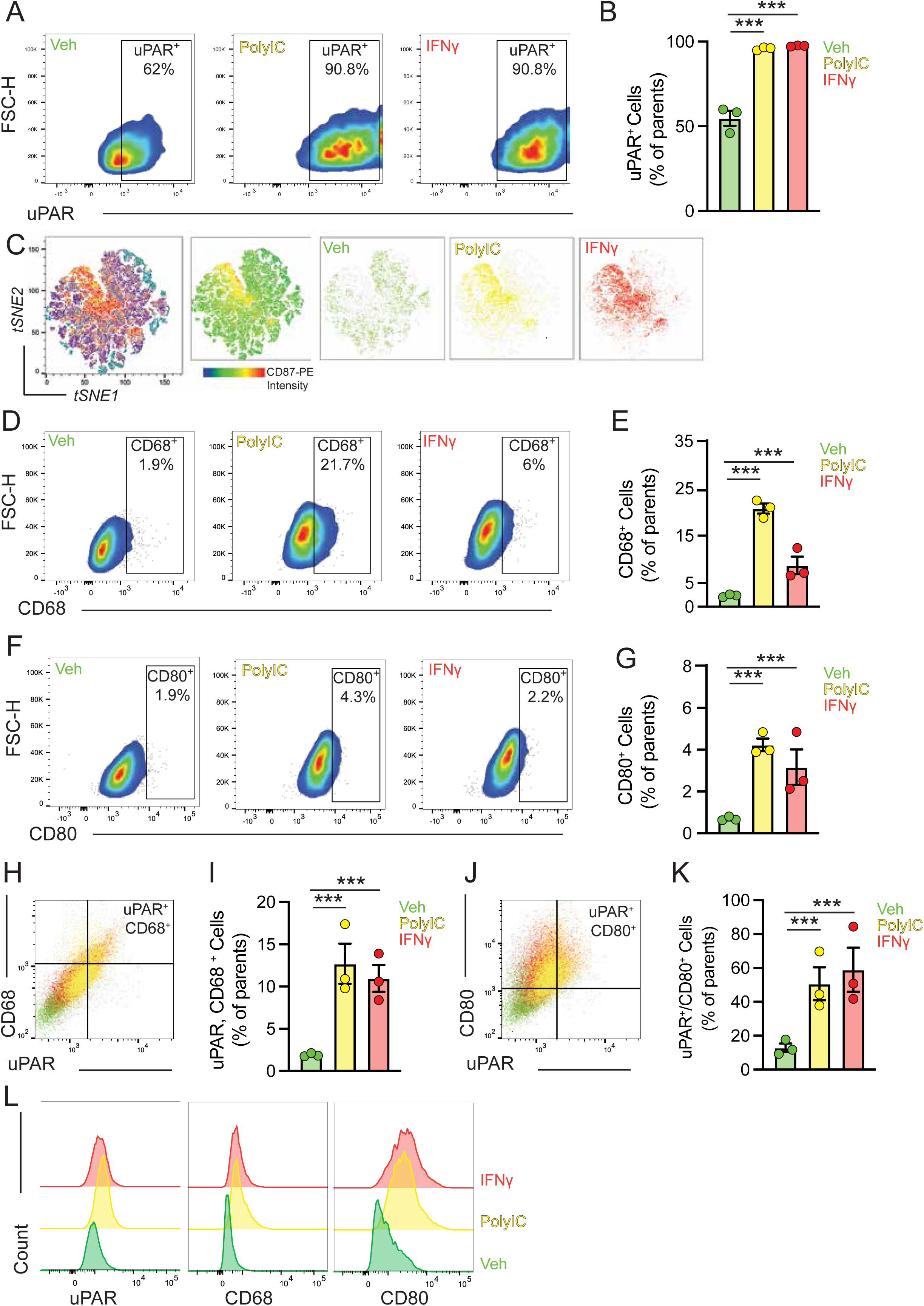
Disease-associated microglia express uPAR at the cell surface. **(A)** Representative flow cytometry gating of C20 microglial cells treated for 48 hours with polyIC and IFN-γ, stained for uPAR. X-axis: uPAR fluorescence intensity; Y-axis: forward scatter (FSC). **(B)** Quantification of the percentage of uPAR-positive C20 cells under the indicated conditions. Data represent the mean ± S.E.M. of three technical replicates, representative of three independent experiments. (*p < 0.05, **p < 0.01, ***p < 0.001). **(C)** UMAP visualization of multiparameter flow cytometry data. A total of 27,000 viable single C-20 cells (nine samples: 3,000 events × three technical replicates per condition) stained for CD87/uPAR-PE, CD68-BV421, and CD80-APC-Cy7 were concatenated and transformed using arcsinh (co-factor = 150). UMAP embedding was performed using the FlowJo plugin (nearest neighbors = 15, minimum distance = 0.3). Left: composite UMAP colored by CD87-PE median intensity (blue → yellow → red). Right: same map showing distribution of cells exposed to vehicle (PBS, green), polyIC (10 μg/mL, yellow), or IFN-γ (20 ng/mL, red). **(D)** Representative flow cytometry gating of C-20 cells treated for 48 hours with polyIC and IFN-γ, stained for CD68. X-axis: CD68 fluorescence intensity; Y-axis: FSC. **(E)** Quantification of CD68-positive C-20 cells in each condition. Data are shown as mean ± S.E.M. of three technical replicates, representative of three independent experiments. (*p < 0.05, **p < 0.01, ***p < 0.001). **(F)** Representative flow cytometry gating of C-20 cells treated for 48 hours with polyIC and IFN-γ, stained for CD80. X-axis: CD80 fluorescence intensity; Y-axis: FSC. **(G)** Quantification of CD80-positive C-20 cells in each condition. Mean ± S.E.M. of three technical replicates from three independent experiments. (*p < 0.05, **p < 0.01, ***p < 0.001). **(H)** Overlaid histograms showing fluorescence intensity distributions of uPAR, CD68, and CD80 under the three treatment conditions: PBS (green), polyIC (yellow), and IFN-γ (red). Rightward shifts reflect increased marker surface expression following stimulation. Statistical analysis: one-way ANOVA with Tukey’s post-hoc test; n = 9 (three independent experiments with three replicates per condition). ***p < 0.001. **(I)** Representative density plot showing co-expression of uPAR (X-axis) and CD68 (Y-axis) after polyIC stimulation. Q2 quadrant indicates double-positive cells. **(J)** Quantification of uPAR⁺/CD68⁺ double-positive microglia (% of viable singlets). Mean ± S.E.M.; n = 9. ***p < 0.001. **(K)** Representative plot of uPAR (X-axis) versus CD80 (Y-axis) expression following 48-hour IFN-γ stimulation. **(L)** Quantification of uPAR⁺/CD80⁺ double-positive cells. Statistics as in panel (I).

Taken together, these data demonstrate that defined inflammatory stimuli drive uPAR expression within distinct, multidimensionally defined disease-associated microglial states. The concordance between unsupervised phenotypic clustering, classical activation markers, and uPAR expression reinforces uPAR as a surface marker enriched in pathogenic microglial subsets and corroborates our multiplex imaging analyses showing selective uPAR elevation in microglia in ALS tissues (**Figure 3**).

### uPAR-Expressing Microglia Induce Neurite Degeneration in Cortical-Like Neurons

We next sought to determine whether the enrichment of uPAR-expressing microglia was intrinsically toxic and impacted neuronal integrity in microglia-neuronal co-culture systems. To accomplish this, we first stimulated the C20 cells with polyIC or IFNγ to induce a uPAR^+^ disease-like state, as in **Figure 4**. We then evaluated the impact of these microglial cells, both with and without stimulation, using human iPSC-derived, cortical-like i^3^Neurons obtained from three control donors and three C9orf72 ALS/FTD patients^42^. Neurons were co-cultured with either PBS-treated or polyIC or IFNγ-stimulated C20 microglia, and neurite integrity was assessed after 24 and 48 hours through immunostaining for the neuronal marker Map2 (**Figure 5A, Supplementary Figure 12A**). Co-culture of neurons with C20 microglia, regardless of their activation state, resulted in significant neurite retraction compared to neurons cultured alone, highlighting a baseline level of microglia-mediated neuronal distress (**Figure 5B,C**). We conducted high-content imaging analyses to characterize further and quantify the extent of neurite degeneration and the distribution of neurite lengths. Data were organized into bins of 10 µm increments, revealing that neurons co-cultured with polyIC- or IFNγ-stimulated microglia exhibited a significant shift to shorter neurites (length between 10-50 µm) compared to neurons co-cultured with unstimulated, PBS-treated microglia (**Supplementary Figure 12B-G**). These observations were consistent across all i^3^Neuronal lines examined and suggested that reactive/disease-associated-like microglia actively promote neurite retraction, a pathological neurotoxic event that typically precedes neuronal death.

**Fig. 5.**
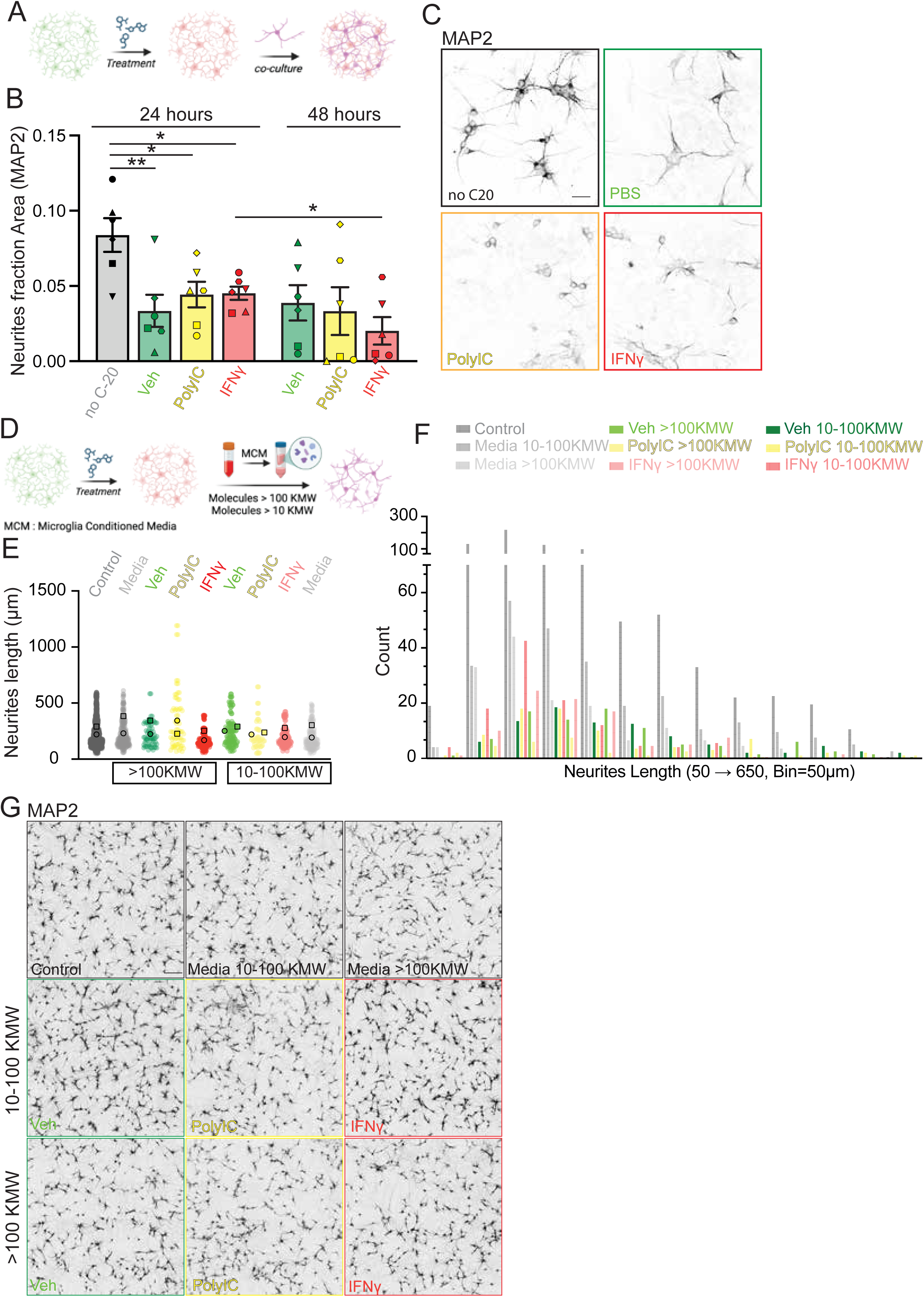
Disease-associated microglia curtail neurite outgrowth through high-molecular-weight soluble factors. **(A)** Schematic outlining the experimental setup for neuron–microglia co-culture assays. **(B)** Quantification of neurite coverage. The fractional area occupied by MAP2-positive processes per 20 × field was measured by NIS Element. Each symbol = different line; data are mean ± SEM. Statistical analysis: one-way ANOVA with Tukey’s post-hoc test: * p < 0.05, ** p < 0.01. **(C)** Representative high-magnification MAP2 images after 24 h of I3 neurons co-cultured with C20. Scale bar, 10 µm. **(D)** Schematic of the experiment assessing neuron responses to C20 conditioned media. **(E)** Plot of single-neurite lengths. Lengths were traced semi-automatically by NIS Element from stitched 10 × mosaics; ≥600 neurites per condition pooled from three independent experiments. **(F)** Histogram showing neurite length distributions under different treatment conditions. Frequency distribution (bin = 10 µm) for the nine treatment groups. **(G)** Low-magnification mosaics (4 ×, entire well) illustrating MAP2 network integrity in control neurons, neurons treated with un-fractionated media, or neurons treated with 10 KMW or 100 KMW fractions from PBS, poly I:C or IFNγ primed microglia. Scale bar, 250 µm.

To determine whether secreted microglial factors contribute to the neurotoxic spread, we treated neurons with conditioned media (CM) obtained from activated C20 cells. CM was fractionated into two aliquots: molecular weight (10–100 kDa) and high molecular weight (>100 kDa) (**Figure 5D**). PBS-treated media and freshly prepared C20 media fractions served as controls to exclude the potential confounding effects of serum-derived factors (FBS). Notably, CM from the high molecular weight fraction did not induce neurite retraction compared to control conditions (**Figure 5E-G**). Conversely, neurons exposed to CM from the lower molecular weight fraction derived from IFNγ-stimulated microglia exhibited subtle but consistent neurite retraction (**Figure 5E-G**). Importantly, FBS-derived molecules, in both molecular weight fractions, had no significant impact on neuronal health compared to PBS controls (**Figure 5E-G**). These findings strongly support the hypothesis that reactive microglia, overexpressing uPAR, negatively influence neuronal health through the well-described aberrant phagocytic capacity, rather than the spread of low and high-molecular weight soluble factors.

### CAR-T-cell Therapy Directed Against uPAR Efficiently Removes Pathogenic Microglia

Having established that uPAR is preferentially expressed in specific disease-associated, neurotoxic microglia, we then set out to develop a CAR-T cell-based strategy to selectively target and eliminate this pathogenic subpopulation using the cell-surface uPAR as a molecular target (uPAR–CAR-T). These uPAR–CAR-T cells were obtained by transducing human T lymphocytes with a lentiviral vector encoding a third-generation CAR and GFP marker^43^. Transduction efficiency of CAR-T was evaluated through GFP expression measured by flow cytometry (∼30%, **Supplementary Figure 13A**). The transduced cell pool was comparable to untransduced T cells, with a predominance of CD8⁺ over CD4⁺ cells (**Supplementary Figure 13B, C**). CAR-T cells were co-cultured overnight with C20 microglial cells pre-treated with polyIC or IFN-γ, and uPAR⁺ cell elimination was assessed by flow cytometry using DRAQ7 staining^44,45^. (**Supplementary Figure 13B**, gating strategy). No significant increase in cell death was observed in polyIC- or IFNγ-treated uPAR^+^ C20 cells alone. Co-culture with untransduced T cells resulted in a modest increase in cell death, from a baseline of ∼15% to ∼30% (a two-fold increase in mortality), likely reflecting stress from the co-culture conditions. Notably, PBS-treated C20 cells exposed to uPAR-CAR-T cells exhibited a nearly threefold increase in mortality (from ∼15% to ∼50%) (**Figure 6A, B**). The effect was even more pronounced in polyIC- and IFNγ-treated C20 cells, where uPAR expression induced an approximately fivefold increase in cell death (**Figure 3A, B; Figure 6A, B**). To confirm the relationships between these populations and their responses to treatments, we then took an unsupervised approach and plotted the flow cytometry results using the t-SNE algorithm (**Figure 6C**). Some of the clusters showed higher values of DRAQ7, indicating the cells were undergoing cell death. When we then annotated the clusters, the high DRAQ7 clusters belonged to the group of C20-treated cells with uPAR-CAR-T (cluster 3), specifically subclusters 8 and 9, which had a significantly higher DRAQ7 intensity corresponding to the C20 samples that were pre-treated with polyIC and INFψ.

**Fig. 6.**
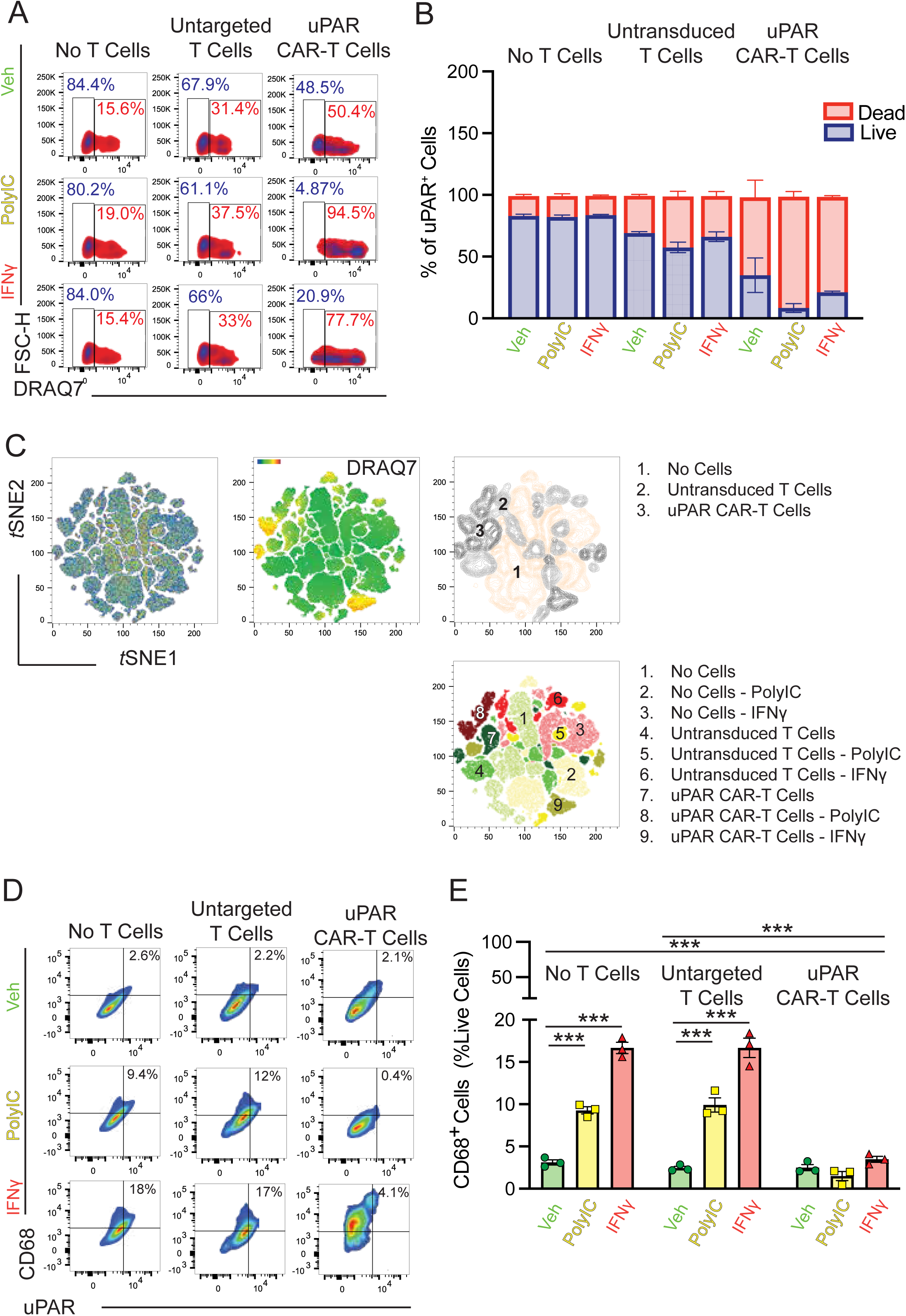
uPAR-CAR-T cells selectively eliminate disease-associated microglia and blunt their inflammatory phenotype. **(A)** Cytotoxicity assay design and representative viability plots. Shown are density plots of forward scatter (FSC-H, surrogate for size/complexity) versus the impermeant viability dye DRAQ7. The upper-right gate delineates DRAQ7⁺ (dead) microglia. **(B)** Quantification of CAR-mediated killing. Stacked bars display the mean fraction of live (DRAQ7⁻, blue) and dead (DRAQ7⁺, red) uPAR⁺ microglia for each condition (three independent donor-matched experiments, each performed in triplicate). **(C)** High-dimensional mapping of cell states. All viable events from the 27 samples (3000 cells per sample) were concatenated, arcsinh-transformed (co-factor 150) and embedded with t-SNE (perplexity = 30, 1000 iterations) using FlowJo v10.9.0. Left, composite plot; center, DRAQ7 signal overlaid (blue → yellow → red). **(D)** Phenotypic profiling of surviving microglia. Density plots (uPAR-PE, x-axis; CD68-BV421, y-axis) illustrate antigen persistence and phago-lysosomal activation after each treatment. **(E)** Bar graph shows the percentage of CD68⁺ cells within the live microglial gate. Data are means ± SEM from the same biological replicates as in (B). Statistical analysis: one-way ANOVA with Tukey’s post-hoc test; n = 9 (three independent experiments with three replicates per condition). ***p < 0.001.

To assess whether our treatment effectively reduced the population of specific uPAR-expressing disease-associated microglia, we quantified the proportion of CD68⁺/uPAR⁺ double-positive cells among the viable C20 cells following co-culture with T cells. Notably, treatment with uPAR-CAR-T cells led to a striking and selective decrease in CD68⁺/uPAR⁺ microglia, with levels dropping approximately 5- to 8-fold, from 10-20% to less than 2-5%. In contrast, treatment with untargeted CAR-T cells did not significantly reduce this population (**Figure 6D, E**). These *in vitro* results demonstrate that uPAR-CAR-T cells selectively and efficiently eliminate uPAR-expressing disease-associated microglia, including reactive populations. In addition, we employed an orthogonal approach using FITC-directed CAR-T cells and FITC-labeled anti-uPAR antibody to validate these findings further. C20 cells were incubated with FITC-conjugated anti-uPAR, allowing for selective labeling of uPAR-expressing cells, which could then be recognized by FITC-specific CAR-T cells (**Supplementary Figure 14A**). Following 24-hour incubation with an anti-FITC antibody, we observed a robust increase in fluorescence, confirming adequate labeling. However, when C20 cells were co-cultured with FITC-CAR-T cells, the proportion of uPAR⁺ cells was significantly reduced from ∼80% to ∼40% (**Supplementary Figure 14A, C**). To control the specificity of this effect, we conducted experiments using untransduced T cells, with or without FITC-anti-uPAR. In all control conditions, there was no significant reduction in cell viability, CD11b⁺ microglia, or uPAR⁺ microglia (**Supplementary Figure 14D, F**). Together, these results validate the selectivity of uPAR-directed CAR-T-cell therapy and establish the C20 microglial line as a relevant *in vitro* platform for assessing target engagement, specificity, and functional efficacy in developing microglia-targeting CAR-T strategies **uPAR- CAR-T Cells Do Not Elicit Neurotoxicity in Human iPSC-Derived Cortical Neurons** To evaluate whether uPAR-CAR-T cells have potential killing effects on neurons, we co-cultured human iPSC-derived i^3^-neurons with either untransduced T cells or uPAR-CAR-T cells using the same experimental conditions applied in the microglial assays (**Figure 7A**). In i^3^Neurons, uPAR was predominantly localized to the nucleus (**Figure 7B**). Neurite outgrowth was assessed *using* high-content imaging, similar to our previous experiments. We observed no reduction in average neurite length across all tested lines, including both control and C9-ALS/FTD-derived neurons (**Figure 7C, E**). These results indicate that uPAR-CAR-T cells do not compromise neuronal integrity, even in disease-relevant contexts. Together, these data demonstrate that the intracellular, specifically nuclear, localization of uPAR in neurons renders them inaccessible to CAR-T cell engagement, supporting the selectivity and safety profile of uPAR-CAR-T cell therapy for targeting specific disease-associated microglia in the CNS.

**Fig. 7.**
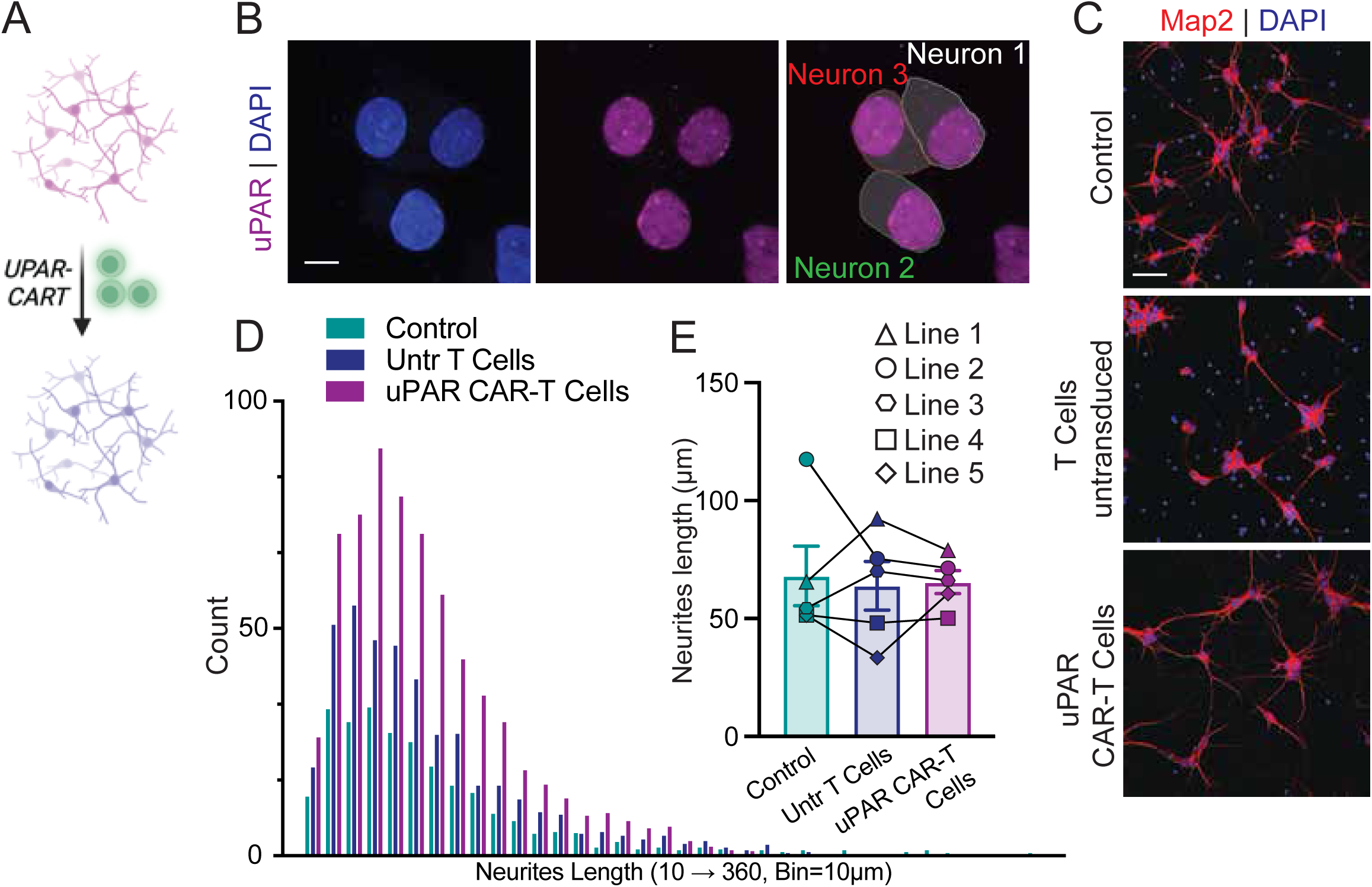
uPAR-CAR-T cells do not elicit neurotoxicity in human iPSC-derived cortical neurons. **(A)** Schematic of the experimental paradigm. Human iPSC-derived i3-neurons were generated using a doxycycline-inducible NGN2 cassette and matured for 21 days in Neurobasal medium supplemented with N2, B27, BDNF, and NT-3. Mature neurons from three controls and three C9-ALS/FTD lines were co-cultured overnight with either untransduced T cells or uPAR-CAR-T cells at a 1:5 (neuron:T cell) ratio under the same conditions used for microglial assays. **(B)** Representative confocal images of i3-neurons immunostained for uPAR (magenta) and DAPI (blue). In neurons, uPAR is localized exclusively to the nucleus. **(C, D)** High-content imaging of MAP2-positive neurites from representative control neurons exposed to untransduced or uPAR-CAR-T cells. Images were acquired with a Nikon A1 confocal microscope at 20× magnification, and neurite tracing and length distribution were quantified using the NIS-Elements software neurite analysis plugin (10 µm bin width). **(E)** Quantification of mean neurite length across five i3-neuron lines (Lines 1–5) reveals no reduction after exposure to uPAR-CAR-T cells compared with untransduced T cells or untreated controls. Data are presented as mean ± SEM.

## DISCUSSION

A central goal of ALS therapeutics is to halt the propagation of neurodegeneration by disrupting neuroinflammatory cycles that amplify disease spread. Our findings demonstrate that this objective can be achieved by selectively targeting and eliminating pathogenic microglial subsets using CAR-T cell-based immunotherapy. By integrating human ALS tissue analyses, spatial proteomics, transcriptomics, and functional *in vitro* modeling, we identified a candidate, uPAR, as a selective surface marker of pathogenic and disease-associated microglia in ALS. We demonstrated that it can be exploited to enable a highly specific and compelling CAR-T cell-based immunotherapy that selectively eliminates toxic uPAR^hi^ disease-associated microglia. Briefly, we found that uPAR is markedly upregulated in microglia within degenerating cortical and spinal cord regions of ALS postmortem tissue, while remaining virtually undetectable in unaffected areas or control brains. uPAR^hi^ microglia were also present in stroke samples, consistent with injury-evoked microglial activation, supporting uPAR as a marker of disease-amplifying, rather than homeostatic, microglia across various contexts of neuroinflammation. Using a human microglial cell model (C20), we demonstrated that inflammatory stimuli robustly induce surface uPAR expression on C20 cells, providing a controlled platform for testing CAR-T cell engagement and specificity. A uPAR-directed CAR-T construct efficiently induced elimination of activated, uPAR⁺ C20 microglia but not in uPAR⁻ microglia or human iPSC-derived neurons, establishing apparent antigen selectivity and functional specificity. These findings provide foundational evidence for uPAR-CAR-T cell selectivity and support ongoing preclinical studies aimed at eliminating disease-amplifying microglia, reducing neuroinflammation, and potentially slowing the progression of ALS.

Microglia have emerged as critical contributors to the pathogenesis of ALS^17^. Extensive transcriptomic and histological studies show that these CNS-resident macrophages transition into a pro-inflammatory state in neurodegenerative settings, characterized by increased expression of genes involved in cytokine signaling, antigen presentation, phagocytosis, and tissue remodeling^46^. While such activation may initially serve a protective function, persistent activation leads to synaptic stripping, oxidative stress, and neuronal loss^47^. Several treatments have been tested in animal models and, to some extent, in patients, to prevent the pro-inflammatory action of microglia. However, these anti-inflammatory therapies have been only moderately successful, most likely due to the complex nature of neuroinflammation, which involves various cellular and molecular pathological mechanisms and produces numerous inflammatory mediators, particularly cytokines, which are challenging to target with a single drug treatment^48^. This suggests that perhaps ablation of microglia may be required to halt this inflammatory cascade at its source. Experimental studies in rodent models of ALS have shown that pharmacological or genetic ablation of microglia reduces neuronal death and delays disease progression^49^. However, this approach could also pose a threat to the CNS^50^, further supporting the concept that targeted microglial removal could offer a more effective therapeutic benefit. However, current small-molecule approaches face substantial translational barriers due to off-target toxicity and lack of cellular specificity. CAR-T cells, in contrast, offer a means of precisely directing cytotoxicity to pathologically altered cells through surface antigen recognition.

uPAR is a glycosylphosphatidylinositol (GPI)-anchored receptor best known for its roles in cell motility, extracellular matrix remodeling, and immune signaling. Although largely absent from the adult CNS under homeostatic conditions, it becomes strongly induced in states of inflammation, ischemia, and neurodegeneration^33,51^. In ALS, uPAR expression has been observed in reactive microglia and astrocytes in degenerating motor regions^52^. We found that uPAR expression marks a distinct, compartmentalized uPAR^hi^ microglial subpopulation enriched in degenerating regions, where these cells often co-express IBA1 and CD68, form dense clusters in spinal and cortical white matter, and localize within microglial–astrocytic and vascular niches that support region-specific propagation of inflammatory signaling. These findings extend recent models of CNS-compartmentalized neuroinflammation, in which chronic microglial activation and microglia-astrocyte crosstalk perpetuate injury within closed CNS niches^53^. This phenotype aligns with terminally activated, degeneration-associated microglial states described in MS and AD, reinforcing uPAR^hi^ microglia as central effectors that shape the inflammatory landscape of the ALS CNS and highlighting uPAR as both a pathological hallmark and a therapeutic entry point for microglia-targeted immunomodulation. Consistent with these spatial proteomic findings, stimulation of C20 human microglia with pro-inflammatory agents drove robust upregulation of surface uPAR alongside CD68 and CD80, mirroring activation states reported in DAM across ALS/FTD tissues and models^54^. These uPAR⁺ microglia-like cells induced neurite shortening in co-cultured human iPSC-derived neurons, supporting a functional link between uPAR expression and pathogenic microglial activity. Importantly, uPAR-directed CAR-T cells selectively engaged and eliminated these activated C20 cells in an antigen-dependent manner while sparing uPAR^low^/–microglia and human iPSC-derived neurons. Neuronal resistance reflects the predominantly nuclear localization of uPAR under healthy conditions, shielding it from CAR recognition, as well as potential expression of inhibitory ligands such as PD-L1 that suppress T-cell activation^55^. Together, these findings demonstrate that uPAR identifies a highly reactive, disease-amplifying microglial subset and that uPAR-targeted CAR-T cells can eliminate these populations with marked specificity, offering a mechanistic rationale for selectively extinguishing compartmentalized neuroinflammation while preserving neural integrity.

Our findings have important translational implications. The use of CAR-T cells in the CNS, once considered prohibitively difficult, is now actively explored in glioblastoma and multiple sclerosis^3,49^. Moreover, uPAR CAR-T cells have effectively eliminated senescent cells in aging and fibrotic mouse models, reduced systemic inflammation, and improved tissue function^25,35,36^. These precedents demonstrate the technical feasibility and clinical relevance of adapting CAR-T technology to target CNS-resident cell types in the treatment of chronic diseases. The specificity of uPAR expression in diseased microglia, its low expression at baseline in other adult tissues, and the lack of neuronal toxicity in our *in vitro* system make this an exceptionally promising target candidate for translational development in ALS. *In vivo* validation in ALS animal models will be crucial to establishing the CNS homing, persistence, safety, and efficacy of uPAR CAR-T cells. Regional delivery, modulation of the blood-brain barrier, or chemokine receptor engineering may enhance CNS access. Approaches to reducing cytokine release syndrome (CRS) are already being explored, including the incorporation of inducible “kill-switch” or safety-switch mechanisms that enable rapid, pharmacologically controlled ablation of CAR-T cells in the event of excessive immune activation^56^. Additional specificity could be engineered through dual-antigen recognition systems, further restricting activity to microglia with a disease-associated phenotype^57^. Ultimately, combining CAR-T cell therapy with small molecules that target neuroinflammation, proteostasis, or neurotrophic support may offer synergistic disease-modifying benefits.

In conclusion, this work establishes uPAR as a tractable immunotherapeutic entry point for selectively eliminating disease-amplifying microglia and demonstrates the feasibility of using uPAR-directed CAR-T cells to modulate the neuroinflammatory milieu that drives ALS progression. Together, these findings provide a foundational strategy for advancing microglia-targeted CAR-T therapy as a transformative approach to reset neuroinflammation in ALS and other glia-driven CNS disorders.

## MATERIALS AND METHODS

### Cell cultures

C20 human microglial cells were maintained in a humidified incubator at 37°C with 5% CO_2._ The culture medium consisted of DMEM-High Glucose supplemented with 10% FBS and penicillin–streptomycin. Cells were passaged at a 1:10 split ratio every 4 days. For passaging, cultures were rinsed once with PBS, detached with 0.05% trypsin–EDTA for ≤ 3 min at 37°C, neutralized with complete medium, pelleted at 300 × g for 5 min, and reseeded at 1 × 10⁵ cells cm⁻² in fresh growth medium. I^3^-neurons were derived from human iPSCs harboring a doxycycline-inducible NGN2 cassette. Six well-characterized lines were used: three from healthy controls and three from C9-ALS patients (**Table 1**). hiPSCs were expanded on Matrigel-coated plates in E8 medium and routinely tested for mycoplasma. For differentiation, confluent cultures were dissociated into single cells using Accutase and replated on Matrigel in induction medium consisting of Neurobasal supplemented with N2 and 10 µg/mL⁻doxycycline. After 3 days, induced neurons were dissociated and transferred onto poly-L-ornithine-coated surfaces at 1.6 × 10⁵ cells mL⁻¹ in maturation medium (Neurobasal + N2 + B27 + 50 ng mL⁻¹ BDNF + 50 ng mL⁻¹ NT-3). Half-medium changes were performed twice weekly. Doxycycline was present only during the first 4 days of maturation. Neurons reached functional maturity by day 21 (DIV21) and were used for downstream assays at this stage. For i^3^-neurons-C20 cells co-culture, C20 cells were first activated for 48 h. After treatment, cells were detached with 0.05% trypsin, washed twice with PBS, and resuspended in i^3^-neuron maturation medium. Activated C20 cells were added to DIV21 i^3^-neurons at a 1:5 neuron-to-microglia ratio and maintained for 48 h before analysis. For the conditioned media experiments, the supernatant from C20 cultures that had been stimulated for 48 h was collected, centrifuged at 2,000 × g for 30 min at 4°C to remove debris, and then sequentially filtered through 100 kDa and 10 kDa molecular-weight cutoff Amicon units. The >100 kDa (100 KMW) and 10–100 kDa (10 KMW) fractions were retained, while the <10 kDa flow-through was discarded. Conditioned fractions were added to DIV21 i^3^-neurons for 48 h, after which cells were processed for viability and neurite assays. Immunofluorescence analysis: Cells were fixed in 4% PFA (20 min, 37°C), rinsed in PBS, and permeabilized/blocked for 1 h at room temperature in PermBlock buffer (0.1 % Triton X-100 + 10% FBS + 1% BSA). Primary antibodies were applied overnight at 4 °C in PBS + 0.01% BSA. After two PBS washes, fluorescent secondary antibodies were added for 1 h at room temperature. Nuclei were counterstained with Hoechst (1:3,000, 10 min). Samples were stored in PBS until imaged on a Nikon A1R confocal microscope using NIS-Elements software. Z-stacks or single-plane images were collected with 20× and 63× objectives; additional digital zoom was applied where noted. Mean fluorescence intensity (MFI) per cell was quantified in NIS-Elements. Neurite length and total neurite area were measured with the NIS-Elements neurite-tracing plug-in. Graphs and statistics were generated in Prism v10.4.1.

### Western Blot Analysis of Cell Lysates

Cultured cells were lysed for 15 min on ice in RIPA buffer supplemented with protease and phosphatase inhibitors. Lysates were clarified (500 × g, 5 min, 4 °C), and the supernatants were collected. Protein concentration was determined by BCA assay. Equal amounts of protein (50 µg lane⁻¹) were resolved on 4–15% gradient SDS-PAGE gels at 100 V for 1 h and transferred to nitrocellulose using a semi-dry apparatus. Membranes were blocked for 1 h in TBS-T containing 5% non-fat milk, then incubated overnight at 4 °C with primary antibodies diluted in TBS-T + 5% BSA. After two washes in TBS-T, membranes were exposed to HRP-conjugated secondary antibodies for 1 h at room temperature, developed with femto-sensitivity EC, and imaged on a ChemiDoc-XR system. Total-protein loading was verified by Ponceau S staining or UV detection.

### CAR-T Co-cultures

C20 cells were pre-activated for 48 h, harvested, and plated in fresh C20 medium. Cryopreserved uPAR-CAR-T cells were thawed, washed twice, resuspended in C20 medium, and counted with trypan blue exclusion. CAR-T and C20 cells were cocultured overnight (≈16 h) at a 1:5 microglia-to-CAR-T cell ratio. For neuron co-cultures, uPAR-CAR-T cells were thawed, washed, and resuspended in i^3^-neuron maturation medium, then added to DIV21 neurons at a 1:5 neuron-to-CAR-T cell ratio. Co-cultures were incubated overnight before downstream analyses.

### Human Post-Mortem Tissues

Frozen and formalin-fixed paraffin-embedded (FFPE) motor cortex and lumbar spinal cord samples from control and C9-ALS donors were obtained from the Weinberg ALS Center and TargetALS biobanks (**Table 2**). All tissue procurement, processing, and distribution were conducted in accordance with an IRB-approved protocol, and informed consent was obtained from all donors or their next kin. Samples were used for multiplexed IF and immunoblotting. Western Blot analysis: Approximately 1 mg of wet weight from the mid-motor cortex or cervical spinal cord was homogenized in 1% SDS using an automated Dounce homogenizer for 10 min at maximum speed. Homogenates were centrifuged (1,000 × g, 5 min, room temperature); the supernatants were retained, and the pellets were discarded. IF analysis: FFPE sections were cut onto charged slides, de-paraffinized in xylene, and rehydrated through graded ethanol (100% to 20%). After two PBS rinses, antigen retrieval was performed using a proprietary epitope-retrieval solution (IHC-Tek Epitope Retrieval Solution, IHC World). Sections were permeabilized in 0.5% Triton X-100 + 5% BSA (1 h, room temperature) and incubated overnight at 4°C with primary antibodies diluted in PBS + 0.01% BSA. After two PBS washes, slides were incubated with appropriate fluorescent secondaries (1 h, room temperature), counterstained with Hoechst (1:3,000 in PBS, 10 min.), and mounted in aqueous medium before storage at 4 °C for confocal imaging.

### CAR Design and Lentiviral Production

The CAR incorporated an anti-human CD87 single-chain variable fragment (scFv) derived from clone 09 (ab275627, Abcam) assembled in a heavy-to-light (H2L) format using a (G₄S)₄ linker. The scFv was connected to a CD8α hinge and CD8α transmembrane domain, which in turn was fused to CD28 and 4-1BB costimulatory regions and a CD3ζ activation domain, consistent with a third-generation CAR design^58^. CAR-expressing lentivector production and CAR-T manufacture were conducted as previously described^58^. CAR expression inserts were synthesized (GenScript Biotech) and inserted into the pCDH-EF1α-MCS-T2A-copGFP transfer backbone (CD526A-1, System Biosciences) using XbaI/BamHI restriction digest. All transfer, packaging, and envelope plasmids were propagated in NEB Stable Competent *E. coli* (C3040H, New England Biolabs). Bacteria were grown in Miller LB broth (BP1426-2, Fisher Scientific) supplemented with 1.9% Bacto Yeast Extract (212750, Thermo Fisher Scientific) and 100 µg/mL ampicillin (A8351, Sigma-Aldrich). Plasmids were purified from overnight cultures with the PureLink Expi Endotoxin-Free Maxi Kit (A31231, Thermo Fisher Scientific). DNA pellets were dissolved in endotoxin-free water and stored at −20 °C until use. Lentiviral supernatants were generated by transfecting poly-D-lysine-coated flasks (5 µg/cm²; 354210, Corning) of near-confluent, low-passage, HEK293T/17 cells (CRL11268, ATCC) with pCDH transfer vector, pRSV-Rev (12253, Addgene), pMDLg/pRRE (12251, Addgene), and pMD2.G (12259, Addgene) using Lipofectamine 3000 (L3000150, Thermo Fisher Scientific). Culture media was Advanced DMEM (12491023, Thermo Fisher Scientific) supplemented with 5% heat-inactivated FBS (A38400-01, Gibco) and 1X GlutaMAX (35050-061, Gibco). Media was replaced 6 hours post-transfection and viral supernatants were collected at 24 and 48 hours, combined, filtered (0.45 µm), and concentrated 200-fold by precipitation overnight at 4 °C using 4x diluted PEG-8000 (BP233-1, Fisher Scientific), followed by centrifugation at 1600×g for 1 hour at 4 °C and resuspension in a storage buffer composed of 10 mM Tris (pH 7.4; 648315, EMD Millipore), 10% lactose (61339, Sigma-Aldrich), and 25 mM proline (81709, Sigma-Aldrich) in DPBS. Viral stocks were stored at −80 °C. For titer determination, HEK293T/17 cells were transduced with lentiviral supernatant in the presence of 0.8 µg/mL Polybrene (TR-1003, Millipore Sigma), and GFP-positive cells were quantified on a BD FACSymphony A5 SORP cytometer (BD Biosciences).

### CAR-T production

Human T cells were isolated from PBMCs obtained from healthy donor leukopaks (200-0092, Stemcell Technologies). Negative magnetic selection (130-096-535, Miltenyi Biotec) was used to isolate T cells. T cells were cultured in RPMI-1640 (10-041-CV, Corning) supplemented with 10% heat-inactivated FBS (A38400, Gibco), 1X ITS-G (41400045, Gibco), 10 mM N-acetyl-L-cysteine (A9165, Millipore Sigma), 1X GlutaMAX (35050061, Gibco), 1X Glucose (A24940, Gibco), 1X Sodium Pyruvate (11360070, Gibco), 1X MEM Non-Essential Amino Acids (11140050, Gibco), 1X HEPES Buffer (15630080, Gibco), 1X Penicillin-Streptomycin (15140122, Gibco), and 55 uM 2-Mercaptoethanol (21985023, Gibco). At the start of culture, T cells were seeded at 1×10⁶ cells/mL. Cells were activated using CD3/CD28/CD2 beads (1:1 bead-to-cell ratio, 130-091-441, Miltenyi Biotec) in media supplemented with 10 ng/mL IL-7 and 10 ng/mL IL-15 (BRB Preclinical Biologics Repository, NCI Biological Resources Branch, Frederick, MD). After 24 hours of activation, T cells were transduced with lentivirus at an MOI of 5 in the presence of 0.8 µg/mL Polybrene. Activation beads were removed magnetically 72 hours after stimulation. Cells were transitioned to G-Rex plates (80240M, Wilson Wolf) and maintained in cytokine-supplemented medium. IL-7 and IL-15 were replenished every three days. A full medium exchange was performed on Day 9, and CAR-T cells were harvested on Day 14. Transduction efficiency (% GFP-positive cells) was quantified on a BD FACSymphony A5 SORP cytometer (BD Biosciences). For cryopreservation, T cells were resuspended at 20×10⁶ cells/mL in CryoStor CS10 (07930, Stemcell Technologies). Before experiments, frozen CAR-T cells were slowly thawed into pre-warmed (37°C) RPMI-1640.

### Flow Cytometry

C20 microglia were seeded in 6-well plates (2 × 10⁵ cells per well⁻¹). After 24 h, cultures received one of the following stimuli: LPS 1 µg mL⁻¹ ; R848 1 µg mL⁻¹ ; Pam3CSK4 10 µg mL⁻¹ ; polyIC 10 µg mL⁻¹ ; TNF-α 50 ng mL⁻¹ ; or IFNγ 50 ng mL⁻¹. Fresh aliquots of the same reagents were re-added at 24 h to complete a 48-h exposure. Cells were detached with 0.05% trypsin–EDTA for 2 min at 37°C, the reaction was quenched with PBS + 10% FBS, and suspensions were pelleted at 500 × g for 5 min at 4°C. Pellets were resuspended in 100 µL PBS containing DRAQ7 for viability discrimination, incubated on ice for 30 min, diluted to 1 mL with PBS, and washed once more (500 × g, 5 min, 4 °C). Final staining was performed in flow buffer (PBS + 10% FBS + 1% BSA) with the following antibodies: CD80-APC-Cy7, CD87/uPAR-PE, and CD68-BV421. Samples were incubated at 37°C for 20 min, washed once in PBS, and kept on ice until acquisition. Cryopreserved uPAR-CAR-T cells and donor-matched untransduced T cells were thawed and rested overnight in DMEM-High Glucose supplemented with 10% FBS and penicillin–streptomycin. Cells were pelleted (450 × g, 8 min, 4 °C) and labeled for 30 min at 4°C with LIVE/DEAD™ Fixable Aqua in PBS, followed by staining in flow buffer with the antibody panel (CD80-APC-Cy7, CD87/uPAR-PE, and CD68-BV421). After a final wash (500 × g, 5 min, 4 °C), the samples were maintained on ice for cytometric analysis, which was acquired on a BD Symphony A5 equipped with FACSDiva v8.0.1. UltraComp eBeads were used for single-color compensation. A minimum of 1 × 10⁵ live events was collected per sample. Data were analyzed in FlowJo v10 (BD Biosciences) with sequential gating for singlets, viability dye exclusion, and marker-positive populations. For multidimensionality reduction we used FlowJo v10 using the following parameters: FSC-H, SSC-H, CD87/uPAR-PE and DRAQ7.

### Study Group

A tissue microarray (TMA) was constructed from a cohort of 24 post-mortem human CNS samples obtained from the Weinberg ALS Center biobank. The cohort was designed to enable regional and genotypic comparisons, encompassing four distinct anatomical regions: motor cortex, spinal cord, frontal cortex, and occipital cortex. For each region, five conditions were included, each represented by a single TMA core. This comprised two control samples (one non-neurological control and one stroke patient) and four ALS patients, including two sporadic ALS (sALS), one C9ORF72-mutation (C9_ALS), and one SOD1-mutation (SOD1_ALS) case. P-TDP-43 aggregates in affected areas of the different cohorts are provided in **Supplementary Figure 15**.

### Spatial Proteomics Profiling

Multiplexed imaging was performed on consecutive TMA sections using the PhenoCycler-Fusion 2.0 platform (Akoya Biosciences). A 23-plex antibody panel was employed **(Table 3),** combining markers from the commercial STP6001 STEP Core Panels for “Neuro Core” and “Neuroinflammation” with two custom-conjugated antibodies: uPAR (R&D Systems, AF807), phospho-TDP43 (ProteinTech, 22309-1-AP). For each TMA section, raw imaging data were acquired as multi-channel QPTIFF files.

### Cell Segmentation and Quality Control

Image processing and downstream analysis were conducted using the SPACEc workflow^59^. Cell segmentation was performed independently on each TMA core independently using the *sp.tl.cell_segmentation* function, which implements Mesmer, a deep learning model^60^ using whole-cell mode (compartment=“whole-cell”), with the DAPI channel specified as the nuclear marker (nuclei_channel=“DAPI”). We assessed segmentation quality by visually inspecting a random subset of images using the *sp.pl.show_masks* function with DAPI as the nucleus channel and IBA1 as an additional channel to validate cytoplasmic boundaries (**Supplementary Figure 16**). For further visual inspection, segmented objects were imported into TissUUmaps ^61^. Cells within regions exhibiting tissue artifacts (e.g., folds, tears, air bubbles) or low signal intensity were manually annotated and excluded. Subsequently, the matrices from all cores of the same TMA section were concatenated using *sp.pp.read_segdf*. A quality filtering step was then applied with *sp.pp.filter_data* to remove the lowest 1% of cells based on both cell area and DAPI intensity (**Supplementary Figure 17**). The expression matrix was normalized using an arcsinh transformation (cofactor = 150) with the *sp.pp.format* function, following the standard normalization best practices for CODEX imaging^62,63^. Markers exhibiting non-specific background staining (CD80, P16, Ki67) were excluded from subsequent analysis, and phospho-TDP43 was used only as a pathological marker for subcellular localization.

### Cell Phenotyping and Spatial Analysis

Processed data, including cell metadata and expression matrices, were stored as an AnnData object for subsequent analysis in Python v3.12.2. Unsupervised clustering was performed with Scanpy, testing multiple Leiden resolutions (0.3-0.8) across n_neighbors parameters (10-15). Optimal parameters were empirically chosen to best resolve canonical cell populations. Cell types were annotated based on the expression of canonical lineage markers: Neurons: (NeuN, MAP-2), Oligodendrocytes (Olig-2), Endothelial cells (Collagen IV, Claudin-5, Vimentin), Vascular cells (SMA), Ependymal (Vimentin), Astrocytes (GFAP, AQP4, Vimentin) M2-like Macrophages, (CD163) and Microglia cells (TMEM119, Iba-1, CD68). Further, to investigate microglial heterogeneity, the microglia populations were subclustered. The subpopulations were re-annotated based on high expression of key activation markers, defining microglial states characterized by high Iba1 (micro_hIBA1) or high CD68 (micro_hCD68) expression. The microglia subclusters with baseline expression of TMEM119, Iba-1, and CD68, were classified as resting microglia (micro_resting). Cell clusters were visualized in two dimensions using Uniform Manifold Approximation and Projection (UMAP) (**Supplementary Figure 3**). Further, the combined microglia population from all states was classified as Microglia_ALL. A population of unidentifiable cells was excluded to prevent misclassification and minimize noise signals in downstream analysis. Finally, the spatial concordance of the final annotations was validated by overlaying the cells with the channels of canonical markers, confirming the agreement between the cell classification and protein localization (**Supplementary Figure 18**).

### uPAR^+^ Thresholding and Cellular Neighborhood Analysis

To account for the ubiquitous baseline expression of uPAR across cell types (**Supplementary Figure 19**), a tissue-specific positive threshold was applied, as previously described^64^. Within each TMA core, cells were classified as uPAR^+^ if their expression level exceeded two times the mean uPAR expression of all cells in that TMA core. Cell type proportions were calculated by normalizing the number of each cell type to the total number of cells within each individual TMA core, yielding cell percentages. To characterize the spatial organization of uPAR^+^ cells and their interaction with neighboring cell types within each CNS tissue, we performed a cellular neighborhood (CN) analysis using the *sp.tl.neighborhood_analysis* function from SPACEc^59^. This analysis was conducted independently for each tissue. The optimal number of cellular neighborhoods for each dataset was determined using the elbow method, following initialization of the spatial graph with a k-nearest neighbors’ parameter of k = 20 and a targeted search space of n_neighborhoods = 20 for subsequent Leiden/k-means clustering. The optimal k value identified from this method was subsequently used for the final CN calculation (**Supplementary Figures 6-9 A**). To evaluate the enrichment of specific cell types within each CN, we generated expression heatmaps with *sp.pl.cn_exp_heatmap* functions. The log_2_ fold-change over tissue average measures how abundant or enriched a specific cell type is within a given CN, compared to its average abundance across the entire tissue section (**Supplementary Figures 6-9 B**). Positive values indicate that a cell type is more abundant in each CN compared to the tissue-wide average, whereas negative values indicate depletion. The resulting neighborhoods were visualized using the spatial scatter plot (catplot) with the *sp.pl.catplot* function (**Supplementary Figure 6-9 C**). Finally, spatial context maps per condition were computed with *sp.tl.build_cn_map* to define the frequency of each CN using a threshold of 0.85 for the percentage of cells required to assign a CN (**Supplementary Figures 6-9 D-I**).

### RNAseq and scRNAseq analysis of existing datasets

RNA-sequencing data from human ALS tissues were obtained from the GEO super series GSE137810 and analyzed using reference-based cell-aware deconvolution. Cell-type proportions were estimated with ADAPTS^65^ using a single-nucleus RNA-seq reference from the Allen Brain Institute (Human MTG 10x SEA AD) to generate a cell-type signature matrix. Cell-type–specific differential expression was assessed using TOAST^66^ by modeling bulk expression as a function of experimental group and estimated cell-type proportions. TOAST-derived, composition-adjusted normalized expression values and gene-level contrasts were used directly for all downstream analyses. Microarray expression data from spinal cords of SOD1^G93A^ mice across disease stages were retrieved from GSE18597^22^, and RNA-seq data from rNLS8 TDP-43 mice were obtained from NCBI Sequence Read Archive (Accession Number PRJNA624791)^23^. All datasets were imported into R as raw or rlog-transformed counts and log-transformed when appropriate for downstream visualization and analysis. Single-nucleus RNA-sequencing data from patient tissues (GSE219281) were examined using the public browser interface (https://brainome.ucsd.edu/C9_ALS_FTD/); plots were exported and post-labeled for clarity.

## Acknowledgments

We thank the members of the Weinberg ALS Center for their critical reading of the manuscript, valuable insights, and collegial feedback.

## Funding

DoD grant HT9425-23-1-0258 (DT, ARH, AES); ALS United Mid-Atlantic (PP); The Farber Family Foundation (PP, DT, ARH); The Aldrich Foundation (PP).

## Authors Contribution

Conceptualization: MEC, DT, PP, ARH

Methodology: KK, SSM, DT-M, CS, BO, IMS

Investigation: MEC

Visualization: MEC, DT-M

Funding acquisition: DT, ARH, PP, AES

Project administration: DT, PP

Supervision: PP, DT

Writing – original draft: MEC

Writing – review & editing: DT, PP, ARH

## Competing interests

The Authors declare no competing interests.

## Data and materials availability

The pipelines used for spatial proteomics analysis are available on GitHub at: https://github.com/GenomeGaia/ALS_multiplex. The QPTIFF files, as well as the original segmented files and final cell metadata and expression matrices saved in AnnData format and will be available at datadryad.org.

## LIST OF SUPPLEMENTARY MATERIALS

**Table 1.** Human induced pluripotent stem cell (iPSC) lines utilized for neuron derivation. Source and demographic information for i^3^Neuron lines used in this study Figures 4-6. NA = not applicable, NS = not specified

| **Genotype** | **GroupID** | **Age of Onset** | **Sex** | **C9 RepeatSize** | **insert / site** | **source** |
| --- | --- | --- | --- | --- | --- | --- |
| **C9orf72** | ALS | 49 | Male | >44 | Ngn1/2 + mCherry in CLYBL locus | Barmada lab |
| **C9orf72** | ALS | 56 | Male | >25 | Ngn1/2 + mCherry in CLYBL locus | Barmada lab |
| **Control** | CTRL | NA | Male | normal | Ngn1/2 + iRFP in CLYBL locus | Barmada lab |
| **Control** | CTRL | NA | Female | normal | Ngn1/2 + mCherry in CLYBL locus | Barmada lab |
| **C9orf72** | ALS | NS | Female | NS | Ngn2 + mCherry in AAVS1 locus | Ward lab |
| **Control** | CTRL | NA | Male | normal | Ngn2 in AAVS1 locus | Ward lab |

**Table 2.** Demographic characteristics of the patient’s cohort.

| Patient | Source | Sex/Race | age at DOD | Autopsy Interval | Diagnosis | Notes | Experiment | Experiment 2 | Experiment 3 | Experiment 5 |
| --- | --- | --- | --- | --- | --- | --- | --- | --- | --- | --- |
| D | Target ALS | M/C | 59 | N/D | Control | Metastatic synovial sarcoma | WB Fig 1 | WB Supp Fig1 |  |  |
| F | Target ALS | M/B | 71 | N/D | Control | Respiratory failure secondary to lung cancer and infection | WB Fig 1 | WB Supp Fig1 |  |  |
| G | Target ALS | M/C | 89 | N/D | Control | Cardiorespiratory failure; Pneumonia, myodysplastic syndrome, acute renal failure | WB Fig 1 | WB Supp Fig1 |  |  |
| H | Target ALS | M/C | 82 | N/D | Control | N/A | WB Fig 1 | WB Supp Fig1 |  |  |
| JWAC 6 | Jefferson Weinberg ALS Center Biobank | F/C | 91 | 16-20 hours | Control | ----- | WB Fig 1 | WB Supp Fig1 | Multiplex Akoya | IF Fig 1-Supp Fig1 |
| N1 | NIH | F | 70 | N/D | Control | ----- | WB Fig 1 |  |  |  |
| N2 | NIH | M | 72 | N/D | Control | ----- | WB Fig 1 |  |  |  |
| JWAC | Jefferson Weinberg ALS Center Biobank | M/C | 63 | 16-20 hours | Control | stroke | WB Fig 1 |  | Multiplex Akoya |  |
| A | Target ALS | M/C | 59 | N/D | C9orf72 | ----- | WB Fig 1 | WB Supp Fig1 |  |  |
| B | Target ALS | F/C | 55 | N/D | C9orf72 | ----- | WB Fig 1 | WB Supp Fig1 |  |  |
| C | Target ALS | M/C | 66 | N/D | C9orf72 | ----- | WB Fig 1 | WB Supp Fig1 |  |  |
| I | Target ALS | F/C | 61 | N/D | C9orf72 | ----- | WB Fig 1 | WB Supp Fig1 |  |  |
| J | Target ALS | M/C | 68 | N/D | C9orf72 | ----- | WB Fig 1 | WB Supp Fig1 |  |  |
| JWAC 29 | Jefferson Weinberg ALS Center Biobank | M/C | 55 | 44-48 hours | C9orf72 | ----- | WB Fig 1 | WB Supp Fig1 | IF Fig 1-Supp Fig1 |  |
| JWAC 104 | Jefferson Weinberg ALS Center Biobank | M/C | 54 | 16-20 hours | C9orf72 | ----- | WB Fig 1 | WB Supp Fig1 | IF Fig 1-Supp Fig1 |  |
| JWAC 18 | Jefferson Weinberg ALS Center Biobank | M/C | 80 | 44-48 hours | sALS | ----- | WB Fig 1 |  |  |  |
| JWAC 30 | Jefferson Weinberg ALS Center Biobank | M/C | 48 | 12-16 hours | sALS | ----- | WB Fig 1 |  |  |  |
| JWAC 60 | Jefferson Weinberg ALS Center Biobank | M/C | 55 | 20-24 hours | sALS | ----- | WB Fig 1 |  |  |  |
| JWAC 1 | Jefferson Weinberg ALS Center Biobank | F/C | 63 | 44-48 hours | sALS | ----- | WB Fig 1 | WB Supp Fig1 |  |  |
| JWAC 34 | Jefferson Weinberg ALS Center Biobank | F/C | 69 | 20-24 hours | sALS | ----- | WB Fig 1 | WB Supp Fig1 |  |  |
| JWAC 11 | Jefferson Weinberg ALS Center Biobank | M/C | 58 | 12-16 hours | C9orf72 | ----- | WB Supp Fig1 | IF Fig 1-Supp Fig1 | Multiplex Akoya |  |
| JWAC 2 | Jefferson Weinberg ALS Center Biobank | F/C | 63 | 44-48 hours | C9orf72 | ----- | WB Supp Fig1 | IF Fig 1-Supp Fig1 |  |  |
| N3 | NIH | M/C | 58 | N/D | SOD1-ALS | ----- | WB Supp Fig1 |  |  |  |
| N4 | NIH | M/C | 55 | N/D | SOD1-ALS | ----- | WB Supp Fig1 |  |  |  |
| JWAC 46 | Jefferson Weinberg ALS Center Biobank | M/C | 42 | 4-8 hours | SOD1-ALS | ----- | WB Supp Fig1 |  | Multiplex Akoya |  |
| JWAC 42 | Jefferson Weinberg ALS Center Biobank | M/C | 59 | 12-16 hours | sALS | ----- | WB Supp Fig1 | IF Fig 1-Supp Fig1 |  |  |
| JWAC 47 | Jefferson Weinberg ALS Center Biobank | M/C | 73 | 18-12 hours | sALS | ----- | IF Fig 1-Supp Fig1 |  |  |  |
| JWAC 72 | Jefferson Weinberg ALS Center Biobank | M/C | 72 | 4-8 hours | sALS | ----- | IF Fig 1-Supp Fig1 |  |  |  |
| JWAC 93 | Jefferson Weinberg ALS Center Biobank | F/C | 71 | 8-12 hours | sALS | ----- | IF Fig 1-Supp Fig1 |  |  |  |
| JWAC 121 | Jefferson Weinberg ALS Center Biobank | F | 62 | 44-48 hours | sALS | ----- | IF Fig 1-Supp Fig1 |  | Multiplex Akoya |  |
| JWAC 125 | Jefferson Weinberg ALS Center Biobank | M/C | 64 | 48-72 hours | sALS | ----- | IF Fig 1-Supp Fig1 |  | Multiplex Akoya |  |
| N/D = Not Disclosed; F = Female; M = Male; C = Caucasian; B=Black; WB=western blot; IF=immunofluorescence; sALS=sporadic ALS |  |  |  |  |  |  |  |  |  |  |
| JWAC=Jefferson Weinberg ALS Center |  |  |  |  |  |  |  |  |  |  |

**Table 3.**
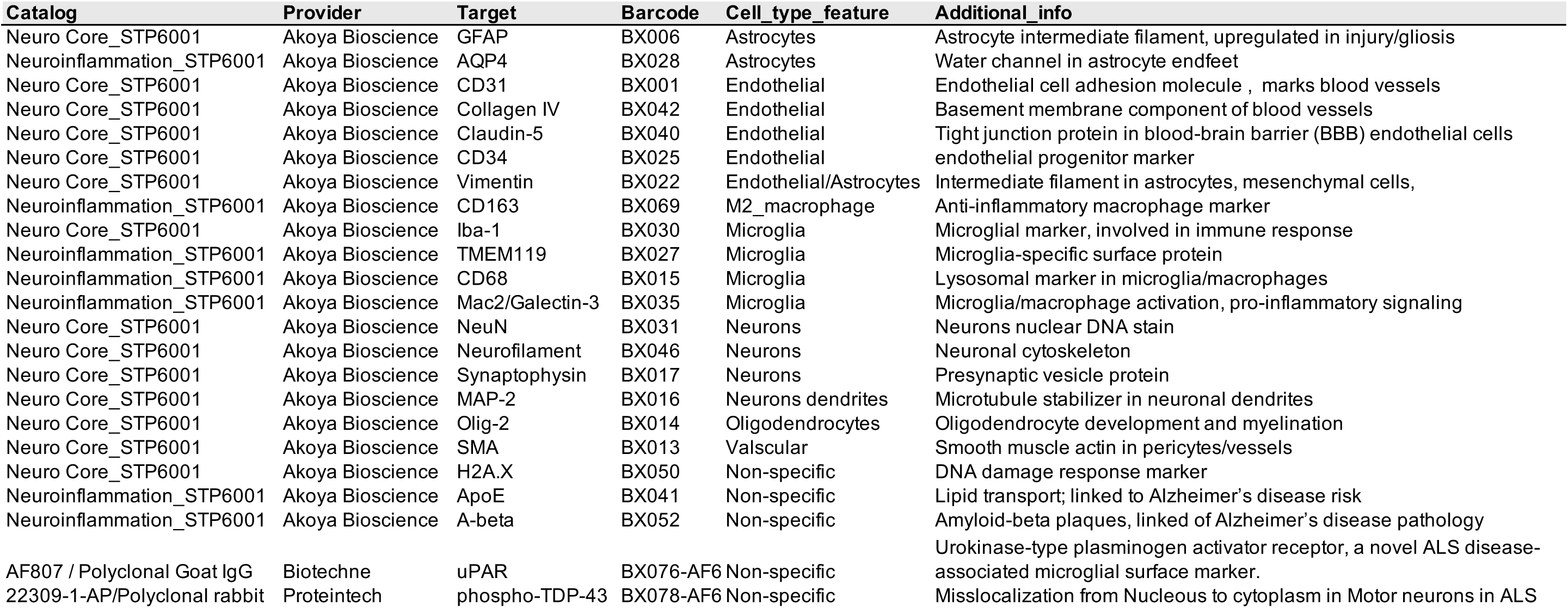
Antibodies used for Akoya multiplex imaging.

**Table 4.**
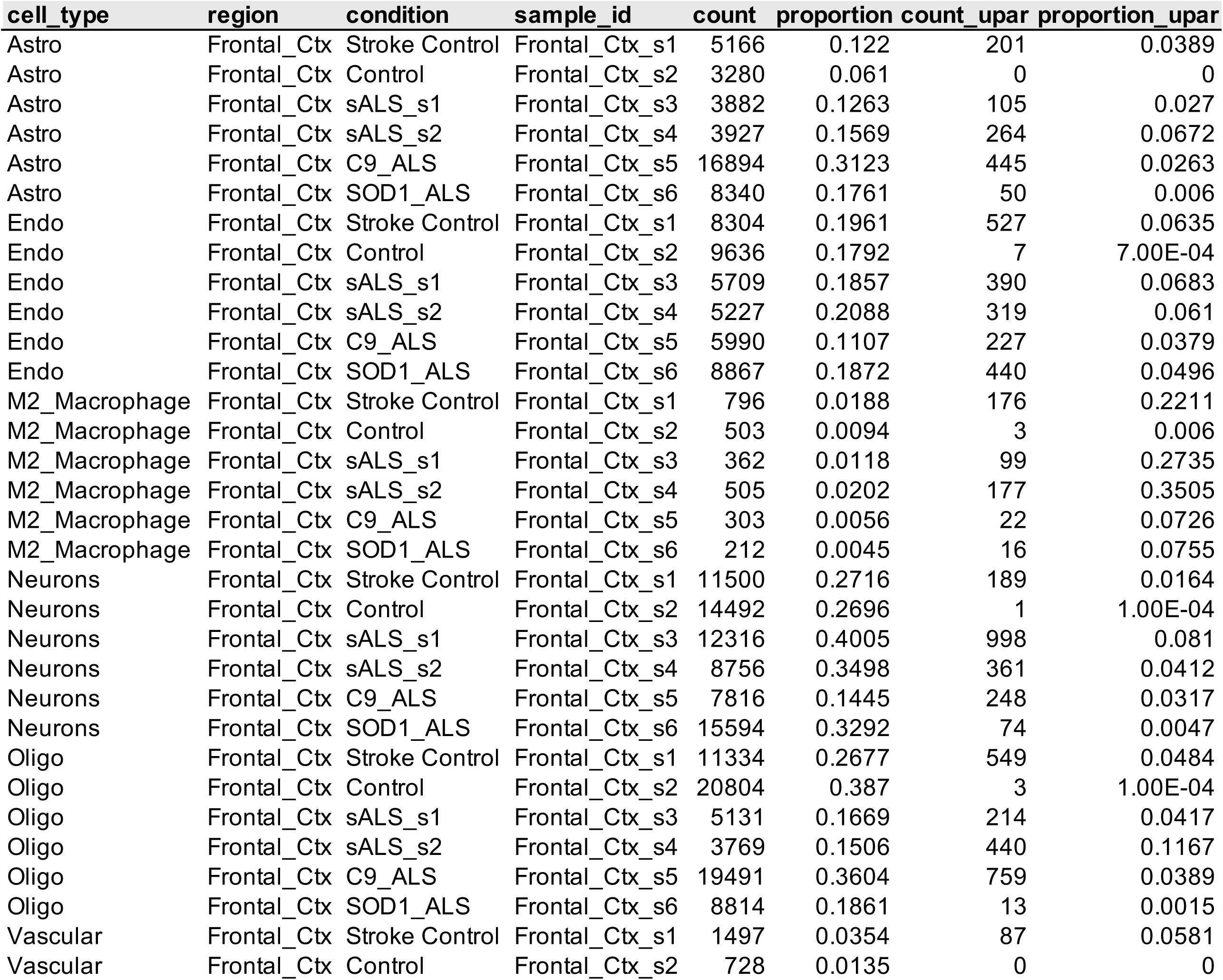
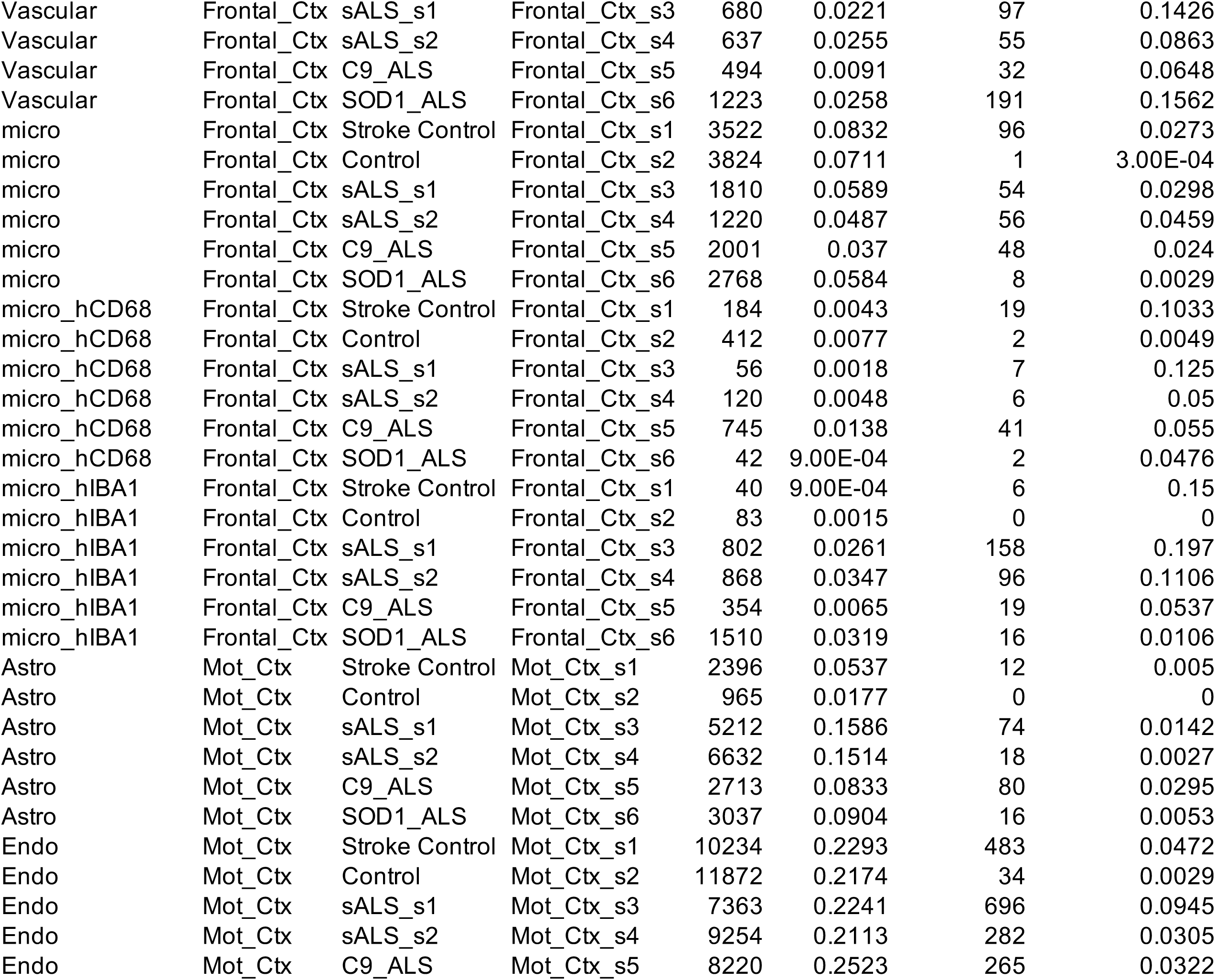
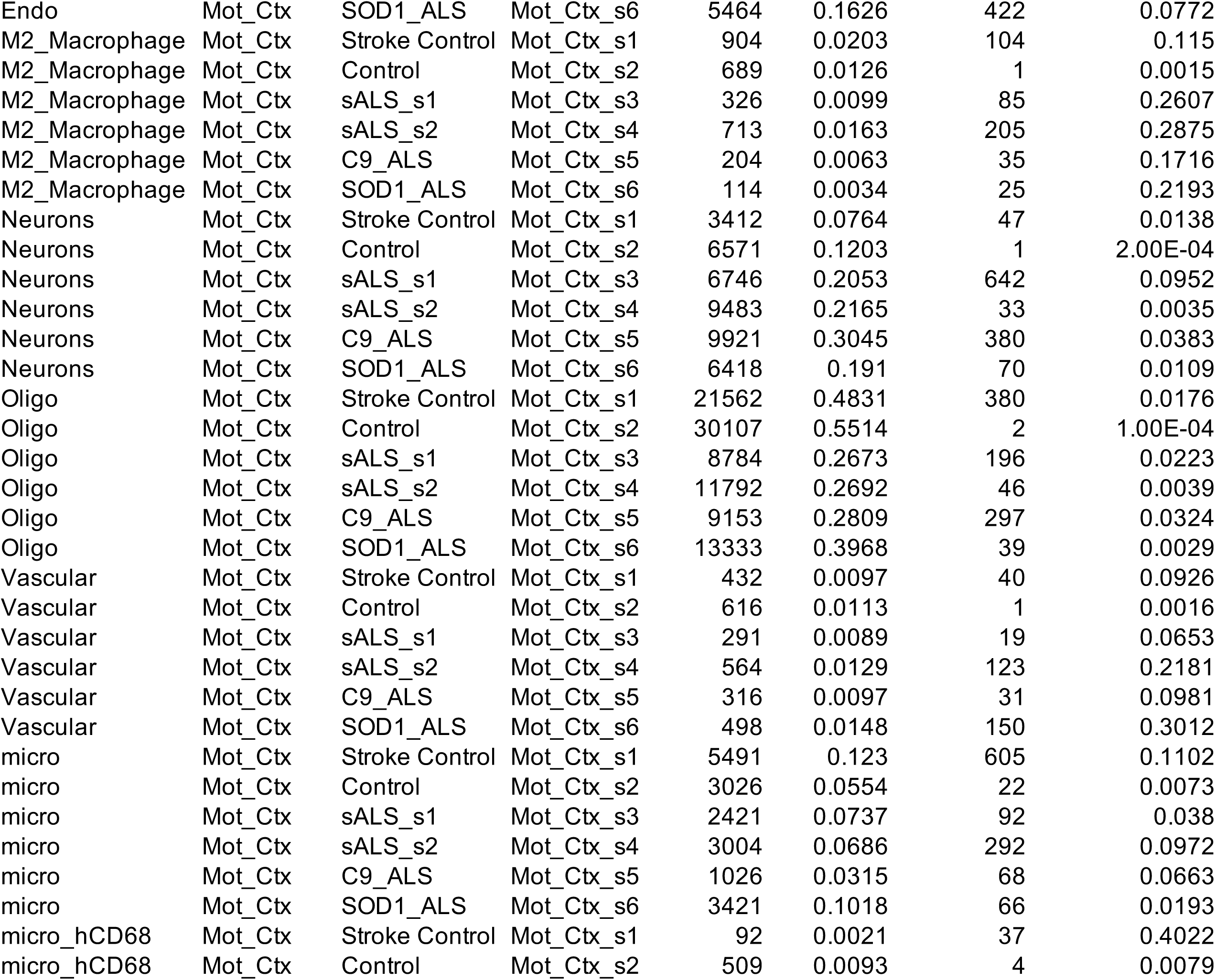
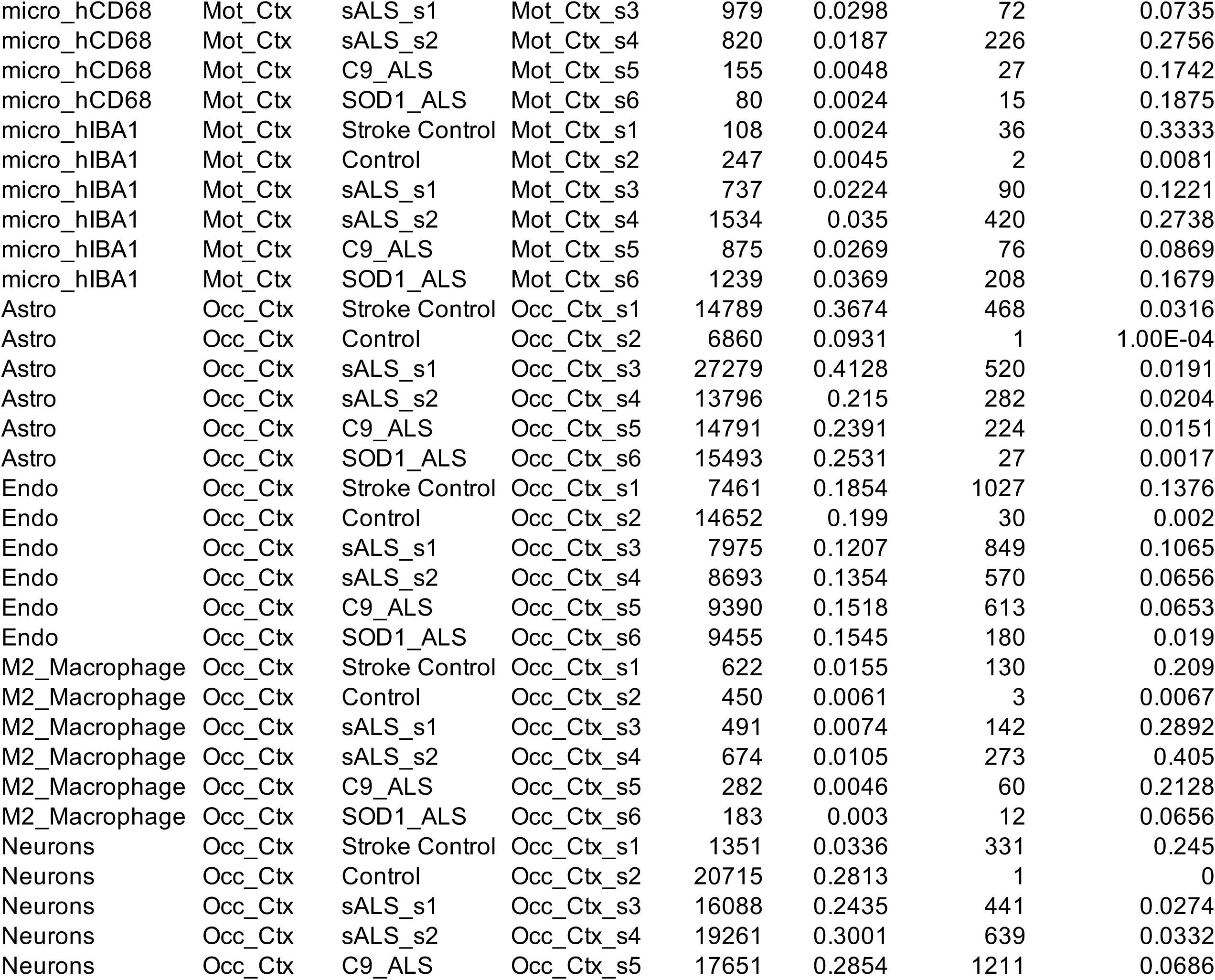
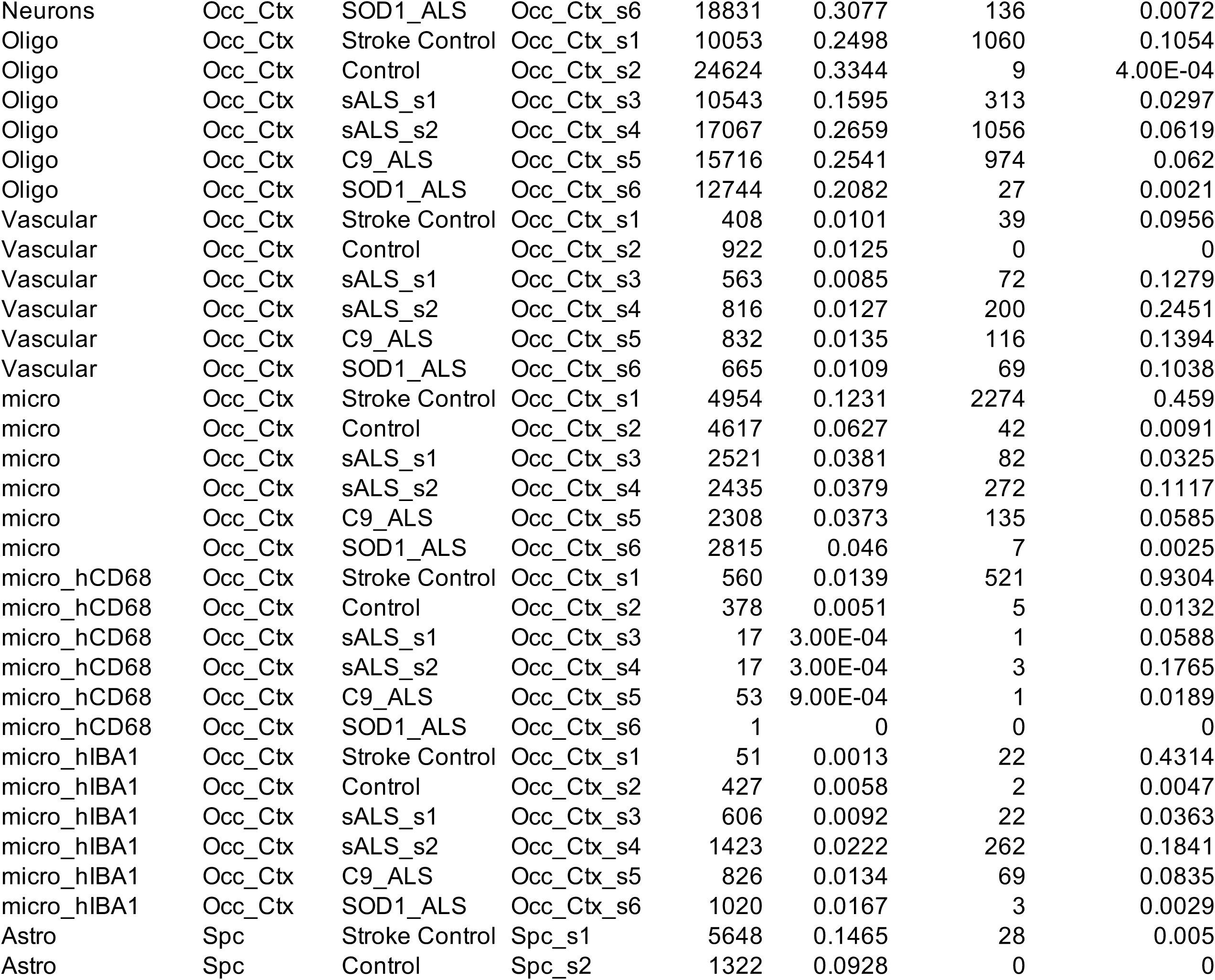
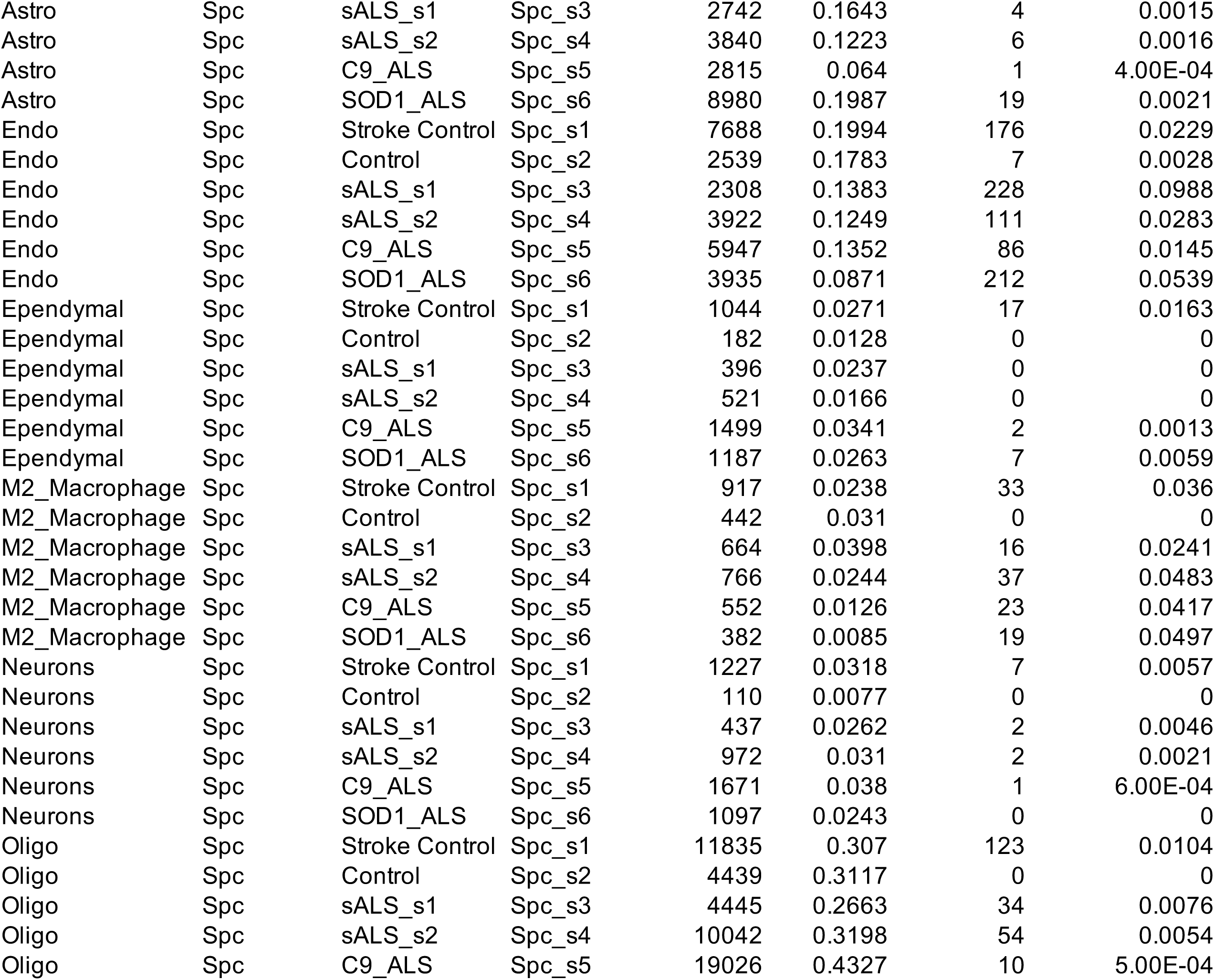
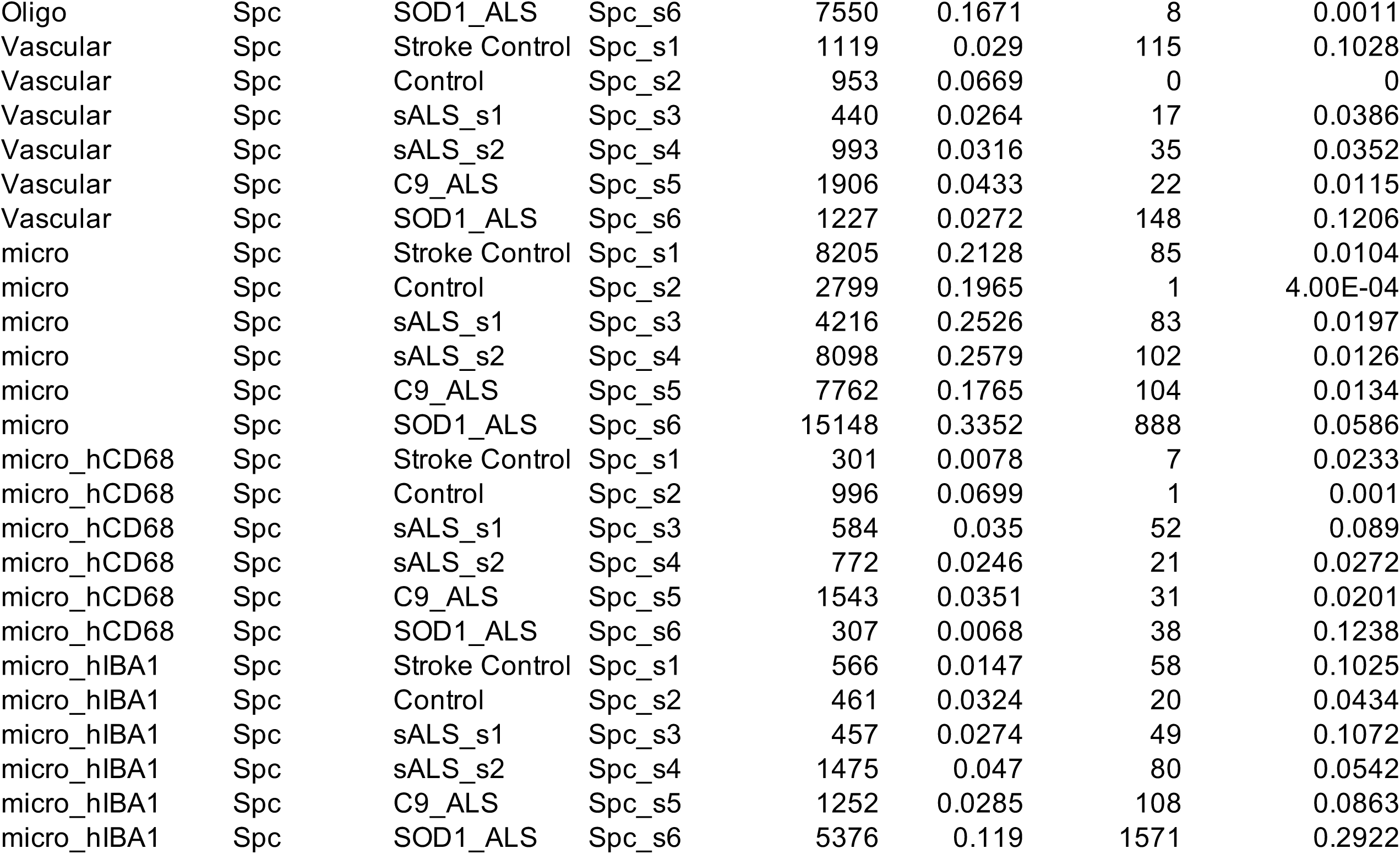
Multiplex analysis cell proportions.

**Fig. S1. RNAseq from Post-Mortem Tissues Reveals Increased Expression of uPAR in ALS Cases. (A)** Floating box showing uPAR expression in cerebellum (CB), frontal cortex (FCX), temporal cortex (TCX), cervical spinal cord (CSC), and lumbar spinal cord (LSC) in post-mortem tissues from ALS, ALS/other neurodegenerative diseases (ALS cases with co-morbidities, e.g., ALS/FTD), and control cases. Bulk RNA-seq data were analyzed using reference-based cell-aware deconvolution, and TOAST-derived, composition-adjusted normalized expression values were used for comparisons. **(B)** UMAP performed on single cells RNAseq from post-mortem tissues from FCX and MCX. **(C)** Analysis of spinal cord PLAUR RNA levels in control and SOD1^G93A^ mice at indicated time points (*p<0.05) **(D)** Quantification of PLAUR RNA levels in microglia isolated from the cortex and spinal cord of TDP-43 (rNLS8) mice across different disease stages.

**Fig. S2. uPAR Is Elevated in ALS-Affected Tissues. (A)** Western blot for uPAR (∼50 kDa) in occipital cortex homogenates from control, C9-ALS, and sporadic ALS cases. **(B)** Quantification of uPAR levels normalized to total protein and expressed as fold change relative to control. (**C**) Western blot for uPAR (∼50 kDa) in spinal cord homogenates from control, C9-ALS, SOD1-ALS, and sporadic ALS cases. **(D)** Quantification of uPAR levels normalized to total protein and expressed as fold change relative to control. Data shown as mean ± S.E.M (*p<0.05, **p<0.01, ***p<0.001). Symbols are biological replicates (Ctrl, n = 5; C9-ALS/FTD, n = 9; SOD1-ALS, n = 3; sALS, n = 4). **(E)** Representative sections of mid motor cortex were stained for astrocytes (GFAP, Alexa 546, red), uPAR (CD87, Alexa 647, magenta), and nuclei (DAPI, blue). Scale bars = 20 µm. **(F)** The percentage of GFAP⁺ cells that were also uPAR⁺ was calculated from three non-overlapping regions of interest per donor. Data are represented as mean ± S.E.M.; symbols are biological replicates (Ctrl, n = 4; C9-ALS/FTD, n = 3; sALS, n = 3). One-way ANOVA with Dunnett post-hoc. **The** plot shows the distribution of uPAR mean fluorescence intensity (MFI) within individual GFAP⁺ astrocytes. One-way ANOVA with Tukey post-hoc: non-significant.

**Fig. S3. Single-Cell Clustering and Annotation of the Spatial Proteomics Dataset.** Analysis for (**A**) motor cortex, (**B**) spinal cord, (**C**) frontal cortex, and (**D**) occipital cortex. The number of recovered cells for spatial analysis ranged from approximately 8,000 to 15,000 in the spinal cord, 40,000 to 85,000 in the occipital cortex, and 32,000 to 55,000 in the motor and frontal cortex. For each tissue, the left panel shows a heatmap of the mean scaled expression of canonical lineage-defining markers across the identified clusters. The middle panel displays a UMAP projection of all single cells, colored by annotated cell type. The right panel shows the spatial scatter plot (catplot) of cells mapped to their original tissue coordinates and the number of cells in each sample.

**Fig. S4.** (**A**) Spatial scatter plot (catplot) of cells, with tissue anatomy outlined (dark green: white matter, light green: grey matter). Red points represent the combined uPAR^+^ microglia population (micro, micro_CD68^+^, and micro_IBA1^+^). (**B**) Field of view from the SOD1-ALS spinal cord visualized in TissUUmaps. Individual channels for DAPI (nuclei), Iba1 (microglia), uPAR, and CD68 are shown. Red points are overlaid to indicate the spatial coordinates of uPAR+ microglia cells.

**Fig. S5. Mac2/Galectin-3 expression in the SOD1-ALS spinal cord.** (**A**) Violin plots showing the distribution of Mac2/Galectin-3 expression levels across all annotated cell types. (**B-C**) Representative multiplexed images from the SOD1-ALS spinal cord, acquired in QuPath (v0.6.0), displaying the same field of view in the (B) grey matter and (**C**) white matter. For each region, individual channels for Iba1 (microglia), Mac2/Galectin-3, and uPAR are shown.

**Fig. S6. Cellular Neighborhood (CN) analysis of the motor cortex. (A)** Elbow plot with the distortion score used to determine the optimal number of neighborhoods (k). **(B)** Heatmap showing cell type enrichment within CNs. The log2 fold-change values indicate enrichment (positive) and depletion (negative) of cell types that define the unique composition of each neighborhood. **(C)** Spatial distribution of CNs across each condition. Each point represents a single cell colored by its assigned CN. **(D, E)** Spatial context maps showing CN frequency and co-occurrence per condition. Rows represent the number of CN combinations within a local window: the first row shows windows dominated by a single CN; subsequent rows show windows where multiple CNs co-occur. The size of the dots represents the frequency of each CN and connecting asterisks (*) indicate a high frequency of spatial proximity between pairs of CNs.

**Fig. S7. CN analysis of the spinal cord. (A)** Elbow plot with the distortion score used to determine the optimal number of neighborhoods (k). **(B)** Heatmap showing cell type enrichment within CNs. The log2 fold-change values indicate enrichment (positive) and depletion (negative) of cell types that define the unique composition of each neighborhood. **(C)** Spatial distribution of CNs across each condition. Each point represents a single cell colored by its assigned CN. **(D, E)** Spatial context maps showing CN frequency and co-occurrence per condition. Rows represent the number of CN combinations within a local window: the first row shows windows dominated by a single CN; subsequent rows show windows where multiple CNs co-occur. The size of the dots represents the frequency of each CN and connecting asterisks (*) indicate a high frequency of spatial proximity between pairs of CNs.

**Fig. S8. CN analysis of the frontal cortex. (A)** Elbow plot with the distortion score used to determine the optimal number of neighborhoods (k). **(B)** Heatmap showing cell type enrichment within CNs. The log2 fold-change values indicate enrichment (positive) and depletion (negative) of cell types that define the unique composition of each neighborhood. **(C)** Spatial distribution of CNs across each condition. Each point is a single cell colored by its assigned CN. **(D, E)** Spatial context maps showing CN frequency and co-occurrence per condition. Rows represent the number of CN combinations within a local window: the first row shows windows dominated by a single CN; subsequent rows show windows where multiple CNs co-occur. The size of the dots represents the frequency of each CN and connecting asterisks (*) indicate a high frequency of spatial proximity between pairs of CNs.

**Fig. S9. CN analysis of the occipital cortex. (A)** Elbow plot with the distortion score used to determine the optimal number of neighborhoods (k). **(B)** Heatmap showing cell type enrichment within CNs. The log2 fold-change values indicate enrichment (positive) and depletion (negative) of cell types that define the unique composition of each neighborhood. **(C)** Spatial distribution of CNs across each condition. Each point represents a single cell colored by its assigned CN. **(D, E)** Spatial context maps showing CN frequency and co-occurrence per condition. Rows represent the number of CN combinations within a local window: the first row shows windows dominated by a single CN; subsequent rows show windows where multiple CNs co-occur. The size of the dots represents the frequency of each CN and connecting asterisks (*) indicate a high frequency of spatial proximity between pairs of CNs.

**Fig. S10. Identification of Compounds Enhancing uPAR Expression in C20 Cells**. (A) Histograms depicting fluorescence intensities of uPAR across treatments. (B) Quantification of the percentage of uPAR⁺ C20 cells across tested conditions. Data represent three technical replicates from three independent experiments. Values are presented as mean ± S.E.M. (*p<0.05, **p<0.01, ***p<0.001). Quantification of the MFI of uPAR⁺ C20 cells across tested conditions. Data represent three technical replicates from three independent experiments. Values are presented as mean ± S.E.M. (*p<0.05, **p<0.01, ***p<0.001). (D) Representative confocal images of pNF-κB S536 in C20 across treatments (Actin is in green, pNF-κB S536 is in fire pseudo color) (E) Quantification of pNF-κB S536 fluorescence intensity per cell. Data represent two technical replicates from one independent experiment. Values are presented as mean ± S.E.M.

**Fig. S11. Flow Cytometry Gating Strategies. (A)** Gating strategy used for flow cytometric analysis in Figure 3. **(B)** Gating strategy used for flow cytometric analysis in Figure 5.

**Fig. S12. Image Processing for Neurite Analysis. (A)** Workflow representation of image segmentation and neurite tracing used in neuronal assays. **(B-G)** Bar plot quantifying neurite lengths in the six different lines tested under the different condition indicated

**Fig. S13. Phenotypic Characterization of uPAR-CR T Cells. (A)** Flow cytometry plots showing CD3-BUV805 vs. GFP in untransduced and uPAR CAR-T cells to assess transduction efficiency. **(B)** Flow cytometry plots showing CD8-BB700 vs. CD4-BV421 to determine T cell subtype composition. **(C)** Bar graph quantifies the proportions of CD4⁺, CD8⁺, double-positive, and double-negative T cells in untransduced and uPAR CAR-T populations.

**Fig. S14. Validation of uPAR-Specific Killing Using FITC-CAR-T Cells. (A)** Flow cytometry plots showing uPAR (CD87-PE) intensity vs. FSC in C20 cells treated with FITC-uPAR alone or in combination with FITC-CAR-T cells. **(B)** Histogram showing uPAR⁺ cell counts under FITC-uPAR alone or FITC-uPAR + FITC-CAR-T cell conditions. **(C)** Bar graph quantifies the percentage of uPAR⁺ C20 cells under indicated conditions. Data represent mean ± S.E.M. (*p<0.05, **p<0.01, ***p<0.001). **(D–F)** Bar graphs showing the percentage of live cells **(D)**, uPAR⁺ cells **(E)**, and CD11b⁺ cells **(F)** in C20 cultures treated with FITC-uPAR alone or in combination with untransduced T cells. Data are from three replicates, shown as mean ± S.E.M.

**Fig. S15. Pathological Mislocalization of pTDP43 in ALS Motor Neurons.** To assess ALS-associated protein pathology, we evaluated the localization of phosphorylated TDP43 (pTDP43). Consistent with the hallmark of ALS, we observed pathological mislocalization of pTDP43 from the nucleus to the cytoplasm of motor neurons in ALS cases. This phenotype was most notable in sporadic ALS (sALS) and C9ORF72-mutation (C9_ALS) cases, as shown in **Supplementary Figure 1**. Representative images of spinal cord motor neurons across conditions, with two neurons shown per condition (**A-B**). Images were acquired using QPath (v0.6.0). For each field, a merged channel view of MAP-2 (neuronal cell body, green) and NeuN (neuronal marker, red) or pTDP43 (purple) is displayed. The scale bar is located at the bottom of all images.

**Fig. S16. Cell Segmentation and Quality Control.** Representative images from the motor cortex display the segmentation results for two randomly selected fields of view. For each field, the left panel displays the raw multiplexed image (DAPI in green, Iba1 in blue), and the right panel shows the corresponding predicted segmentation masks (white outlines) overlaid on the DAPI channel. The segmentation was performed using the MESMER deep learning model through SPACEc^59^.

**Fig. S17. Quality Control Filtering of Segmented Cells per Tissue.** For each of the four CNS tissues, the data quality is shown before and after filtering. The left panel for each tissue is a boxplot of cell area and DAPI intensity distributions across all samples. The two right panels are colored dot plots showing the relationship between cell area and DAPI intensity for all cells, colored by sample, before (middle) and after (right) applying a filter to remove the lowest 1% of cells by both metrics. This step removed potential debris and low-quality cells prior to downstream analysis.

**Fig. S18. Unsupervised clustering, cell type annotation; Quality control of cell type annotation by spatial overlay.** (**A**) Representative multiplexed images from a motor cortex sample (sALS_s2), visualized in QuPath (v0.6.0). Channels shown are GFAP (astrocytes, light blue), NeuN (neurons, green), Collagen IV (endothelial cells, purple), and Iba-1 (microglia, red), illustrating the spatial distribution of major cell lineages. (**B**) Spatial scatter plot (catplot) from TissUUmaps v3.2.1.11 of the same region, with each point representing a single cell colored by its identified annotation as astrocyte, neuron, endothelial, or microglia, validating the concordance between marker expression and spatial classification.

**Fig. S19. Distribution of uPAR expression.** Violin plots show the expression distribution of uPAR for each annotated cell type in (**A**) Motor Cortex, (**B**) Spinal Cord, (**C**) Frontal Cortex, and (**D**) Occipital Cortex. The dashed line in each plot indicates the mean uPAR expression level across all cell types for the respective tissue.

## Notes

https://www.ncbi.nlm.nih.gov/geo/query/acc.cgi?acc=GSE137810

https://www.ncbi.nlm.nih.gov/geo/query/acc.cgi?acc=GSE18597

https://www.ncbi.nlm.nih.gov/sra/?term=PRJNA624791

https://www.ncbi.nlm.nih.gov/geo/query/acc.cgi?acc=GSE219281

### Competing Interest Statement

The authors have declared no competing interest.

