## Supplementary Figures for "A CAR-T Cell-Based Strategy for Eliminating Pathogenic Microglia in ALS"

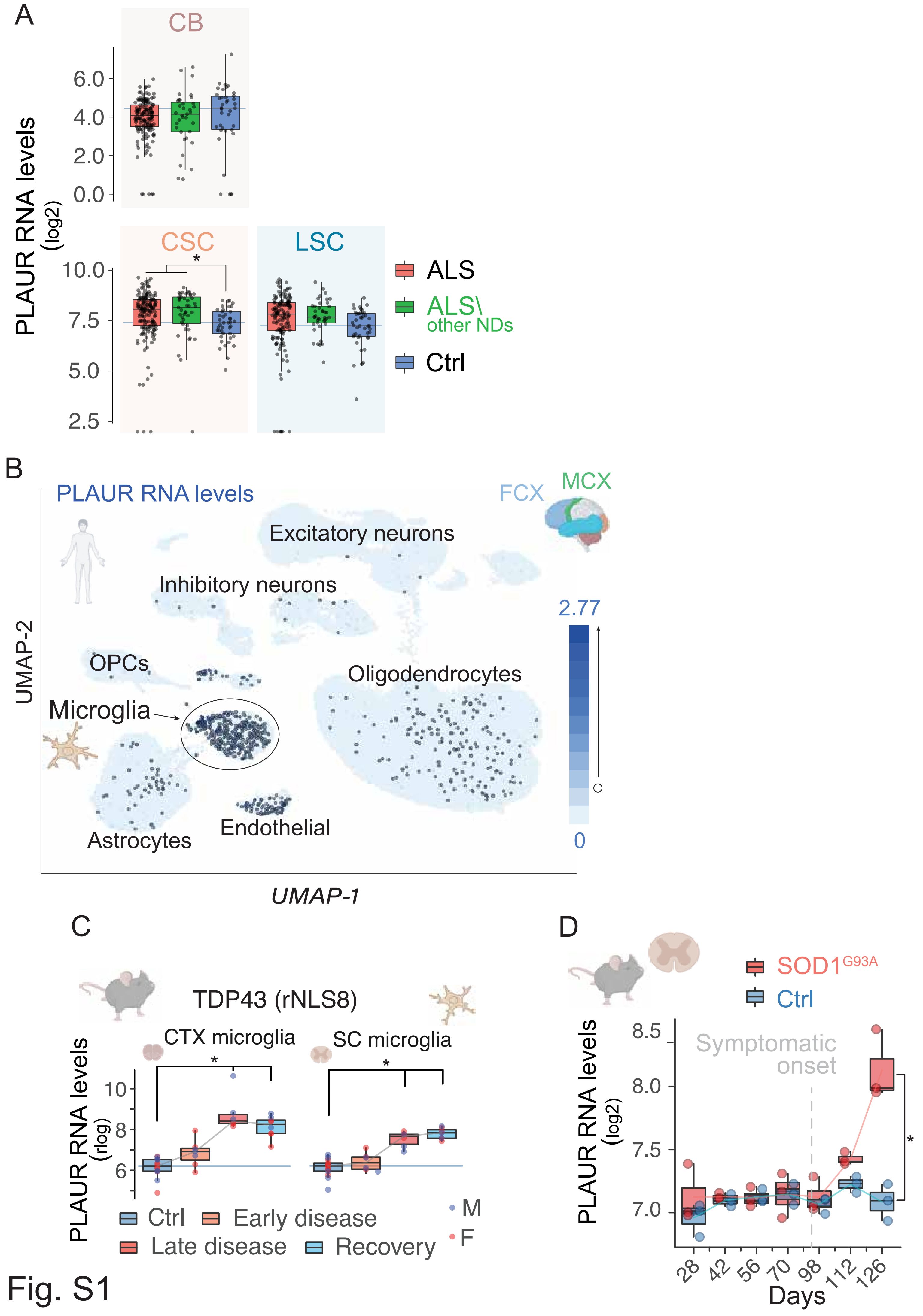
A
CB
6.0
4.0
2.0
0.0
PLAUR RNA levels (log2)
CSC
LSC
10.0
7.5
5.0
2.5
*
ALS
ALS\ other NDs
Ctrl
B
PLAUR RNA levels
FCX
MCX
Excitatory neurons
Inhibitory neurons
OPCs
Microglia
Oligodendrocytes
Astrocytes
Endothelial
2.77
0
UMAP-2
UMAP-1
C
TDP43 (rNLS8)
CTX microglia
SC microglia
PLAUR RNA levels (rlog)
10
8
6
Ctrl
Early disease
Late disease
Recovery
M
F
D
SOD1G93A
Ctrl
PLAUR RNA levels (log2)
8.5
8.0
7.5
7.0
Symptomatic onset
28
42
56
70
98
112
126
Days

Fig. S1

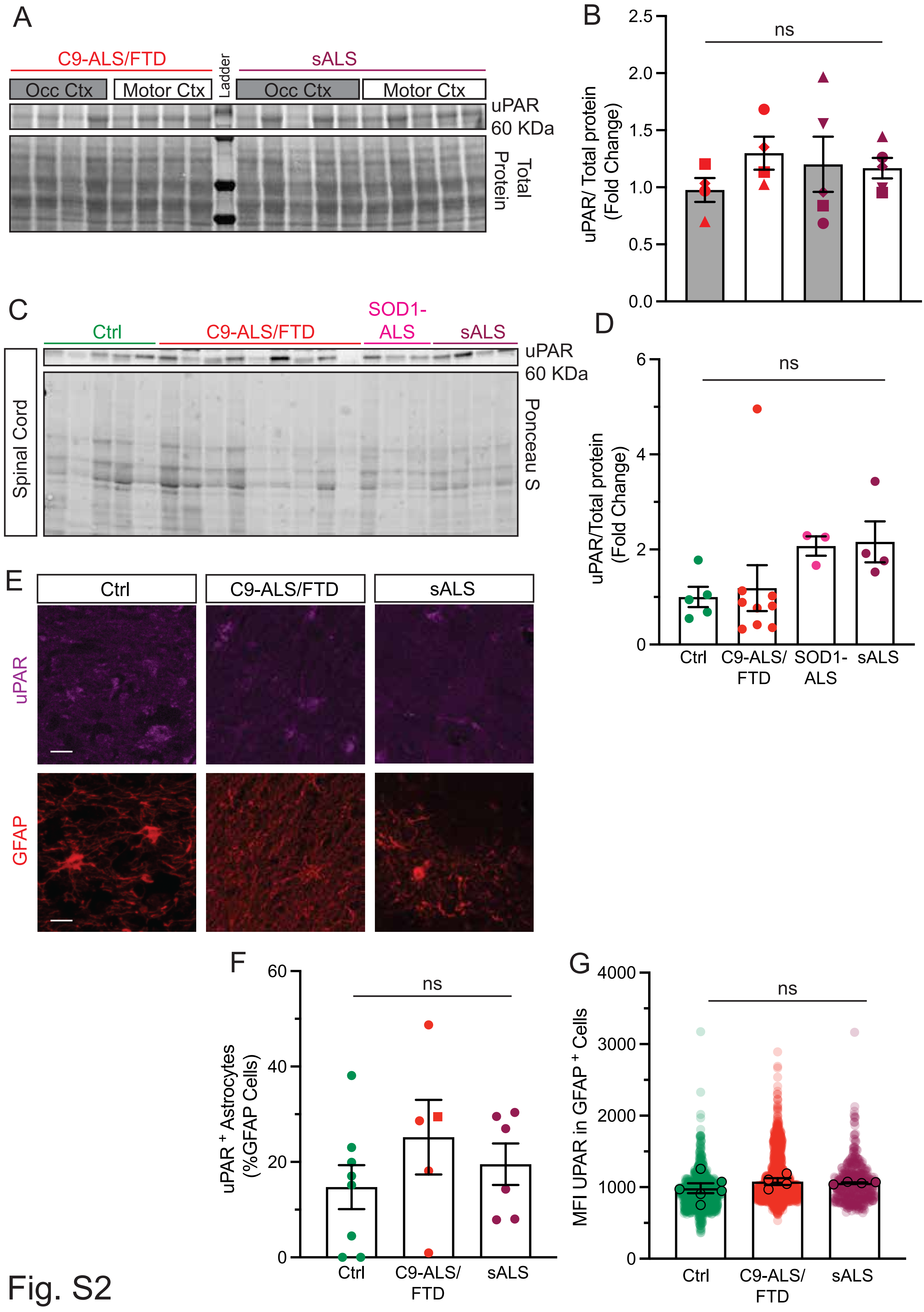

A
C9-ALS/FTD
sALS
Ladder
Occ Ctx
Motor Ctx
Occ Ctx
Motor Ctx
uPAR
60 KDa
Total Protein
B
ns
uPAR/ Total protein (Fold Change)
2.5
2.0
1.5
1.0
0.5
0.0
C
Ctrl
C9-ALS/FTD
SOD1-ALS
sALS
Spinal Cord
uPAR
60 KDa
Ponceau S
D
ns
uPAR/Total protein (Fold Change)
6
4
2
0
Ctrl
C9-ALS/ FTD
SOD1- ALS
sALS
E
Ctrl
C9-ALS/FTD
sALS
uPAR
GFAP
F
ns
uPAR+ Astrocytes (%GFAP Cells)
60
40
20
0
Ctrl
C9-ALS/ FTD
sALS
G
ns
MFI UPAR in GFAP+ Cells
4000
3000
2000
1000
0
Ctrl
C9-ALS/ FTD
sALS


Fig. S2

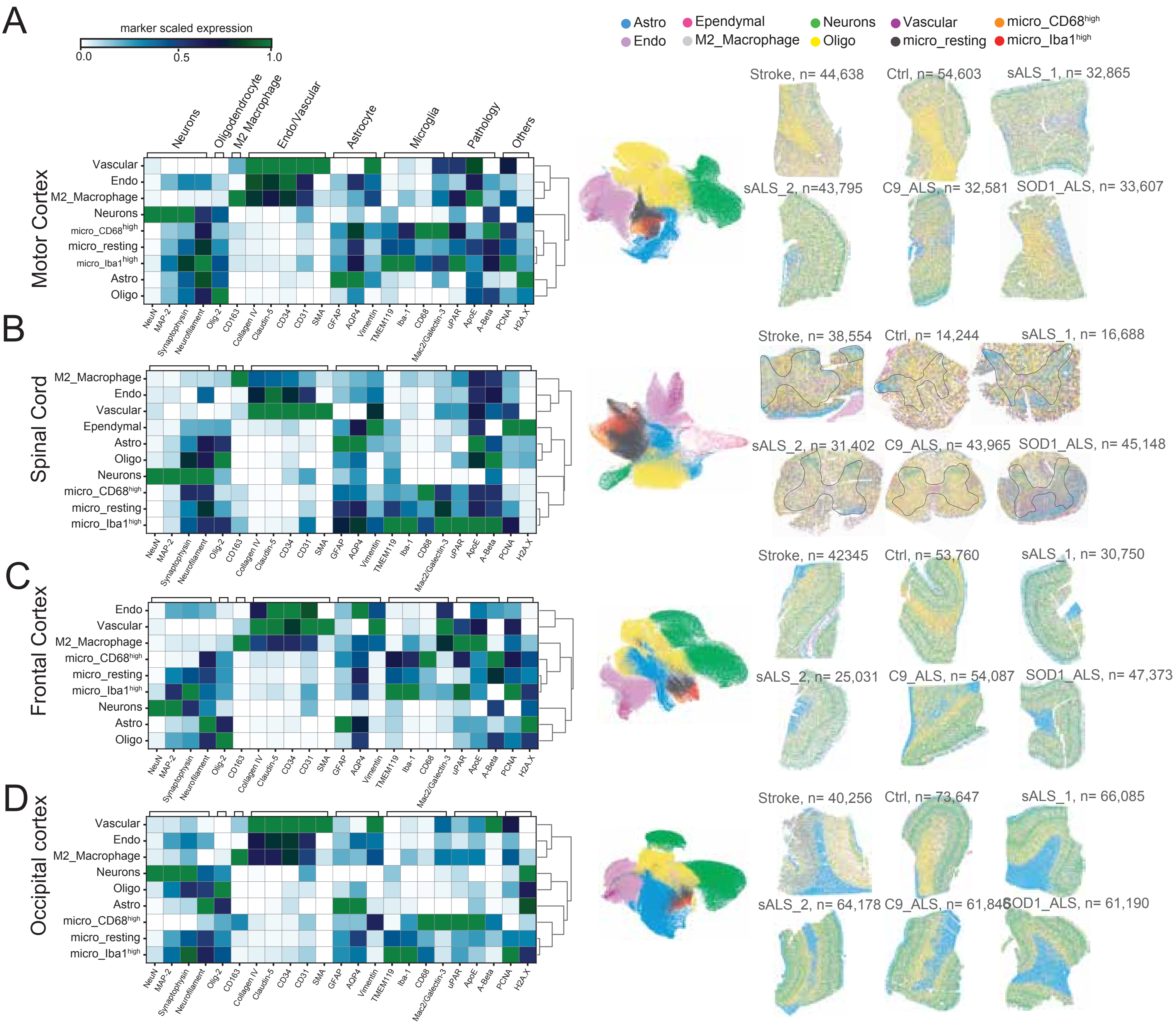
A
B
C
D
marker scaled expression
0.0
0.5
1.0
Neurons
Oligodendrocyte
M2 Macrophage
Endo/Vascular
Astrocyte
Microglia
Pathology
Others
Motor Cortex
Spinal Cord
Frontal Cortex
Occipital cortex
Vascular
Endo
M2_Macrophage
Neurons
micro_CD68high
micro_resting
micro_Iba1high
Astro
Oligo
Ependymal
NeuN
MAP-2
Synaptophysin
Neurofilament
Olig-2
CD163
Collagen IV
Claudin-5
CD34
CD31
SMA
GFAP
AQP4
Vimentin
TMEM119
Iba-1
CD68
Mac2/Galectin-3
uPAR
ApoE
A-Beta
PCNA
H2A.X
Astro
Ependymal
Neurons
Vascular
micro_CD68high
Endo
M2_Macrophage
Oligo
micro_resting
micro_Iba1high
Stroke, n= 44,638
Ctrl, n= 54,603
sALS_1, n= 32,865
sALS_2, n=43,795
C9_ALS, n= 32,581
SOD1_ALS, n= 33,607
Stroke, n= 38,554
Ctrl, n= 14,244
sALS_1, n= 16,688
sALS_2, n= 31,402
C9_ALS, n= 43,965
SOD1_ALS, n= 45,148
Stroke, n= 42345
Ctrl, n= 53,760
sALS_1, n= 30,750
sALS_2, n= 25,031
C9_ALS, n= 54,087
SOD1_ALS, n= 47,373
Stroke, n= 40,256
Ctrl, n= 73,647
sALS_1, n= 66,085
sALS_2, n= 64,178
C9_ALS, n= 61,848
SOD1_ALS, n= 61,190

Fig. S3

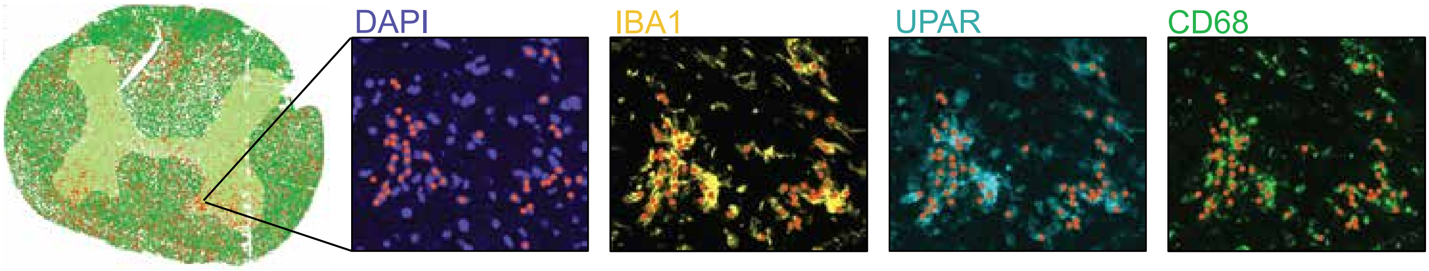


Fig. S4

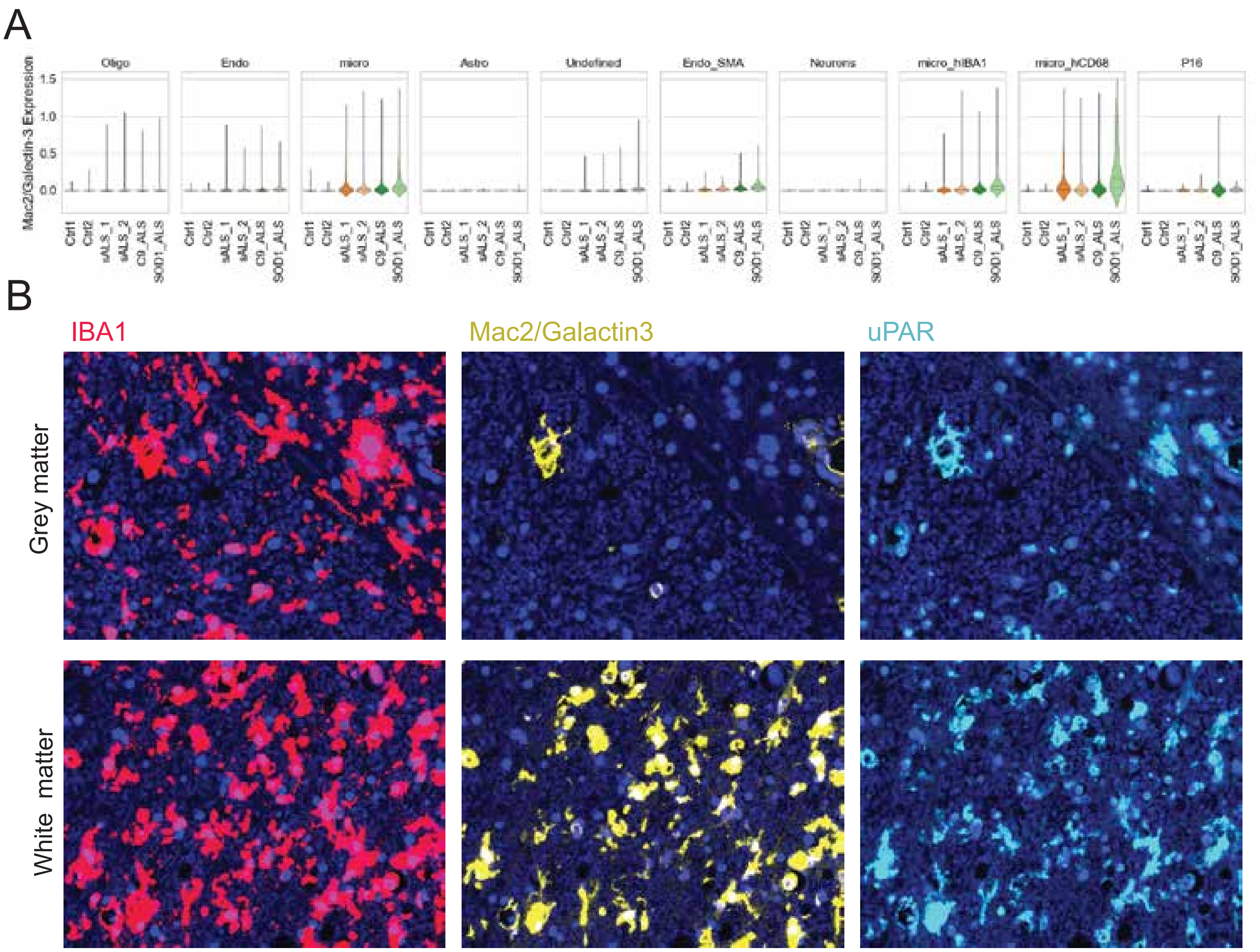

A
Mac2/Galectin-3 Expression
Oligo
Endo
micro
Astro
Undefined
Endo_SMA
Neurons
micro_hIBA1
micro_hCD68
P16
1.5
1.0
0.5
0.0
Ctrl1
Ctrl2
sALS_1
sALS_2
C9_ALS
SOD1_ALS
B
IBA1
Mac2/Galactin3
uPAR
Grey matter
White matter


Fig. S5

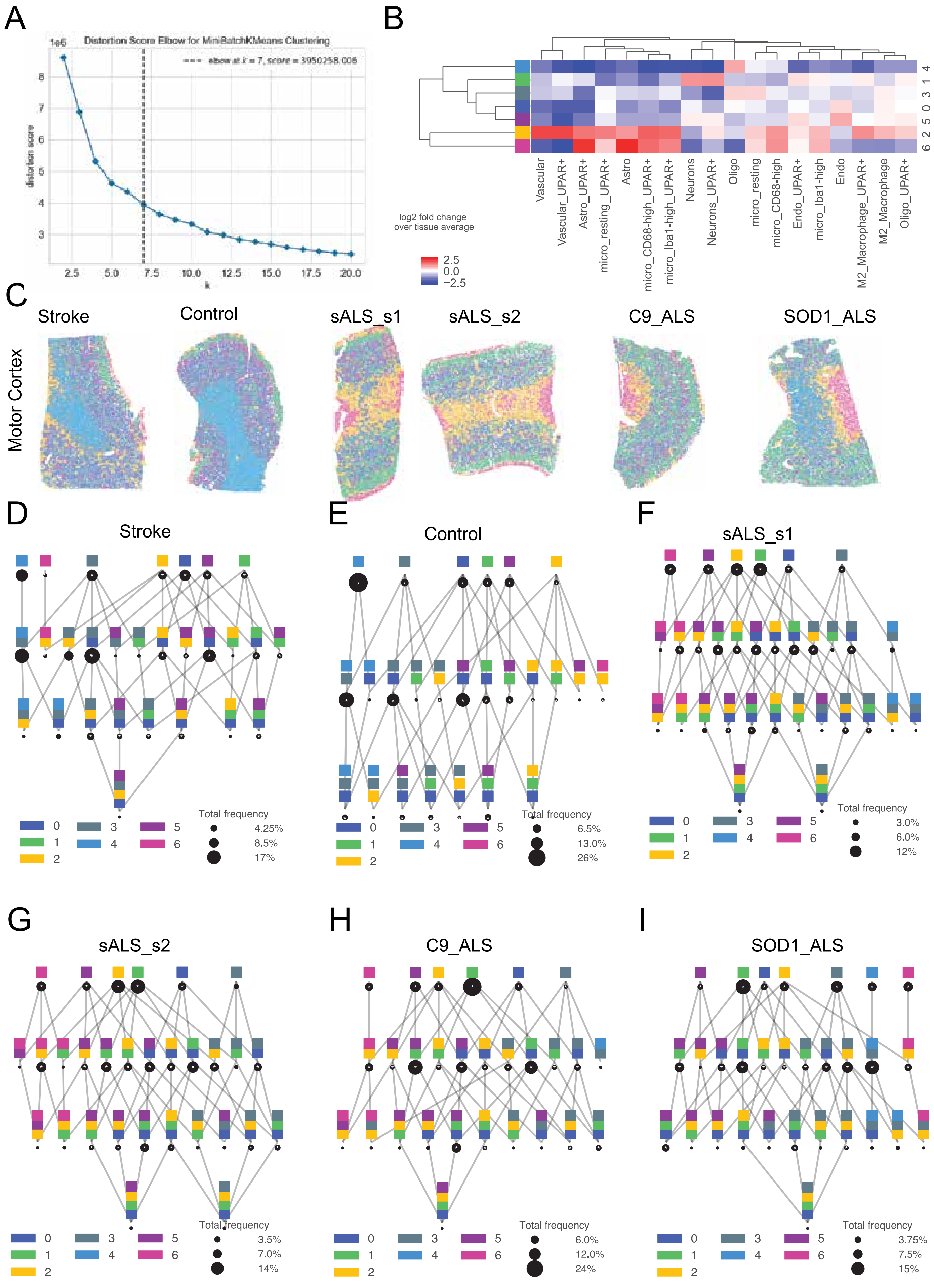
A
Distortion Score Elbow for MiniBatchKMeans Clustering
elbow at k = 7, score = 3950258.006
distortion score
k
B
log2 fold change over tissue average
Vascular
Vascular_UPAR+
Astro_UPAR+
micro_resting_UPAR+
Astro
micro_CD68-high_UPAR+
micro_Iba1-high_UPAR+
Neurons
Neurons_UPAR+
Oligo
micro_resting
micro_CD68-high
Endo_UPAR+
micro_Iba1-high
Endo
M2_Macrophage_UPAR+
M2_Macrophage
Oligo_UPAR+
C
Motor Cortex
Stroke
Control
sALS_s1
sALS_s2
C9_ALS
SOD1_ALS
D
Stroke
Total frequency
4.25%
8.5%
17%
E
Control
Total frequency
6.5%
13.0%
26%
F
sALS_s1
Total frequency
3.0%
6.0%
12%
G
sALS_s2
Total frequency
3.5%
7.0%
14%
H
C9_ALS
Total frequency
6.0%
12.0%
24%
I
SOD1_ALS
Total frequency
3.75%
7.5%
15%

Fig. S6

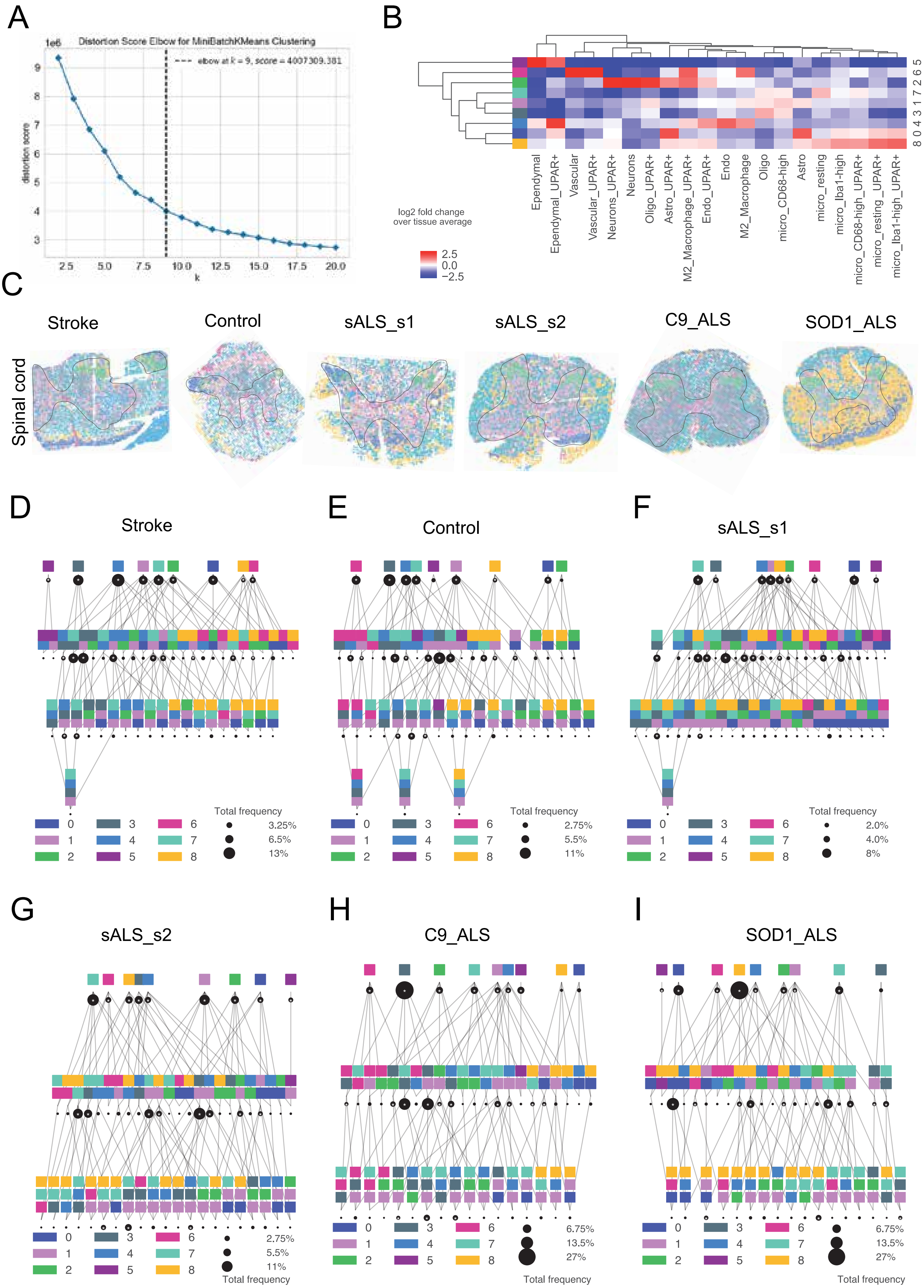
A
Distortion Score Elbow for MiniBatchKMeans Clustering
elbow at k = 9, score = 4007309.381
distortion score
k
B
log2 fold change over tissue average
Ependymal
Ependymal_UPAR+
Vascular
Vascular_UPAR+
Neurons_UPAR+
Neurons
Oligo_UPAR+
Astro_UPAR+
M2_Macrophage_UPAR+
Endo_UPAR+
Endo
M2_Macrophage
Oligo
micro_CD68-high
Astro
micro_resting
micro_Iba1-high
micro_CD68-high_UPAR+
micro_resting_UPAR+
micro_Iba1-high_UPAR+
C
Stroke
Control
sALS_s1
sALS_s2
C9_ALS
SOD1_ALS
Spinal cord
D
Stroke
Total frequency
3.25%
6.5%
13%
E
Control
Total frequency
2.75%
5.5%
11%
F
sALS_s1
Total frequency
2.0%
4.0%
8%
G
sALS_s2
2.75%
5.5%
11%
Total frequency
H
C9_ALS
6.75%
13.5%
27%
Total frequency
I
SOD1_ALS
6.75%
13.5%
27%
Total frequency

Fig. S7

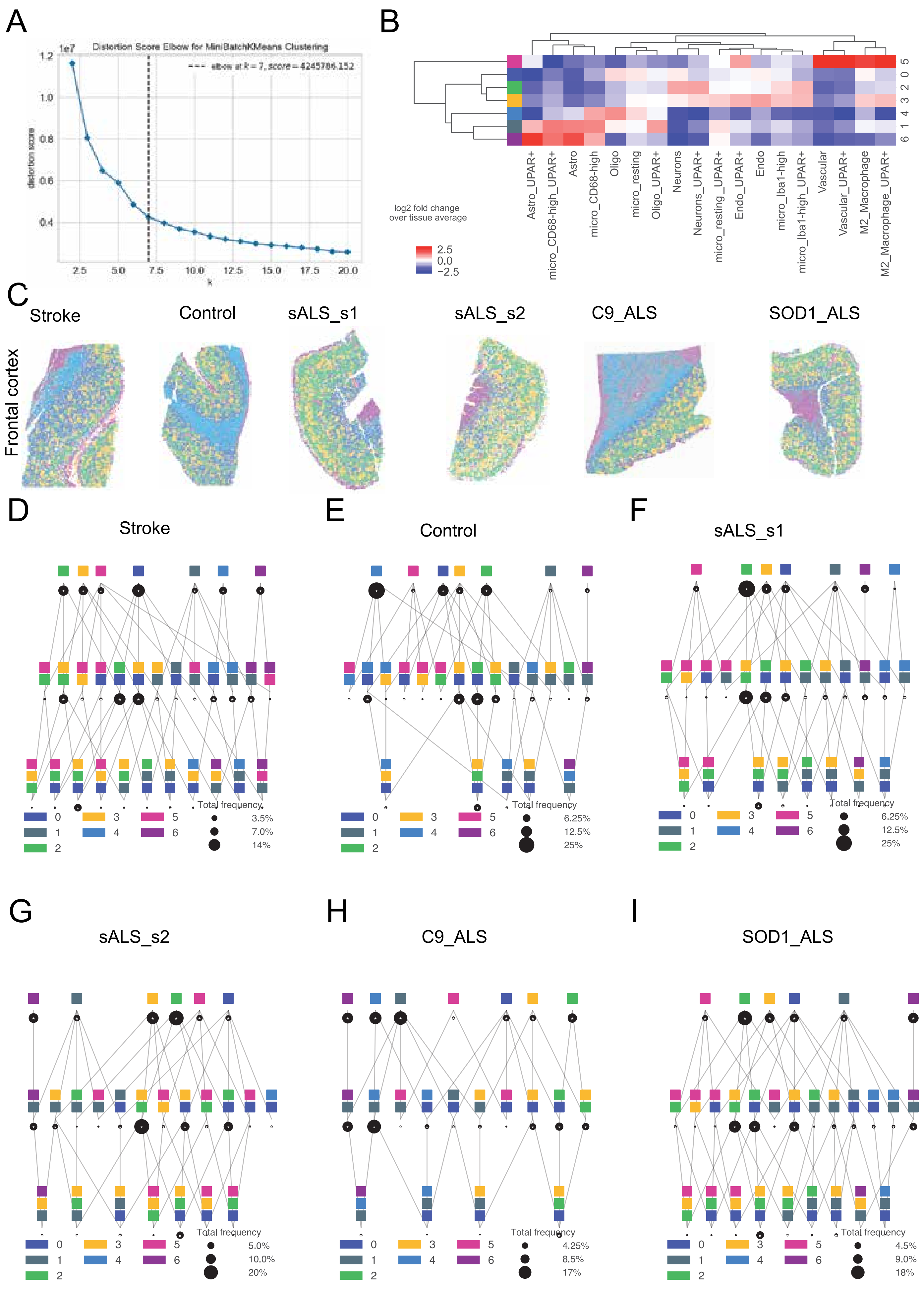
A
Distortion Score Elbow for MiniBatchKMeans Clustering
elbow at k = 7, score = 4245706.152
distortion score
k
B
log2 fold change over tissue average
Astro_UPAR+
micro_CD68-high_UPAR+
Astro
micro_CD68-high
Oligo
micro_resting
Oligo_UPAR+
Neurons
Neurons_UPAR+
micro_resting_UPAR+
Endo_UPAR+
Endo
micro_Iba1-high
micro_Iba1-high_UPAR+
Vascular
Vascular_UPAR+
M2_Macrophage
M2_Macrophage_UPAR+
C
Frontal cortex
Stroke
Control
sALS_s1
sALS_s2
C9_ALS
SOD1_ALS
D
Stroke
E
Control
F
sALS_s1
G
sALS_s2
H
C9_ALS
I
SOD1_ALS
Total frequency

Fig. S8

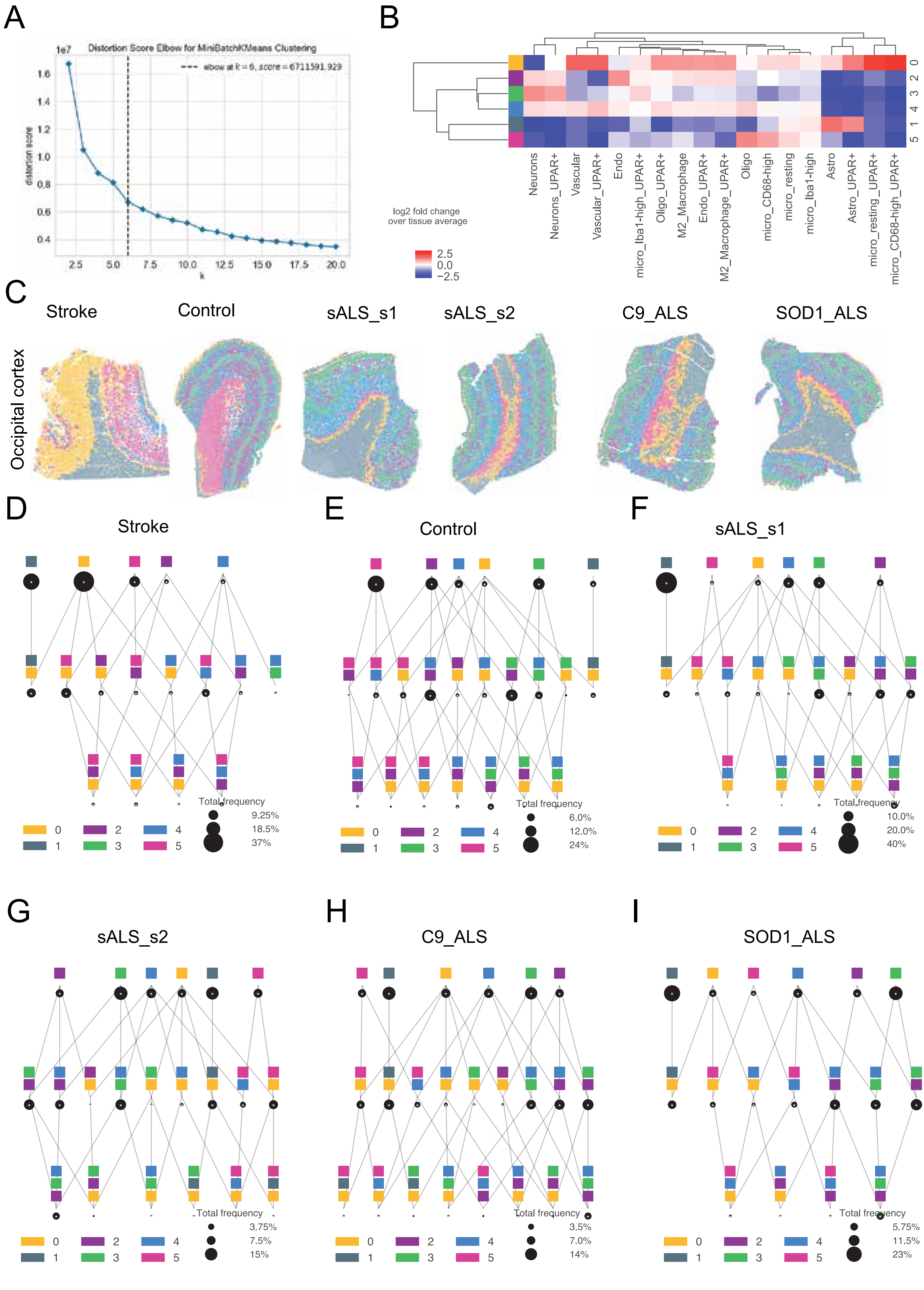
A
Distortion Score Elbow for MiniBatchKMeans Clustering
elbow at k = 6, score = 6711591.929
distortion score
k
B
log2 fold change over tissue average
Neurons
Neurons_UPAR+
Vascular
Vascular_UPAR+
Endo
micro_Iba1-high_UPAR+
Oligo_UPAR+
M2_Macrophage
Endo_UPAR+
M2_Macrophage_UPAR+
Oligo
micro_CD68-high
micro_resting
micro_Iba1-high
Astro
Astro_UPAR+
micro_resting_UPAR+
micro_CD68-high_UPAR+
C
Stroke
Control
sALS_s1
sALS_s2
C9_ALS
SOD1_ALS
Occipital cortex
D
Stroke
Total frequency
9.25%
18.5%
37%
E
Control
Total frequency
6.0%
12.0%
24%
F
sALS_s1
Total frequency
10.0%
20.0%
40%
G
sALS_s2
Total frequency
3.75%
7.5%
15%
H
C9_ALS
Total frequency
3.5%
7.0%
14%
I
SOD1_ALS
Total frequency
5.75%
11.5%
23%

Fig. S9

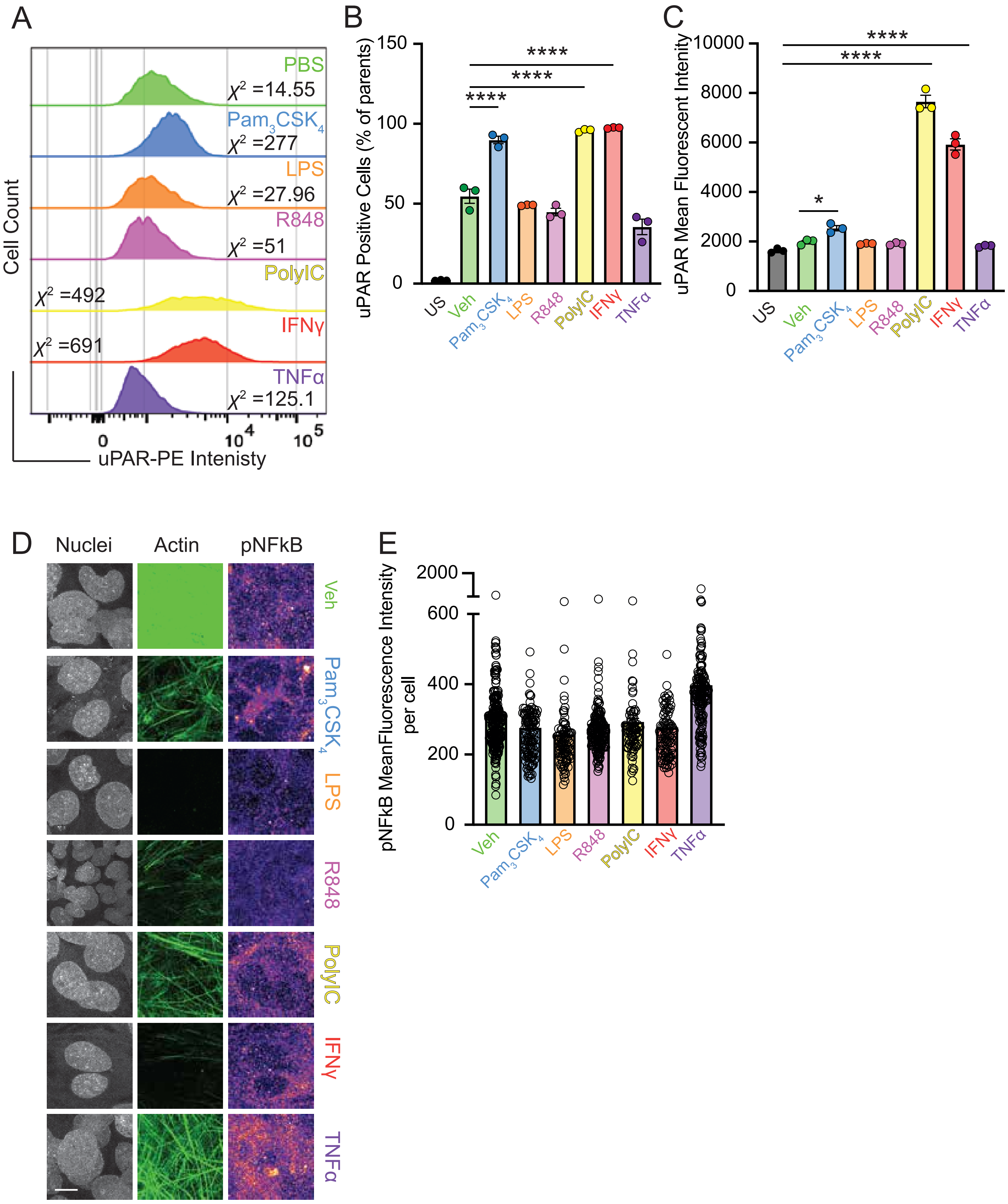

A
PBS
$X^2$ =14.55
$Pam_3CSK_4$
$X^2$ =277
LPS
$X^2$ =27.96
R848
$X^2$ =51
PolyIC
$X^2$ =492
IFNγ
$X^2$ =691
TNFα
$X^2$ =125.1
Cell Count
uPAR-PE Intenisty
B
uPAR Positive Cells (% of parents)
****
US
Veh
$Pam_3CSK_4$
LPS
R848
PolyIC
IFNγ
TNFα
C
uPAR Mean Fluorescent Intenity
*
D
Nuclei
Actin
pNFkB
E
pNFkB MeanFluorescence Intensity per cell


Fig. S10

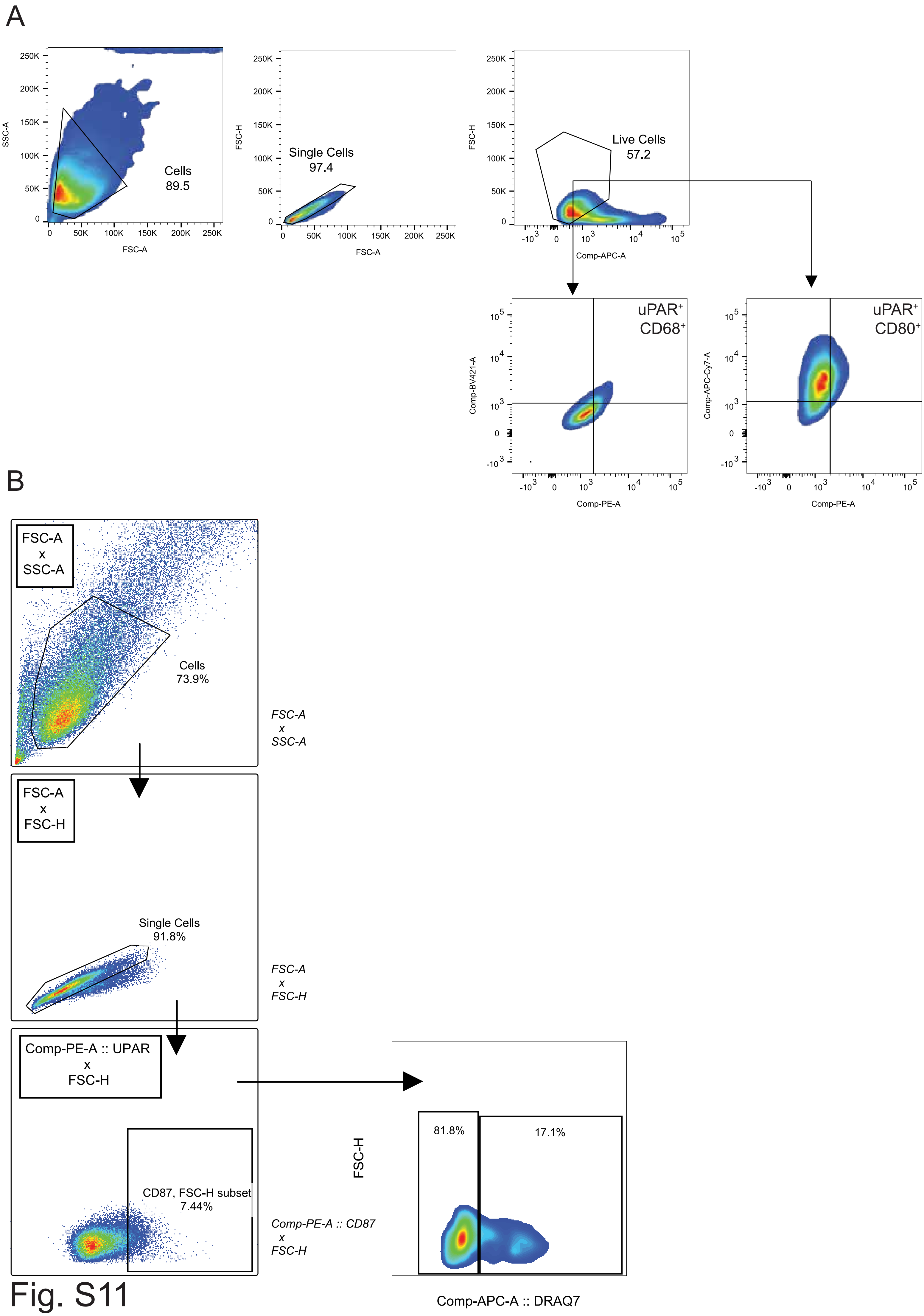

A
SSC-A
FSC-A
Cells
89.5
FSC-H
Single Cells
97.4
Live Cells
57.2
Comp-APC-A
Comp-BV421-A
Comp-PE-A
uPAR+
CD68+
Comp-APC-Cy7-A
uPAR+
CD80+
B
FSC-A x SSC-A
Cells
73.9%
FSC-A x SSC-A
FSC-A x FSC-H
Single Cells
91.8%
FSC-A x FSC-H
Comp-PE-A :: UPAR x FSC-H
CD87, FSC-H subset
7.44%
Comp-PE-A :: CD87 x FSC-H
FSC-H
81.8%
17.1%
Comp-APC-A :: DRAQ7


Fig. S11

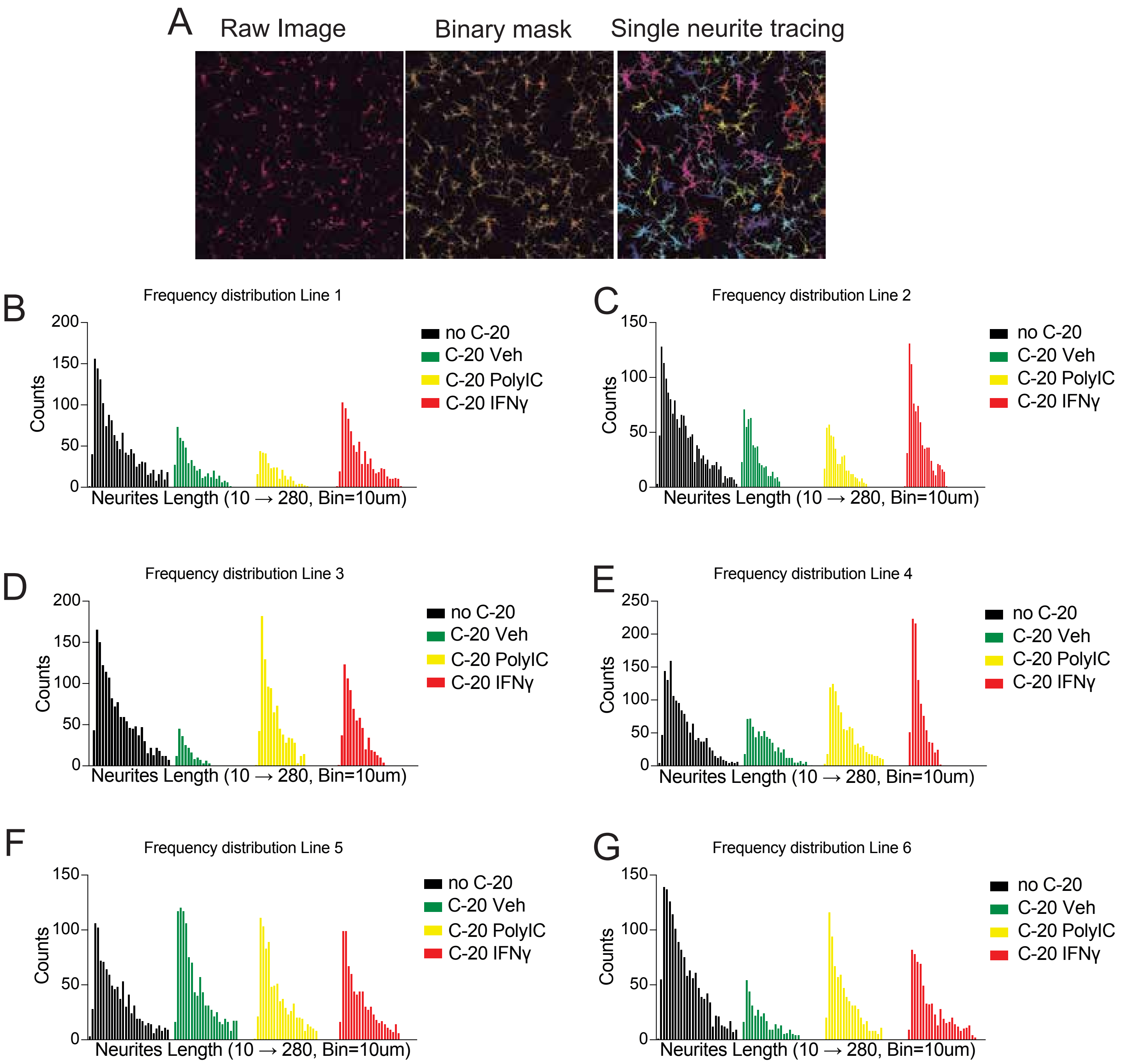
A
Raw Image
Binary mask
Single neurite tracing
B
Frequency distribution Line 1
C
Frequency distribution Line 2
D
Frequency distribution Line 3
E
Frequency distribution Line 4
F
Frequency distribution Line 5
G
Frequency distribution Line 6
Counts
Neurites Length (10 → 280, Bin=10um)
no C-20
C-20 Veh
C-20 PolyIC
C-20 IFNγ

Fig. S12

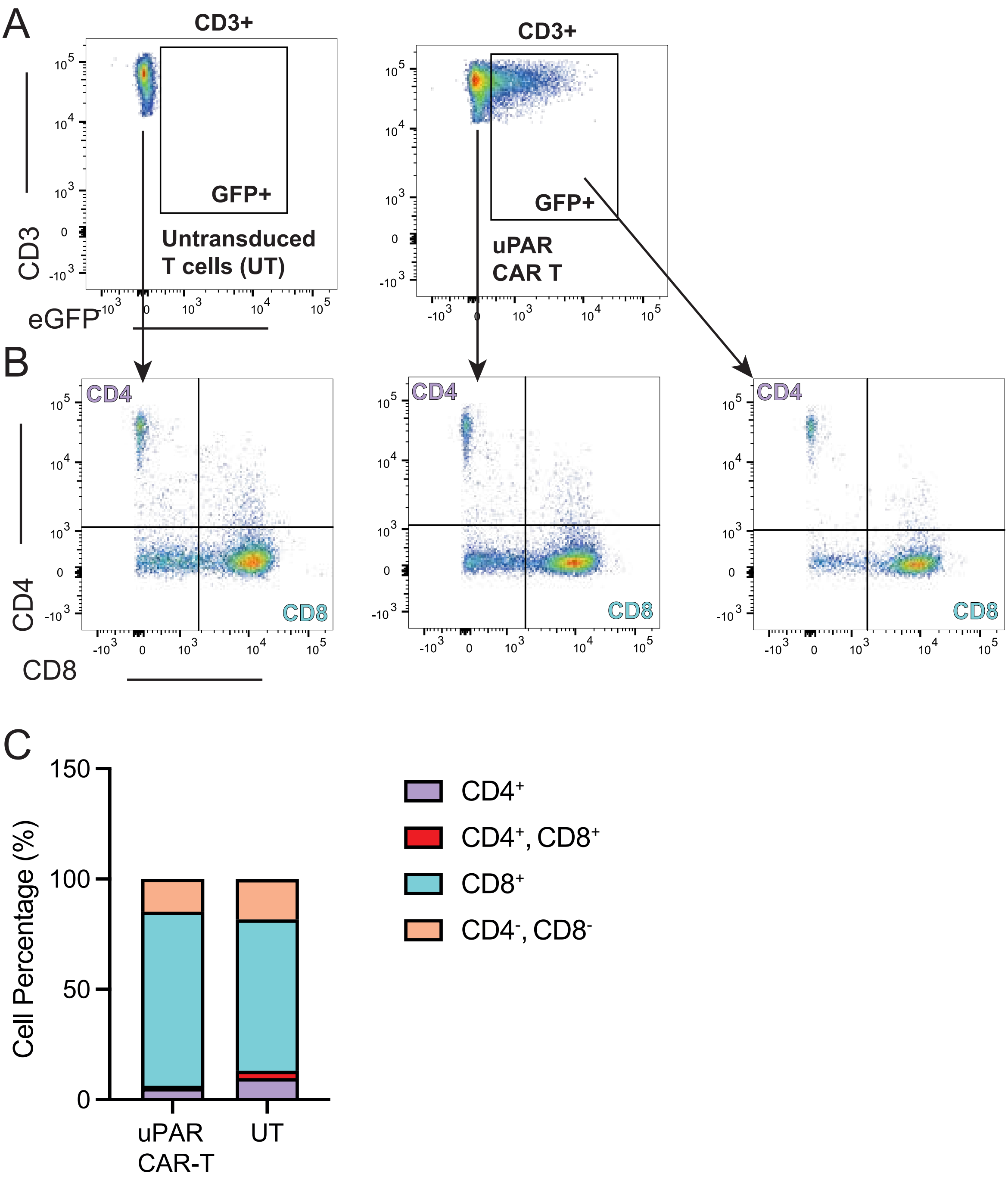
A
CD3+
GFP+
Untransduced
T cells (UT)
CD3+
GFP+
uPAR
CAR T
CD3
eGFP
B
CD4
CD8
CD4
CD8
CD4
CD8
CD4
CD8
C
150
100
50
0
Cell Percentage (%)
uPAR
CAR-T
UT
CD4⁺
CD4⁺, CD8⁺
CD8⁺
CD4⁻, CD8⁻

Fig. S13

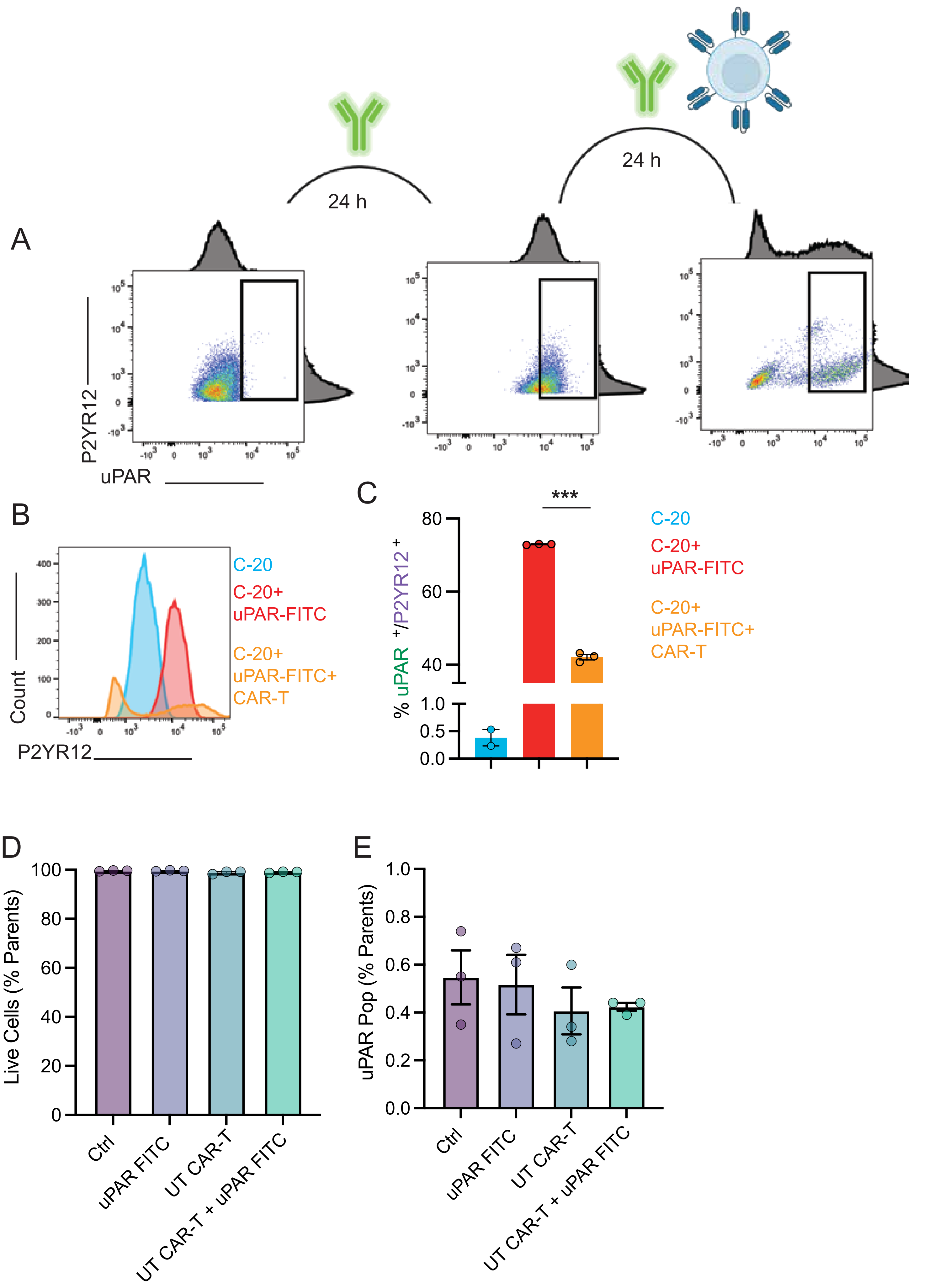
A
24 h
24 h
P2YR12
uPAR
B
Count
P2YR12
C-20
C-20+
uPAR-FITC
C-20+
uPAR-FITC+
CAR-T
C
% uPAR+/P2YR12+
***
C-20
C-20+
uPAR-FITC
C-20+
uPAR-FITC+
CAR-T
D
Live Cells (% Parents)
Ctrl
uPAR FITC
UT CAR-T
UT CAR-T + uPAR FITC
E
uPAR Pop (% Parents)
Ctrl
uPAR FITC
UT CAR-T
UT CAR-T + uPAR FITC

Fig. S14

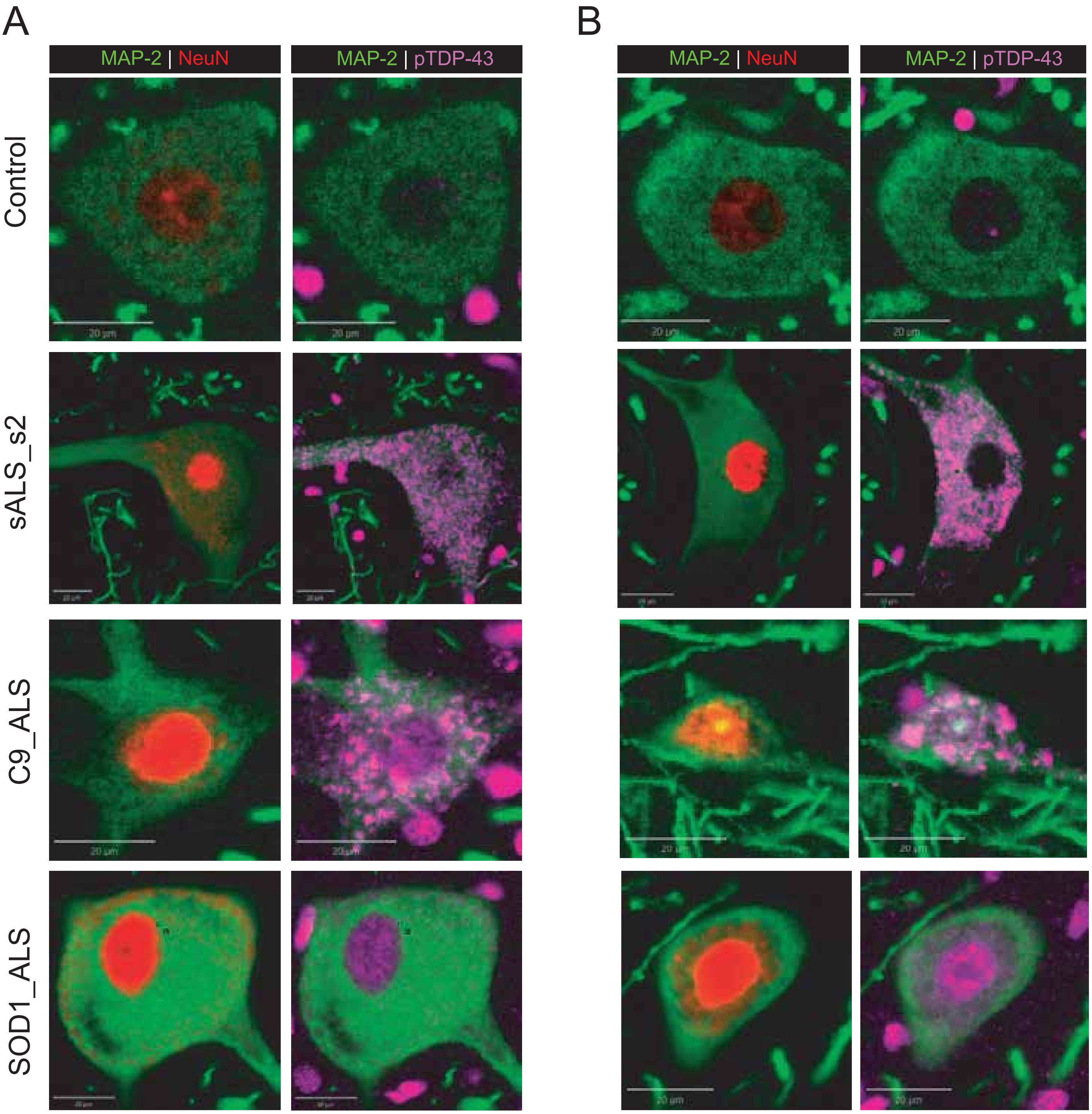


Fig. S15

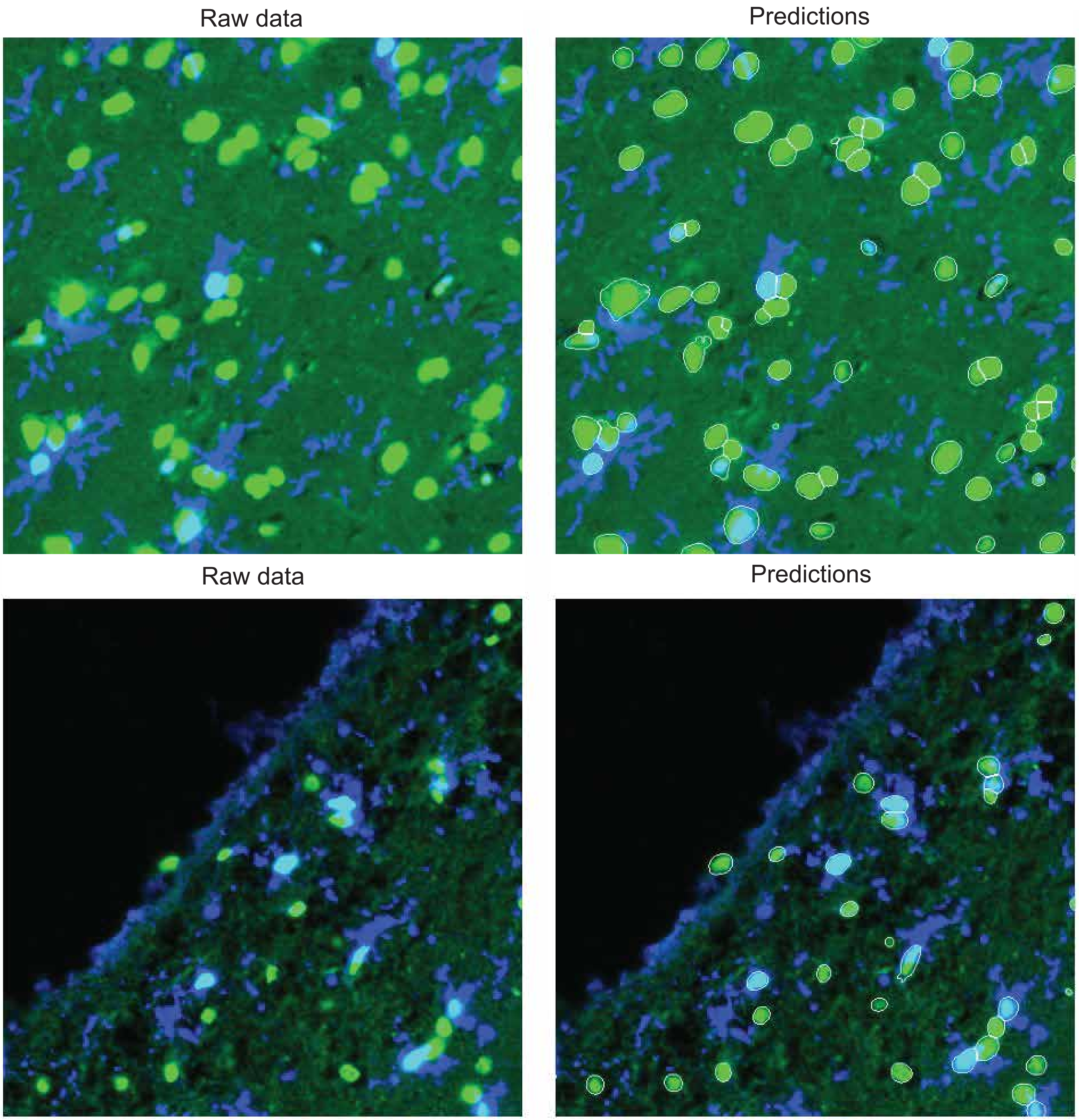


Fig. S16

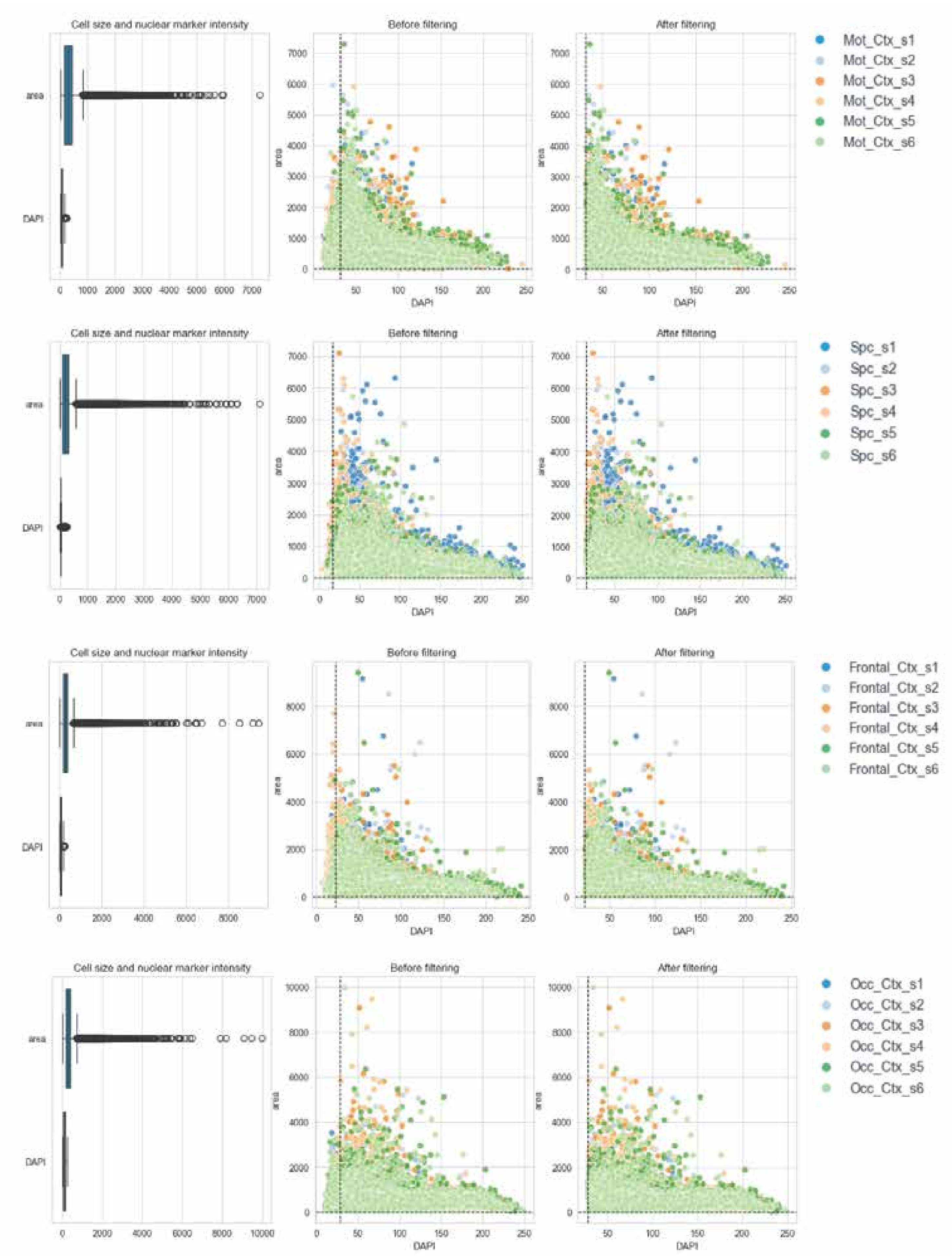

Cell size and nuclear marker intensity
Before filtering
After filtering
area
DAPI
Mot_Ctx_s1
Mot_Ctx_s2
Mot_Ctx_s3
Mot_Ctx_s4
Mot_Ctx_s5
Mot_Ctx_s6
Spc_s1
Spc_s2
Spc_s3
Spc_s4
Spc_s5
Spc_s6
Frontal_Ctx_s1
Frontal_Ctx_s2
Frontal_Ctx_s3
Frontal_Ctx_s4
Frontal_Ctx_s5
Frontal_Ctx_s6
Occ_Ctx_s1
Occ_Ctx_s2
Occ_Ctx_s3
Occ_Ctx_s4
Occ_Ctx_s5
Occ_Ctx_s6


Fig. S17

A

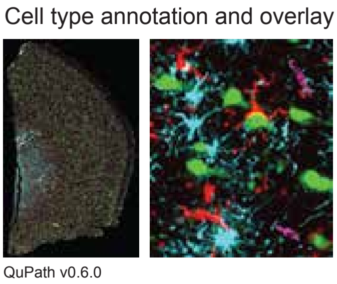
Cell type annotation and overlay
QuPath v0.6.0

B

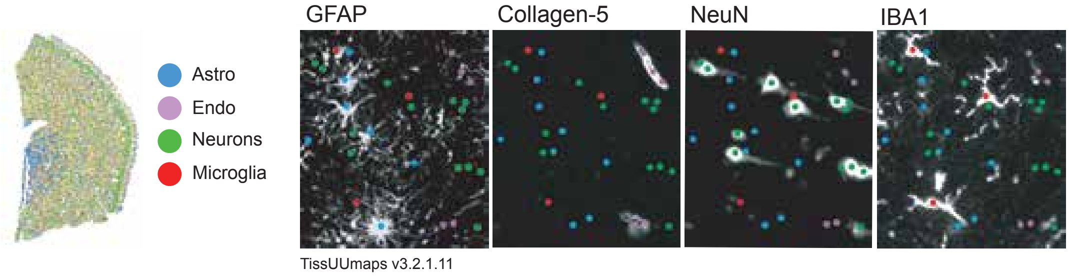
Astro
Endo
Neurons
Microglia
GFAP
Collagen-5
NeuN
IBA1
TissUUmaps v3.2.1.11

Fig. S18

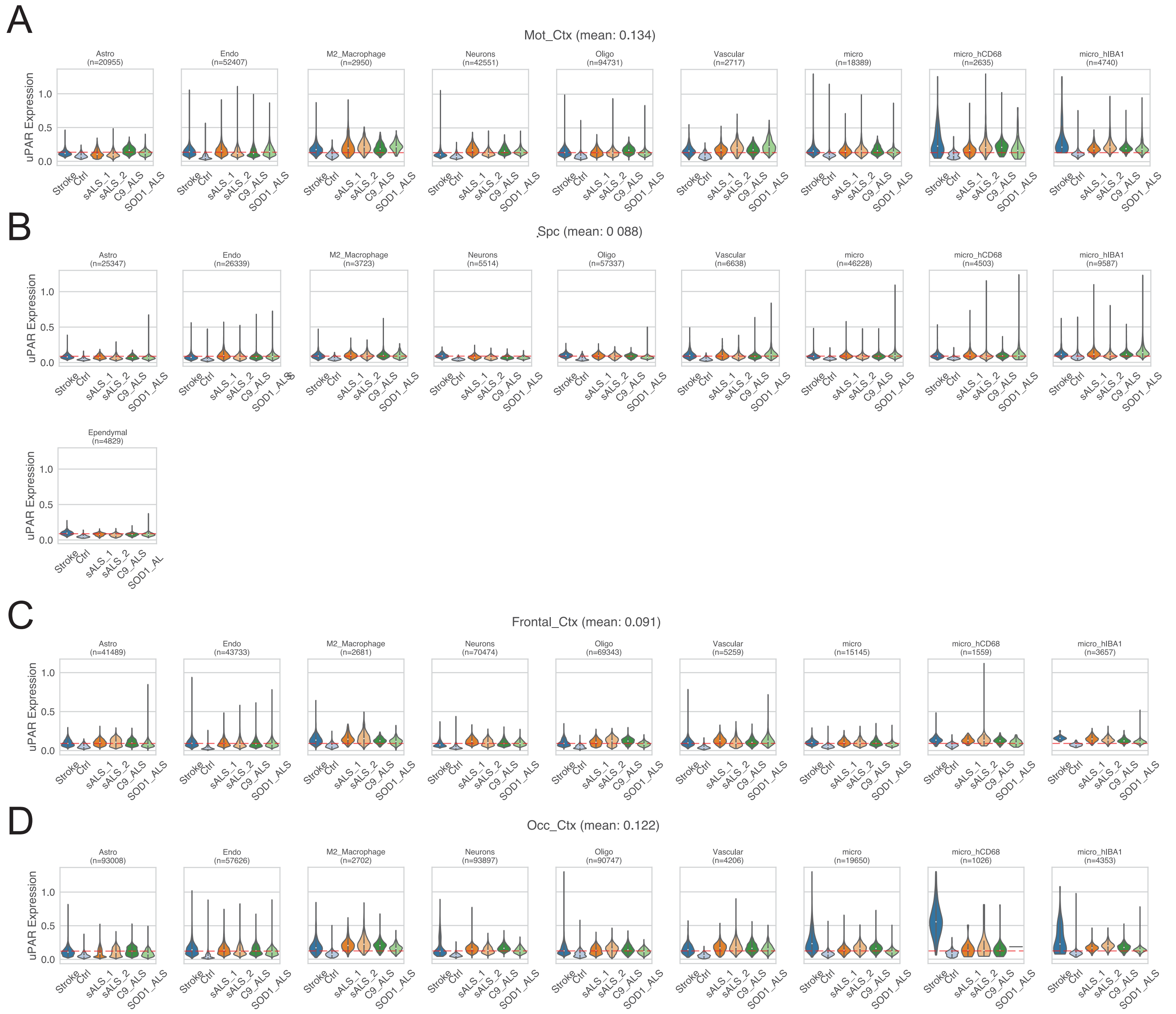
A
Mot_Ctx (mean: 0.134)
Astro (n=20955)
Endo (n=52407)
M2_Macrophage (n=2950)
Neurons (n=42551)
Oligo (n=94731)
Vascular (n=2717)
micro (n=18389)
micro_hCD68 (n=2635)
micro_hIBA1 (n=4740)
B
Spc (mean: 0 088)
Astro (n=25347)
Endo (n=26339)
M2_Macrophage (n=3723)
Neurons (n=5514)
Oligo (n=57337)
Vascular (n=6638)
micro (n=46228)
micro_hCD68 (n=4503)
micro_hIBA1 (n=9587)
Ependymal (n=4829)
C
Frontal_Ctx (mean: 0.091)
Astro (n=41489)
Endo (n=43733)
M2_Macrophage (n=2681)
Neurons (n=70474)
Oligo (n=69343)
Vascular (n=5259)
micro (n=15145)
micro_hCD68 (n=1559)
micro_hIBA1 (n=3657)
D
Occ_Ctx (mean: 0.122)
Astro (n=93008)
Endo (n=57626)
M2_Macrophage (n=2702)
Neurons (n=93897)
Oligo (n=90747)
Vascular (n=4206)
micro (n=19650)
micro_hCD68 (n=1026)
micro_hIBA1 (n=4353)
uPAR Expression
0.0
0.5
1.0
Stroke
Ctrl
sALS_1
sALS_2
C9_ALS
SOD1_ALS

Fig. S19
